# Simulation-based comprehensive study of batch effects in metabolomics studies

**DOI:** 10.1101/2019.12.16.878637

**Authors:** Miao Yu, Anna Roszkowska, Janusz Pawliszyn

## Abstract

Batch effects will influence the interpretation of metabolomics data. In order to avoid misleading results, batch effects should be corrected and normalized prior to statistical analysis. Metabolomics studies are usually performed without targeted compounds (e.g., internal standards) and it is a challenging task to validate batch effects correction methods. In addition, statistical properties of metabolomics data are quite different from genomics data (where most of the currently used batch correction methods have originated from). In this study, we firstly analyzed already published metabolomics datasets so as to summarize and discuss their statistical properties. Then, based on available datasets, we developed novel statistical properties-based *in silico* simulations of metabolomics peaks’ intensity data so as to analyze the influence of batch effects on metabolomic data with the use of currently available batch correction strategies. Overall, 252000 batch corrections on 14000 different *in silico* simulated datasets and related differential analyses were performed in order to evaluate and validate various batch correction methods. The obtained results indicate that log transformations strongly influence the performance of all investigated batch correction methods. False positive rates increased after application of batch correction methods with almost no improvement on true positive rates among the analyzed batch correction methods. Hence, in metabolomic studies it is recommended to implement preliminary experiments to simulate batch effects from real data in order to select adequate batch correction method, based on a given distribution of peaks intensity. The presented study is reproducible and related R package mzrtsim software can be found online (https://github.com/yufree/mzrtsim).

## Introduction

Metabolomic studies focus on variations among metabolites in a given system with respect to their exposure to different stimuli (Kusonmano *et al.*, 2016), such as disease (Madsen *et al.*, 2010, 2; Gonzalez-Riano *et al.*, 2016) or environmental pollutants (Bundy *et al.*, 2009; Roszkowska *et al.*, 2018). Gas chromatography coupled to mass spectrometry (GC-MS) and liquid chromatography coupled to mass spectrometry (LC-MS) are major analytical methods extensively used to reveal changes in the metabolome (Alonso *et al.*, 2015). However, in addition to the variables under study (experimental design), such as different therapies, and/or the use of different analytical methods, unwanted effects unrelated to biological variations among samples (e.g., batch effects) may also affect results in GC-MS/LC-MS based metabolomics analysis (Wehrens *et al.*, 2016). Batch effects are commonly observed in high-throughput analytical methods, which generally involve one or multiple injection sequences during instrumental analysis (Goh *et al.*, 2017; Pinto, 2017). Batch effects can stem from different factors. For instance, they can be related to known factors, such as sample injection order (Wang *et al.*, 2013) or sample amount (Wu and Li, 2016), but could also be attributed to unknown factors unrelated to experimental design. Moreover, as most metabolomics studies are performed over long periods, the obtained results could be additionally influenced by different time points of sample collection, making data analysis and interpretation more complex (Zelena *et al.*, 2009). Therefore, it is crucial to correct and normalize unwanted variables such as batch effects during the data analysis process so as to avoid irreproducible, incomplete, or misleading results.

Many research groups have been evaluating various batch effects correction strategies, but only few have provided a detailed discussion on the influence of such effects on metabolomics data (Wang *et al.*, 2013; Hughes *et al.*, 2014; Brunius *et al.*, 2016; Jr *et al.*, 2017; Ju *et al.*, 2017; Boysen *et al.*, 2018). As shown in Scheme 1A, three theoretical types of batch effects could be found in metabolomics datasets: I. monotonic (increasing or decreasing baseline), II. block (different, but constant blocks/sequences), and III. mixed (with both monotone and block changes observed simultaneously). These three types of batch effects could be corrected by a model considering such monotonic/block/mixed patterns. In high-throughput analytical methods such as GC-MS or LC-MS, batch effects could occur for different compounds or analytes at the same time. More important, such effects could be different for particular analytes due to the different physicochemical properties of the compounds under study or mass spectrometry in-source reactions (Yu *et al.*, 2017; Faber *et al.*, 2014). To overcome this problem, a commonly applied approach entails application of internal standards; however, in untargeted metabolomics studies, the availability of internal standards is narrowed down to selected classes of metabolites; as such, fluctuations in signal intensities for other classes of analytes are unavoidable. As shown in Scheme 1B, if a single compound is used to normalize the data by its response, then bias would be introduced for other peaks possessing different batch effects. Nevertheless, some batch correction methods (Sysi-Aho *et al.*, 2007; van der Kloet *et al.*, 2009) still include the application of internal standards to normalize the whole data, which consequently might overcorrect peaks with different patterns of batch effects.

To date, several different methods have been developed to correct and normalize batch effects in metabolomics data. Kohl et al. compared nine different batch correction methods in an NMR-based metabolomics study, determining that the Quantile and Cubic-Spline Normalization method showed the best performance (Kohl *et al.*, 2012). De Livera et al. reviewed seven normalized methods for removal of unwanted variations, and developed recommendations for selection of batch correction method with respect to the intended analytical application (Livera *et al.*, 2015). Using the Shiny platform, Li et al. developed an online application for sixteen normalization methods, with results identifying the Variance Stabilization Normalization (VSN), Log Transformation, and Probabilistic Quotient Normalization (PQN) methods as most adequate for data normalization in metabolomics studies (Li *et al.*, 2016). Wehrens et al. also compared several batch correction methods, concluding that when Quality Control (QC) samples and batch information were included in the models, general normalization methods also performed well (Wehrens *et al.*, 2016). Overall, the appropriateness of a batch correction method for a given dataset is greatly dependent on whether the statistical model and assumption behind the correction method fit the real data, and as such, selection of an appropriate method should take these considerations into account.

The correction of batch effects in untargeted metabolomics is also challenging with respect to validation of batch correction methods. In order to evaluate the performance of selected correction methods, Li et.al developed the online tool NOREVA to make comparisons among twenty-five correction methods, taking into account five criteria: intra-group variation among samples, the distribution of P-values from differential analysis, consistency of certain markers among different datasets, classification accuracy, and correspondence of reference data (Li *et al.*, 2017). However, except for the last criteria, which actually uses the same routine as that used in targeted analysis, the proposed criteria scantily and no instinctively allow for selection of the most adequate correction method. Besides, each validation method was only based on ideal statistical properties or distributions, which might introduce bias when the intended data has a different distribution or structure.

Moreover, if the batch effects for each peak are known, then validation and comparisons of different batch correction methods would be possible and easy. In this situation, direct counting of peaks with real changes could be performed before and after the data were corrected. However, such correction method cannot be applied when the research is designed to discover novel compounds (biomarkers) or to explore heterogeneity within the samples. In such cases, *in silico* simulations of such data would be a convenient tool to solve these issues. Simulation of metabolomics data has been proposed in multiple studies (Mendes *et al.*, 2005; Parsons *et al.*, 2009; van der Kloet *et al.*, 2009; Jr *et al.*, 2017; Reisetter *et al.*, 2017). However, similarly to batch correction methods, previous simulation methods in metabolomics were developed from data stemming from genomics microarray studies, and as such, did not consider the statistical properties of MS-based metabolomics data, where multiple peaks from the same compound, such as adducts, isotopologues, and fragment ions, would show high correlations. In this case, metabolomics data have a hierarchical structure with correlations at both the peak and compound level (Mahieu and Patti, 2017; Yu *et al.*, 2019). In the case of metabolomics data, the hierarchical structure indicates that all the peaks from the same compound are correlated with each other, and that a given compound might also be correlated with other compounds. Hence, peak level data also contains compound level correlations. Therefore, variances would be biased on compounds with more peaks.

In this study, as a starting point, the statistical properties of GC-MS/LC-MS based data available as already published datasets are discussed in order to elucidate what considerations should be taken into account for *in silico* simulations. Next, we carried out *in silico* simulations of available metabolomic datasets in order to evaluate various batch correction methods under different scenarios. In addition, the whole process developed as part of this study has been made reproducible and transparent by releasing all related functions and data processing code.

## Materials and methods

### Datasets

MTBLS28 (Mathé *et al.*, 2014), MTBLS59 (Franceschi *et al.*, 2012), MTBLS341 (Strehmel *et al.*, 2016), MTBLS351(Pedersen *et al.*, 2016), MTBLS393 (Muhamadali *et al.*, 2016) and the faahKO data package (Saghatelian *et al.*, 2004) were used to attain the statistical properties used in the *in silico* simulations. In total, 14 related datasets were used to simulate 7 different scenarios. These datasets are summarized in Table 1.

**Table 1.**
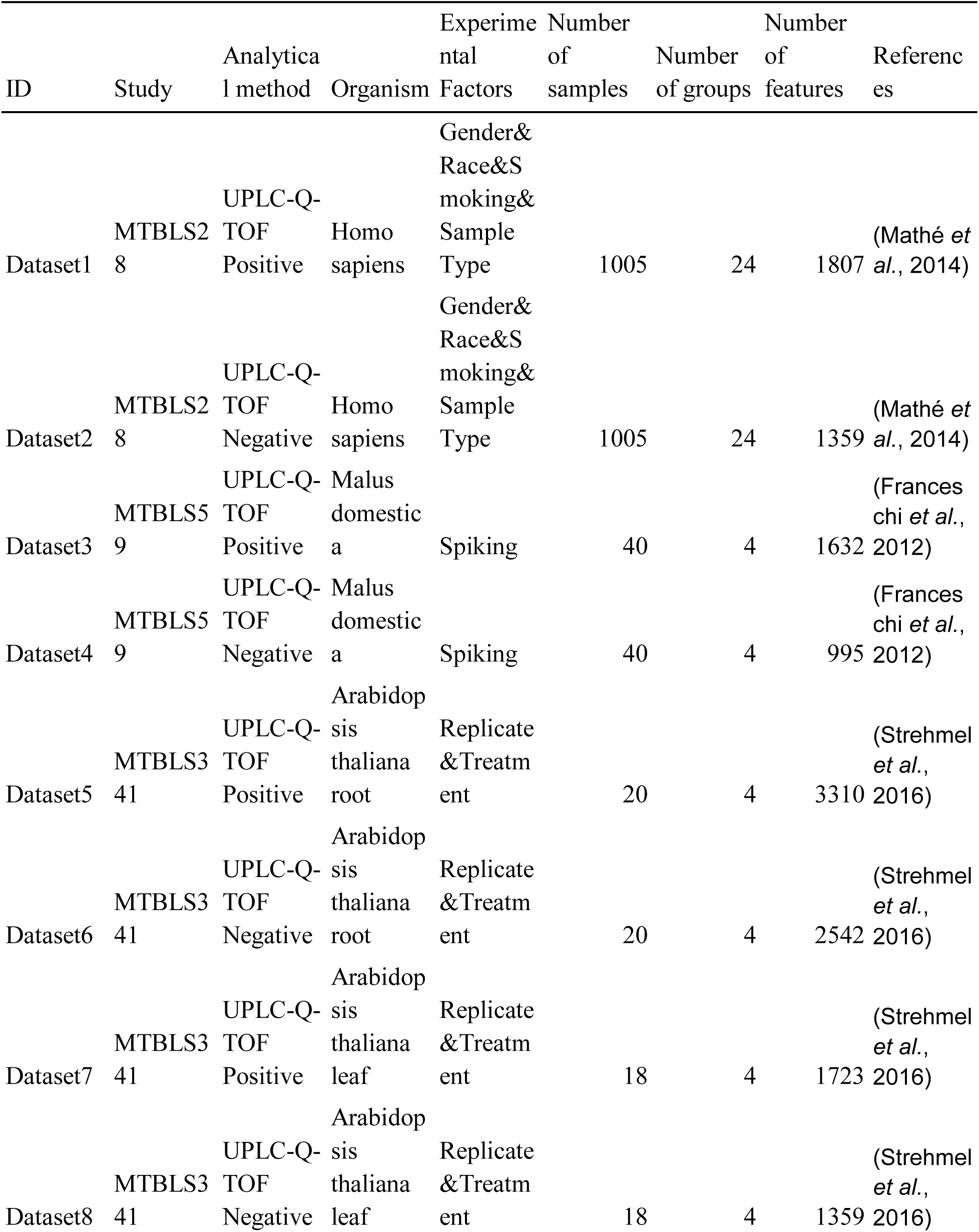
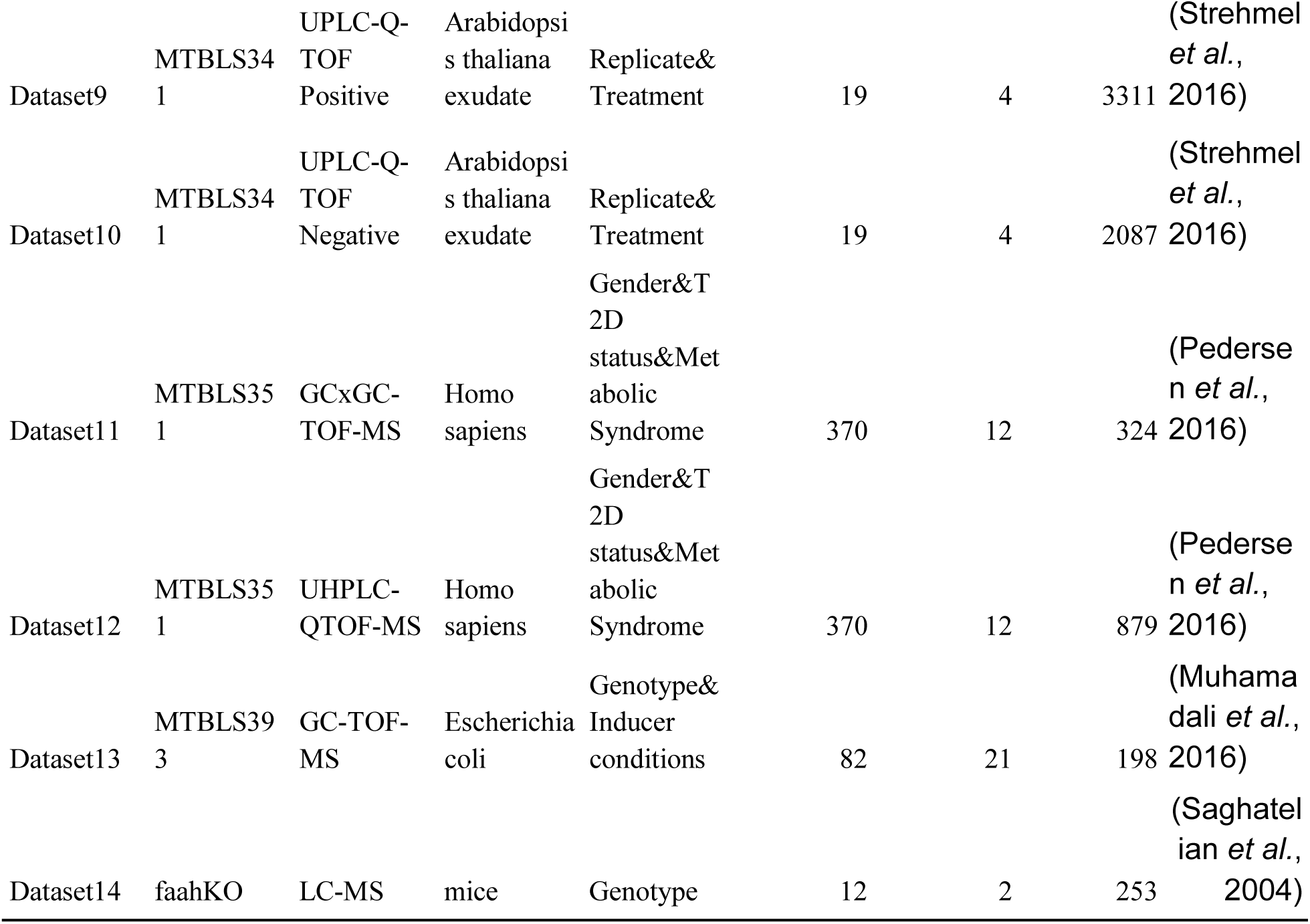
Summary of the datasets used in this study.

### Application of datasets for in silico simulation

In untargeted metabolomics, GC-MS/LC-MS based data is represented by peaks occurring across multiple samples. Statistical analysis of metabolomics data is thus performed based on the experimental design. Given this, *in silico* simulations of such data should thus be commenced with peaks attained from a single sample. The simulation could then be extended to multiple samples based on the information contained in the real datasets.

Therefore, taking into account the experimental design for the selected datasets (Table 1), the first step of this study encompassed calculations of average peak intensities across single samples within particular groups. As presented in Figure 1, the log transformed intensity distribution of detected peaks showed a left-skew pattern for Dataset 1, which contained 24 groups. Various patterns were observed for the other real datasets (Datasets 2-14); however, none of these distributions could be simply expressed as normal distributions (see Figures S1-S13). In order to simulate such peaks’ intensities *in silico*, a Weibull distribution was applied to the data, since such distribution provides the possibility for right- or left-skewed patterns (as observed in real datasets), depending on the corresponding parameters, such as shape and scale. In LC-MS analysis, each compound can generate multiple ions or peaks, such as isotopologues, adducts, and neutral loss, on mass spectrometry (Yu *et al.*, 2019). In GC-MS, specifically in EI mode, each compound would generate multiple fragments along with molecular ions (Yu *et al.*, 2017). In this study, the percentages of the detected compounds, expressed as the ratio between the number of compounds and their corresponding peaks, were defined for real datasets and then included into the *in silico* simulation. When such a parameter is lower than 50%, more than two random peaks could come from a single compound. Moreover, according to previously reported work, 5-20% of peaks could indicate either major variances or real compounds (Mahieu and Patti, 2017; Yu *et al.*, 2019); as such, this assumption was taken into account during our simulation studies. We also assumed that correlated peaks from the same compound would not change the Weibull distribution for other peaks.

**Figure 1.**
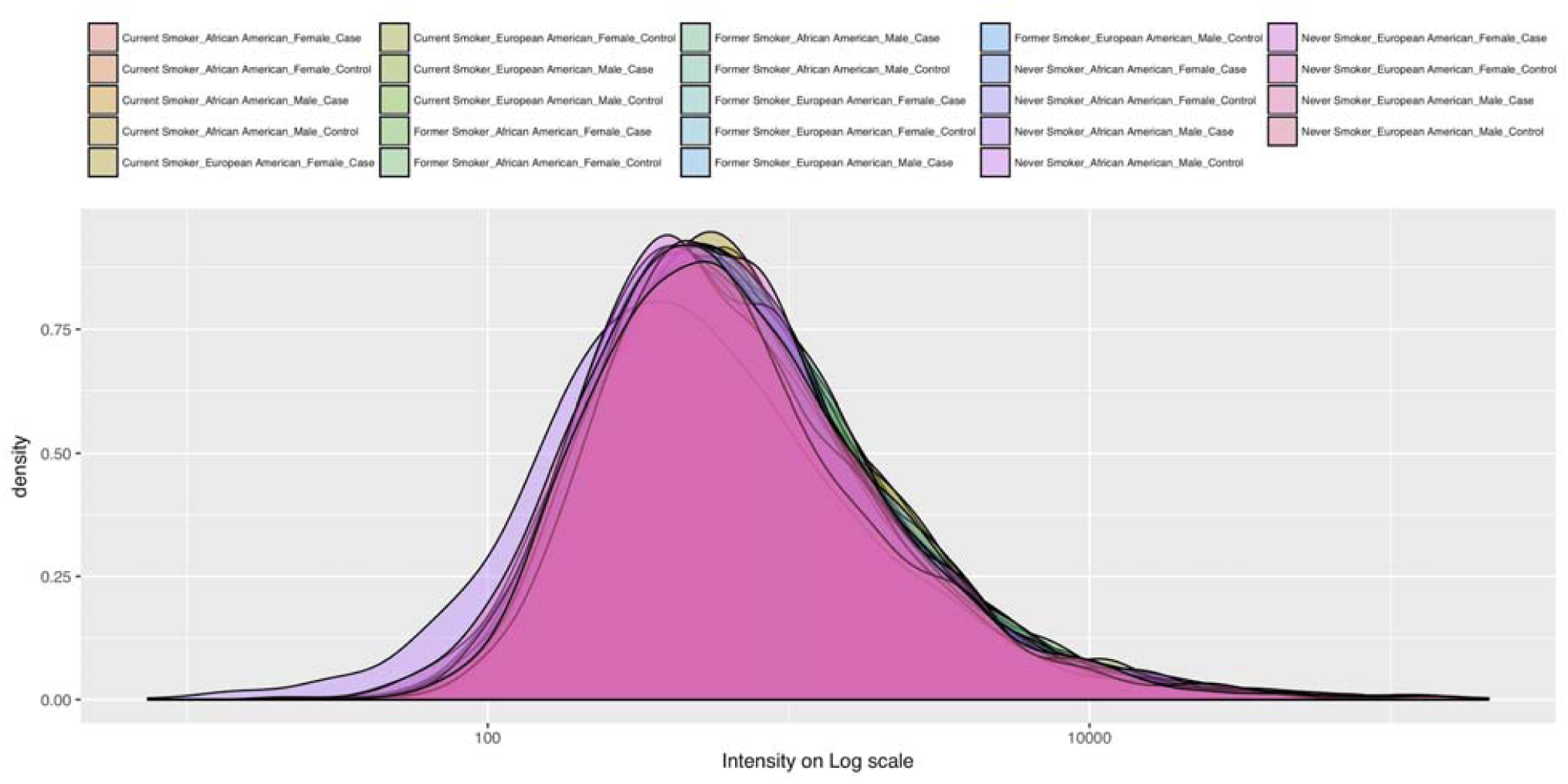
Distribution of real peaks’ intensities on the Log scale for Dataset 1.

In a simulation of the peaks’ intensity, only a few peaks would be generated at first according to the Weibull distribution. For example, for a simulation of 1000 peaks (which are obtained from 100 compounds), only 100 “molecular ions” peaks would be generated at first according to the Weibull distribution. The other 900 peaks being simulated would account for adducts, neural loss, or fragmental ions based on resampling of those 100 “molecular ions” peaks, and would be weighted by a factor calculated from an exponential of normal distributed folds with mean 0 and standard deviation 1. Since peaks from the same compound would be correlated with one another, such a simulation should thus reflect the real ionization and/or fragmental process of compounds occurring during mass spectrometry analysis.

As a next step of the presented work, %RSDs were calculated for samples in particular groups or for biological replicates in each real dataset. Based on the obtained results for real datasets (Figure 2 and Figure s14-s26), the applied %RSDs distribution in *in silico* simulation could also be a Weibull distribution.

**Figure 2.**
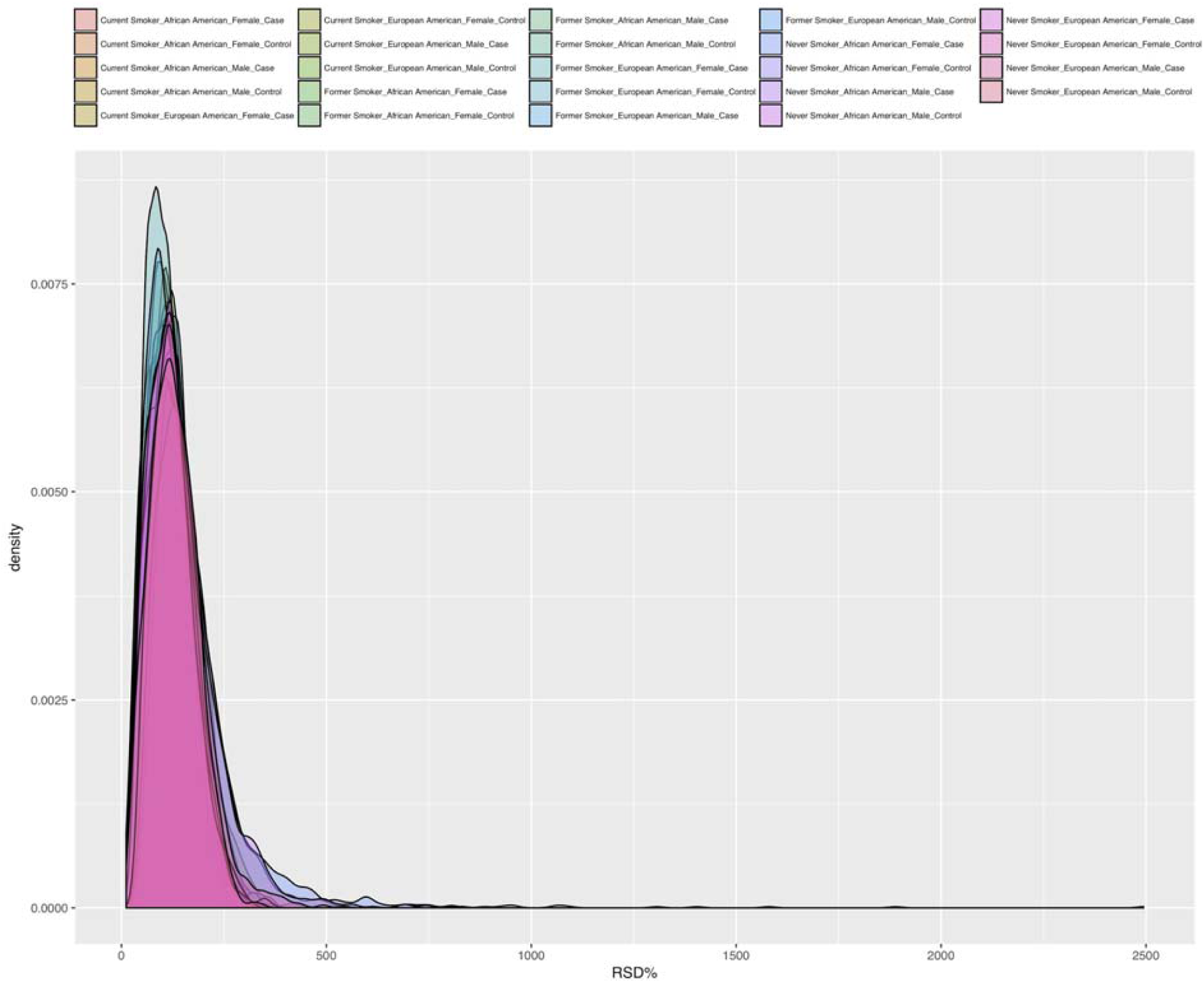
The RSD% distribution of real peaks’ intensities on Log scale within 24 groups of Dataset 1.

After performing a simulation on single samples and then on biological replicates in one group, the fold changes among different groups could be simulated. As presented in Figure 3 and Figures S27-S39, most of the analyzed real datasets’ fold changes among multiple samples in different groups followed an exponential distribution of normal distributed random errors. In this case, most peaks showed no changes among groups; however, few peaks presented large differences among groups. Since it is impossible to predict biological differences among groups, random peaks were selected to calculate a known fold change of intensity.

**Figure 3.**
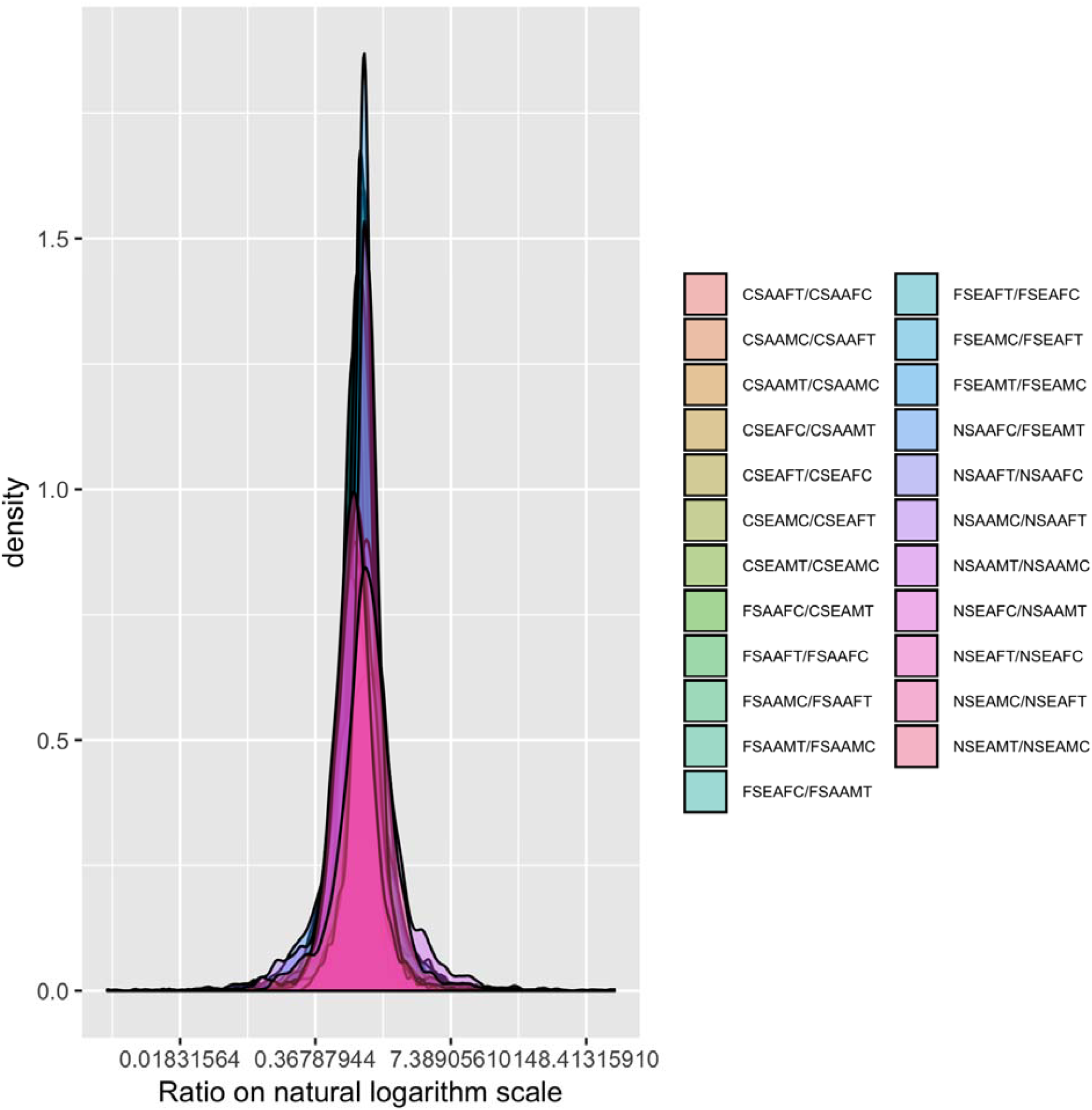
Distributions of ratios of group averages for Dataset 1.

Based on the results from this analysis of multiple samples in real biological datasets, an *in silico* simulation study was carried out to generate 1000 peaks from two created biological groups (case and control), each containing 10 samples. The selected peaks from each real group showed different average intensities (according to the results obtained from the analysis of real samples), where peaks were influenced by different biological conditions (experimental design). Next, for the batch effects analysis, three previously mentioned types of batch effects, namely monotone, block, and mixed effects, were simulated *in silico*. As mentioned, for monotone batch effects, the changed folds typically undergo an increasing/decreasing trend during the injection sequences. For block batch effects, the changes would also follow an exponential distribution of normal distributed random errors. To simulate mixed batch effects, both monotone changes and block changes were considered.

In the simulations performed in this study, block batch effects affected 8, 5, and 7 samples, whereas monotone batch effects were distributed across all samples. Influenced peaks from each block batch showed different average intensities across multiple samples to simulate unwanted batch effects. We considered that all peaks from the same compound were influenced by the experimental design. However, batch effects were randomly assigned at the peak level since such effects might also appear after the in-source ionization process. For peaks from a single sample, the shape and scale for the Weibull distribution were simulated as 2 and 3. The parameters of the Weibull distribution were selected based on the range of parameters observed in real datasets. For the relative standard deviation (%RSD), the shape and scale were set as 1 and 0.18, respectively. Given the nature of such effects, analyses at the peak level, instead of the compound level, were performed in evaluation studies of batch effect correction methods, which are discussed later in this paper.

Next, based on the statistical probability of the available 14 real datasets, we simulated and evaluated seven scenarios that are common in real research datasets (Table 2) in order to evaluate and compare the performance of different batch effects correction strategies already reported in the literature. Scenario 1 is the most common scenario applied for untargeted analysis, with 100 compounds generating 1000 peaks, among which 100 peaks are influenced by experimental design and 100 peaks are influenced by mixed batch effects. In scenario 2, simulated compounds are almost independent of each other, a scenario similar to those observed in targeted analysis approaches. Scenario 3 takes into account peak profiles, where half of the peaks are influenced by the experimental design. This scenario is observed when the phenotype shows obvious changes and untargeted analysis is performed to verify alterations at the metabolite level. Scenario 4 reflects an experiment where half of the peaks are influenced by mixed batch effects. Scenario 5 combines scenario 3 and scenario 4. Scenario 6 and scenario 7 concern the identification of differences between the two distinct types of batch effects: monotone and block. Overall, each of the above-mentioned scenarios was simulated *in silico* 1000 times. Simulation was performed on raw data and also on log transformed data for each scenario. In total, 252000 batch-corrected data were obtained on 14000 different simulated datasets to provide stable and reliable results for further discussion.

**Table 2.** Parameters for different simulation scenarios

|  | compounds# | changed peaks | Batch peaks | Batch type |
| --- | --- | --- | --- | --- |
| Scenario 1 | 100 | 100 | 100 | mixed |
| Scenario 2 | 900 | 100 | 100 | mixed |
| Scenario 3 | 100 | 500 | 100 | mixed |
| Scenario 4 | 100 | 100 | 500 | mixed |
| Scenario 5 | 100 | 500 | 500 | mixed |
| Scenario 6 | 100 | 100 | 100 | monotone |
| Scenario 7 | 100 | 100 | 100 | block |

### Batch correction methods for simulated scenarios

Finally, 17 batch correction methods, including autoscaling (van den Berg *et al.*, 2006), pareto scaling (van den Berg *et al.*, 2006), range scaling (van den Berg *et al.*, 2006), vast scaling (van den Berg *et al.*, 2006), level scaling (van den Berg *et al.*, 2006), total sum (De Livera *et al.*, 2012), median (De Livera *et al.*, 2012), mean (De Livera *et al.*, 2012), Probabilistic Quotient Normalization (PQN) (Dieterle *et al.*, 2006), Variance Stabilization Normalization (VSN) (Kohl *et al.*, 2012), Quantile (Kohl *et al.*, 2012), the robust spline normalization (RSN) from the lumi package (Lin *et al.*, 2008), cyclic loss from the Limma package (Ballman *et al.*, 2004), CUBIC Spline from the affy package (Workman *et al.*, 2002), Surrogate Variable Analysis (SVA) (Leek and Storey, 2007, 2008), independent surrogate variable analysis (ISVA) (Teschendorff *et al.*, 2011), and Principal component regression (PCR) (Yang *et al.*, 2008), in addition to a no correction mode (where an uncorrected dataset was used as baseline), were selected in this study for a comparison of their performance in 7 simulated scenarios based on statistical properties from real datasets. The first 14 methods are general normalization methods, while the former 3 methods are batch correction methods based on linear models that assume latent batch effects could be treated as one variable in a linear model. In normalization methods, usually row-wise or column-wise adjustments are made without taking into account experimental design, whereas in linear model based batch corrections, potential batch variables are considered and included in the linear model to estimate the batch effects.

Statistical analysis was carried out in order to compare and contrast the 18 selected batch correction methods (17 methods and a no-method control), using computations of the true positive rates and false positive rates of 1000 repeated simulations of each of the 7 scenarios used in this study, with the false discovery rate controlled by Benjamini and Hochberg (BH) adjusted p-values. The cutoff of the BH adjusted p-value was set at 0.05, as typically used in metabolomics differential analysis studies. The true positive rate is the ratio of the number of real changed peaks found in real datasets (true positive) and the number of simulated changed peaks. The false positive rate is the ratio between the number of false positive peaks and negative peaks. The simulation process and batch correction algorithm were run using the mzrtsim package and script, which are listed in the supporting information.

## Results and discussion

In total, 18 selected batch correction strategies were applied for each of the 7 scenarios, and their data compared and contrasted. The results of this evaluation study are discussed for each scenario in the next section.

### Comparison 1: Dependent peaks and Independent peaks

In the first comparison, 18 batch correction methods were applied for Scenario 1 (dependent peaks) and Scenario 2 (independent peaks), where 100 simulated peaks out of 1000 peaks were real biological changes, and 100 simulated peaks could be influenced by the batch effects. The influences of existing relationships among peaks on batch effect correction were investigated and compared for each scenario. A comparison between these two scenarios was carried out to investigate whether the applied batch correction methods were influenced by the peaks’ compound dependence.

As presented in Figure 4, when batch correction was performed on raw data, the 14 general batch correction methods (autoscaling, pareto scaling, range scaling, vast scaling, level scaling, total sum, median normalization, mean normalization, PQN, VSN, quantile, robust spline, cyclic loess, and CUBIC spline) showed similar true positive rates compared to data without correction (no correction), indicating that any of these 14 general methods could be effective when applied to cases similar to scenario 1 and scenario 2. On the other hand, the three linear models used in this study (SVA, ISVA, and PCR) were found to increase the false discovery rate, which is not a desired effect. However, once the data was log transformed, batch correction methods yielded a better performance as compared to that observed for raw data (Figure 4). When log-transformed data was used instead, a comparison of batch correction methods’ performances between scenarios 1 and 2 revealed that the linear model-based correction methods provided larger false positive rates in dependent data (scenario observed in untargeted metabolomics study) in comparison to independent data. Therefore, the results suggest that in cases where metabolomics data from GC-MS/LC-MS based studies possess multiple peaks from the same compounds, such methods’ false positive rate would be worse if the data were log transformed prior to correction as opposed to use in its raw format.

**Figure 4.**
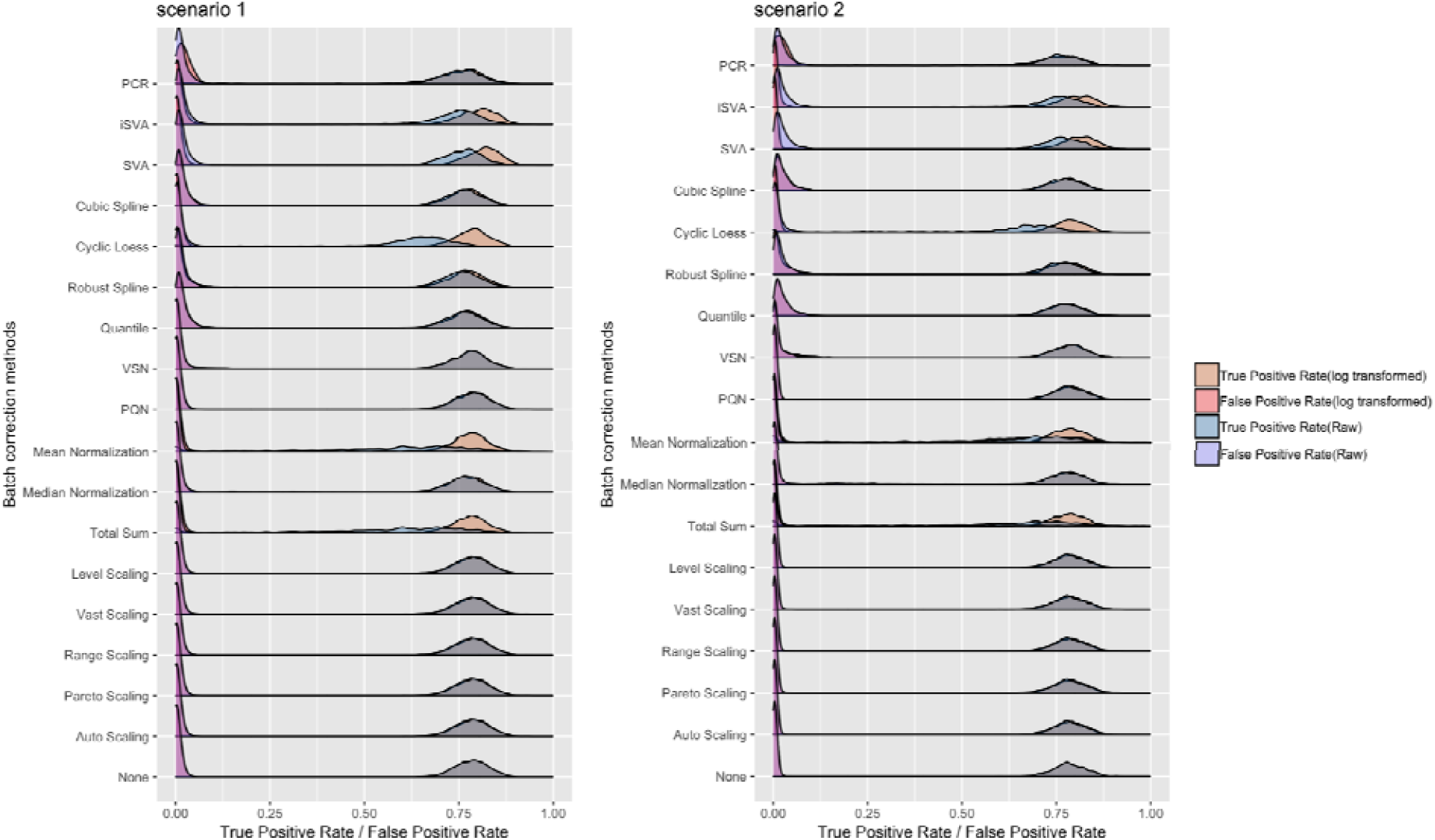
Ridges plot of true positive rates and false positive rates of 1000 times simulated data when the BH adjusted p-value cutoff is 0.05 for scenario 1 and scenario 2, and their log transformed data.

### Comparison 2: experimental design dominated and batch effects dominated scenarios

In the second comparison, we set100 compounds to generate 1000 peaks. In total, 3 unique datasets corresponding to 3 different scenarios were simulated for comparison. First, a dataset with 500 and 100 peaks changed by experimental design and batch effects, respectively, was simulated (scenario 3). This simulation took into account a scenario in which the resulting dataset characteristics would be primarily owed to the experimental design, and where most of the peaks could be used to separate biological groups. The second scenario considered in this comparison entailed a simulation with 100 and 500 peaks that were changed by experiment design and batch effects, respectively (scenario 4). Finally, a third scenario (scenario 5), combining properties of scenario 3 and scenario 4, and having 500 and 500 peaks changed by experiment design and batch effects, respectively, was included in this comparison. These three scenarios are commonly observed in real metabolomics datasets.

In this comparison, “no correction” data was treated as a baseline for each scenario. As presented in Figure 5, application of 17 batch correction methods on raw data revealed that scenarios 4 and 5 yielded much lower true positive rates and false positive rates (around 50% and 60%), respectively, when compared to treatment dominated data (around 80%) in scenario 3. However, most of the batch correction methods used in these three scenarios made no improvements in the differential analysis, especially when applied to the raw data. Only linear model-based correction methods, such as SVA, yielded better performance on true positive rates for batch effects dominated data (scenario 4). However, the tradeoff of using such methods is incurring an increase in false positive rates compared with other batch correction methods. Interestingly, log-transformed data improved most of the correction methods’ performance on true positive rates. In cases where studies are designed to yield high true positive rates, log transformation would be useful prior to batch correction.

**Figure 5.**
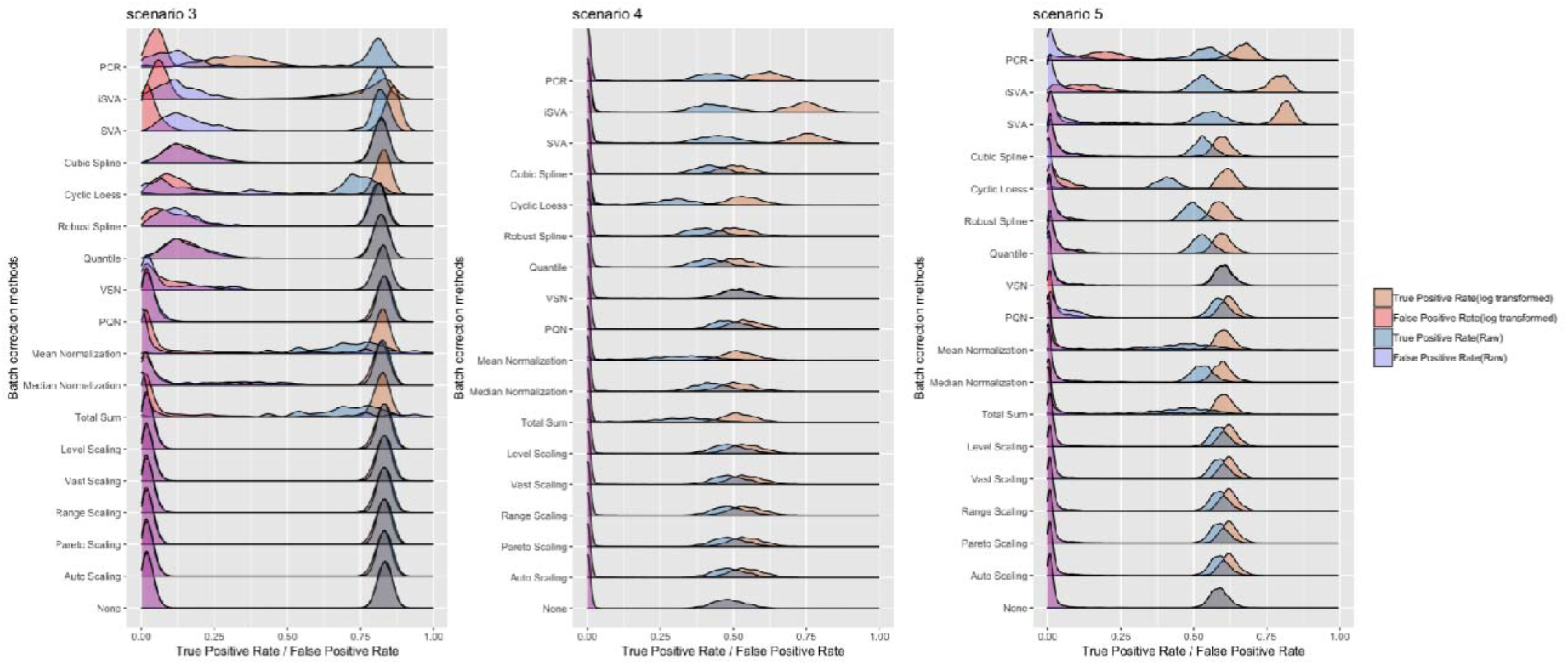
Ridges plot of true positive rates and false positive rates of 1000-times simulated data. The BH adjusted p-value cutoff was set at 0.05 for scenario 3, 4 and 5, and their log transformed data.

### Comparison 3: influence of different types of batch effects

The applied batch effects in the above-mentioned simulations were of mixed (monotone and block) type only. However, in real datasets, observed batch effects may be mixed mode, monotone, or block. Therefore, in the third comparison, the mixed batch effects were separated into two different, not previously analyzed scenarios: one where the dataset was only affected by monotone batch effects (scenario 6), and another one where only blocked batch effects affected the dataset (scenario 7). The performance of different batch correction methods on these two new scenarios was then compared. All other parameters were the same as in scenario 1.

The obtained results indicate that the performances of all batch correction methods used in both scenarios were similar; however, most of the analyzed methods had better performance towards monotone batch effects correction (Figure 6). Nevertheless, log transformation of the data still had a strong influence on linear model-based batch correction methods, especially on false positive rates (see Figure 6). Therefore, the results presented in this comparison point to a pressing need for the development of new batch correction methods capable of addressing specific types of batch effects. Particularly in cases where the applied correction method is based on a linear model (such as the ones observed in scenarios 6 and 7), careful analysis of false positive rates should be carried out on the resulting data.

**Figure 6.**
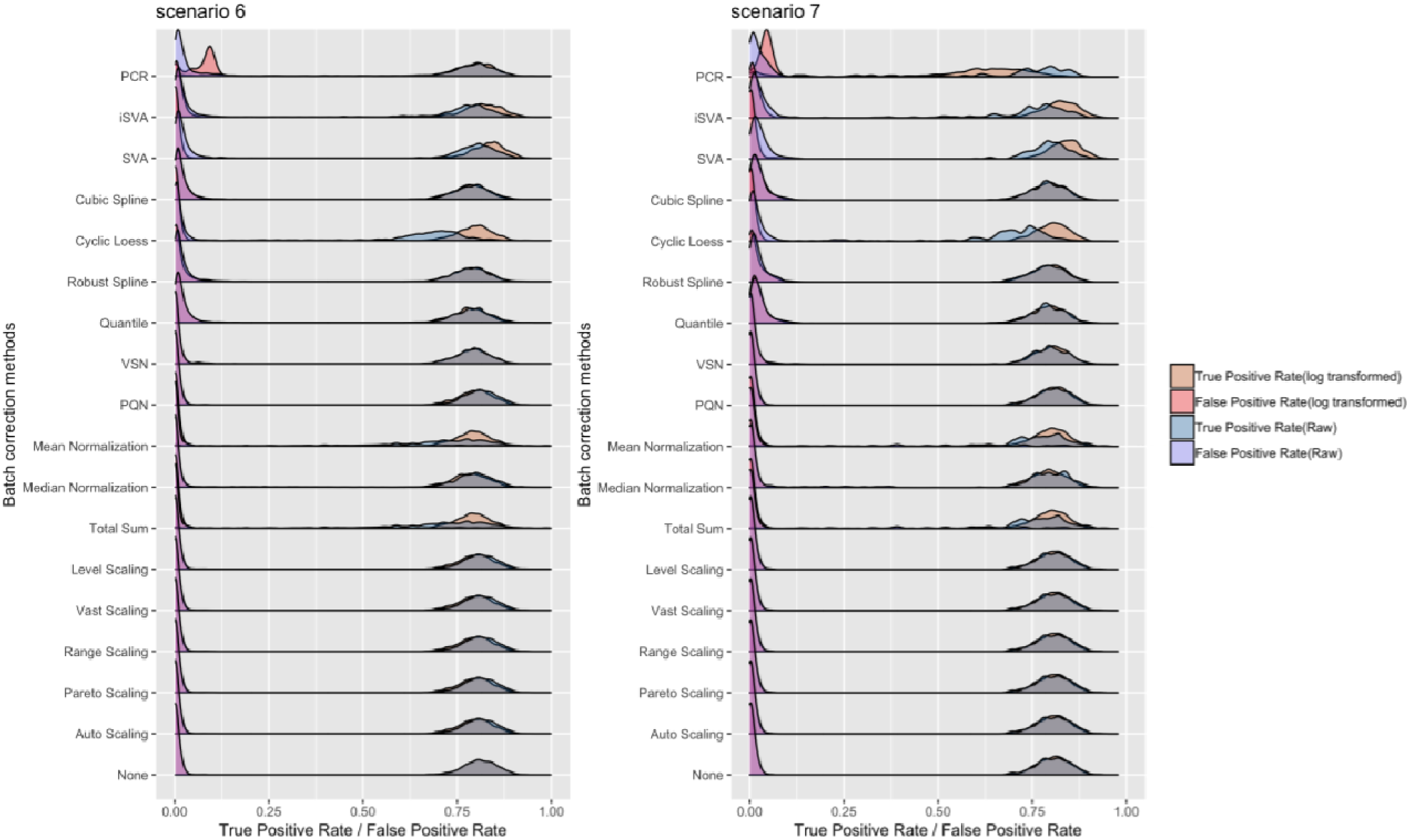
Ridges plot of true positive rates and false positive rates of 1000 times simulated data when the BH adjusted p value cutoff is 0.05 for scenarios 6 and 7, and their log transformed data.

## Implication

The results of the current study, which encompassed 252000 batch corrections on 14000 different simulated datasets, with subsequent differential analysis of the 18 available batch correction strategies on the generated data, show that none of the applied methods were fully useful for each scenario presented in this study, as all included batch correction methods were strongly influenced by the correction models and statistical properties of the data. Therefore, preliminary experiments should be implemented to simulate batch effects from real data. In this aspect, the proposed mzrtsim package could be used to simulate metabolomics data based on theoretical statistical distributions or on statistical distributions of real data. The obtained information could then be used to inform decisions regarding method selection, based on a given distribution of peaks intensity. However, since each peak would possess different batch effects, the application of pooled QC sampling would be a useful tool to control unwanted influences, ascertaining that only peaks with lower RSD% in all pool QC samples are taken into account for further analysis (Dunn *et al.*, 2012). Although the development of novel batch correction methods would be useful, particularly methods capable of addressing specific types of batch effects, methods that ascertain whether it is worthwhile to apply batch corrections might be more meaningful for researchers. However, not all scenarios typically encountered in analytical studies were considered in this work. In the presented scenarios, typical of untargeted metabolomics, the analysis carried out as part of this work has been made reproducible as a script and could be easily modified with different statistical properties or used directly to perform simulations from real datasets.

## Conclusion

Statistical properties and multipeak-based simulations reveal potential issues during data analysis, such as skewed distribution, different types of batch effects, and inner associations within data. Log transformation of datasets might be required as a preprocessing step in order to achieve better results. Corrections based on certain types of samples or peak(s) could cause overcorrection of other peaks. Further, traditional correction methods might increase the false positive rates of the datasets while not yielding much improvement with respect to true positive rates. While SVA, a linear model-based correction method, could be selected as a general batch correction method under different conditions after log transformation of the data, the method nonetheless still poses a risk of increasing false positive rates.

## Acknowledgments

This research was financially supported by Industrial Research Chair of the National Sciences and Engineering Research Council of Canada (NSERC-IRC).

---

**Scheme 1.**
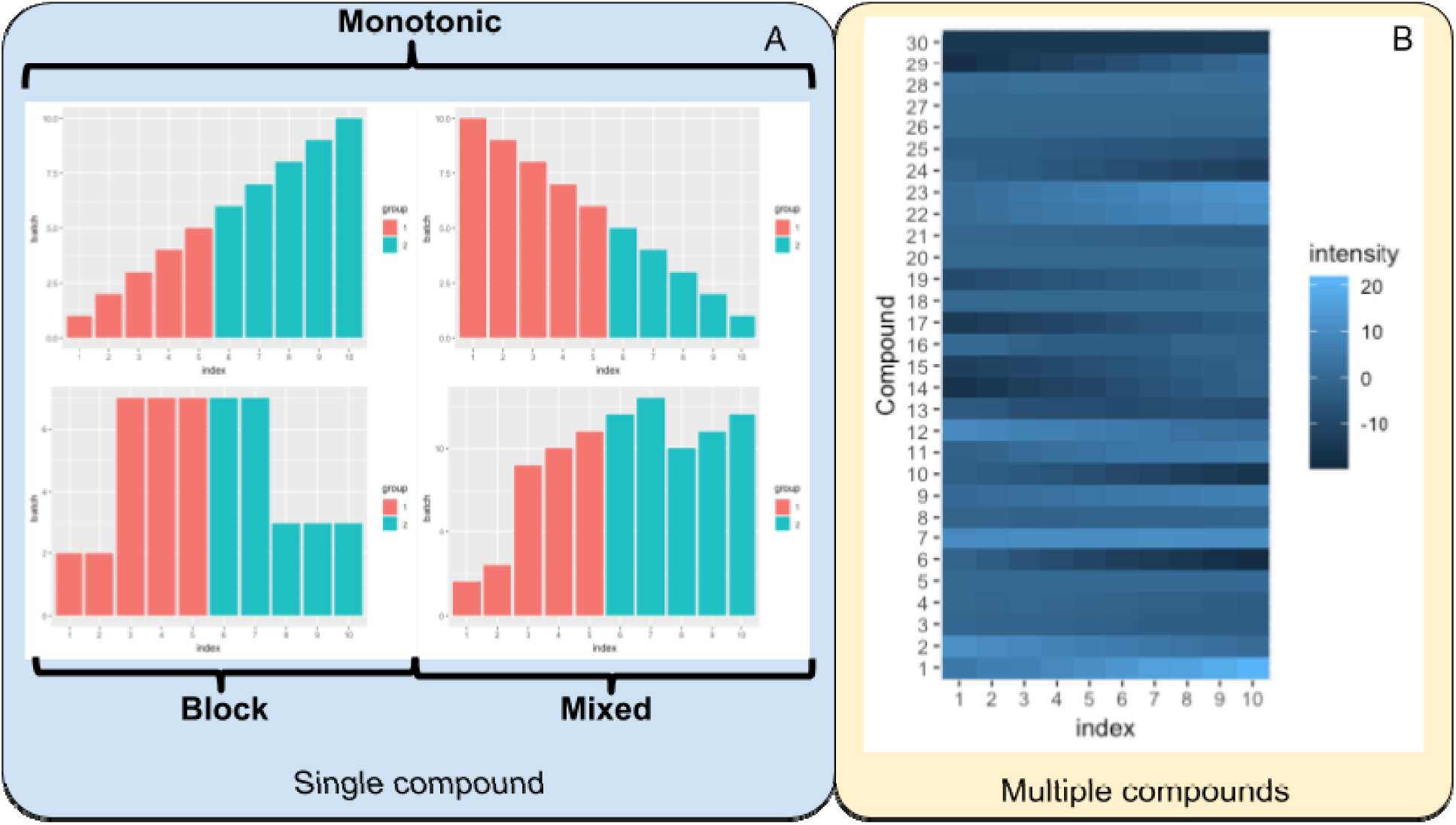
Batch effects in metabolomics studies. (A) Three types of batch effects occurring on a single compound. (B) Demonstration of batch effects on multiple compounds with different types of batch effects observed in high-throughput analysis.

