## Supplemental Table and Figure for "Simulation-based comprehensive study of batch effects in metabolomics studies"

### ^1^Department of Chemistry, University of Waterloo, 200 University Avenue West, Waterloo, Ontario, N2L 3G1, Canada

^2^Department of Pharmaceutical Chemistry, Medical University of Gdańsk, Gdańsk, Poland

^+^current address: Department of Environmental Medicine and Public Health, Icahn School of Medicine at Mount Sinai, New York, USA

**Keywords:** Metabolomics, batch effects, simulation

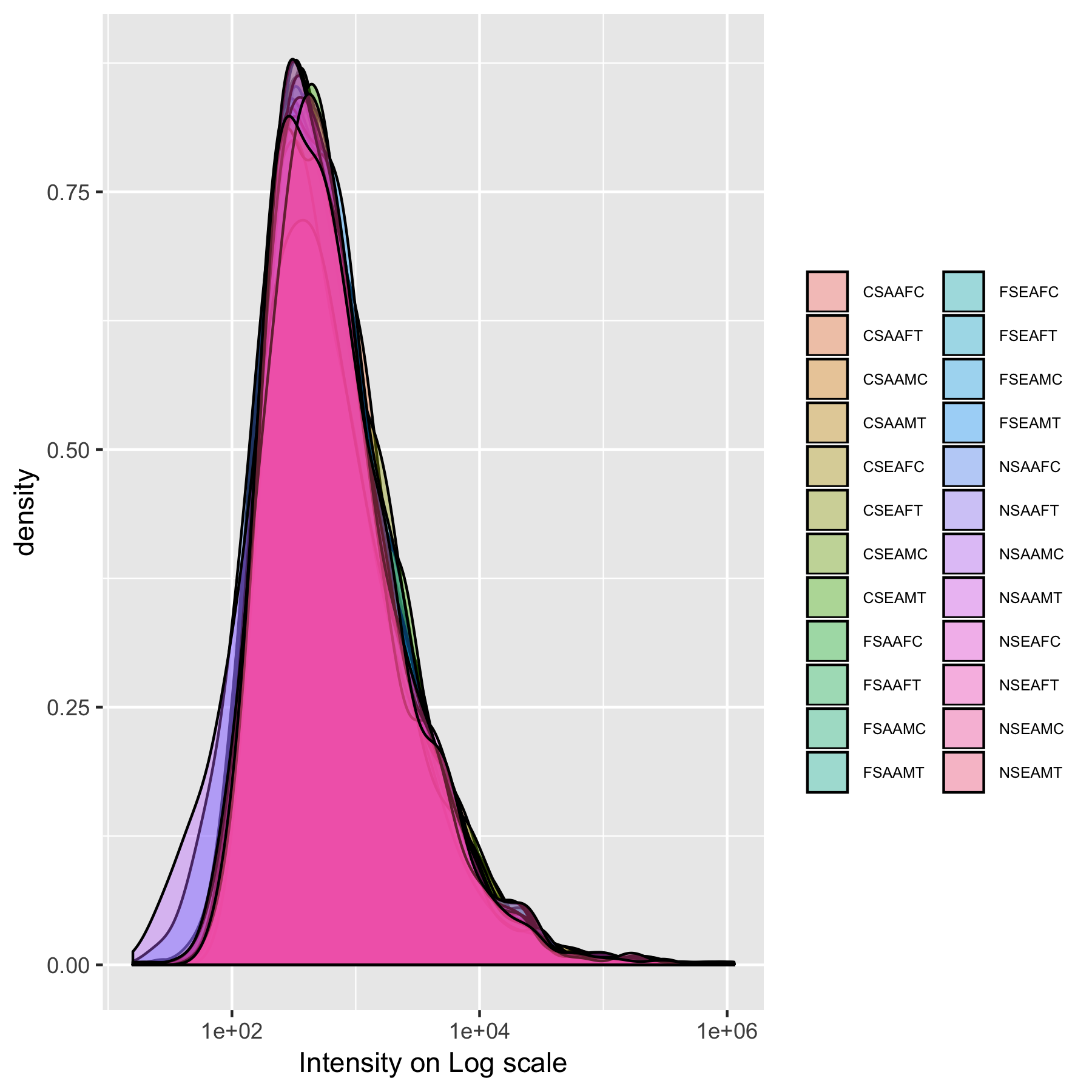

Figure S1. Distribution of peaks’ intensities on Log scale from Dataset 2.

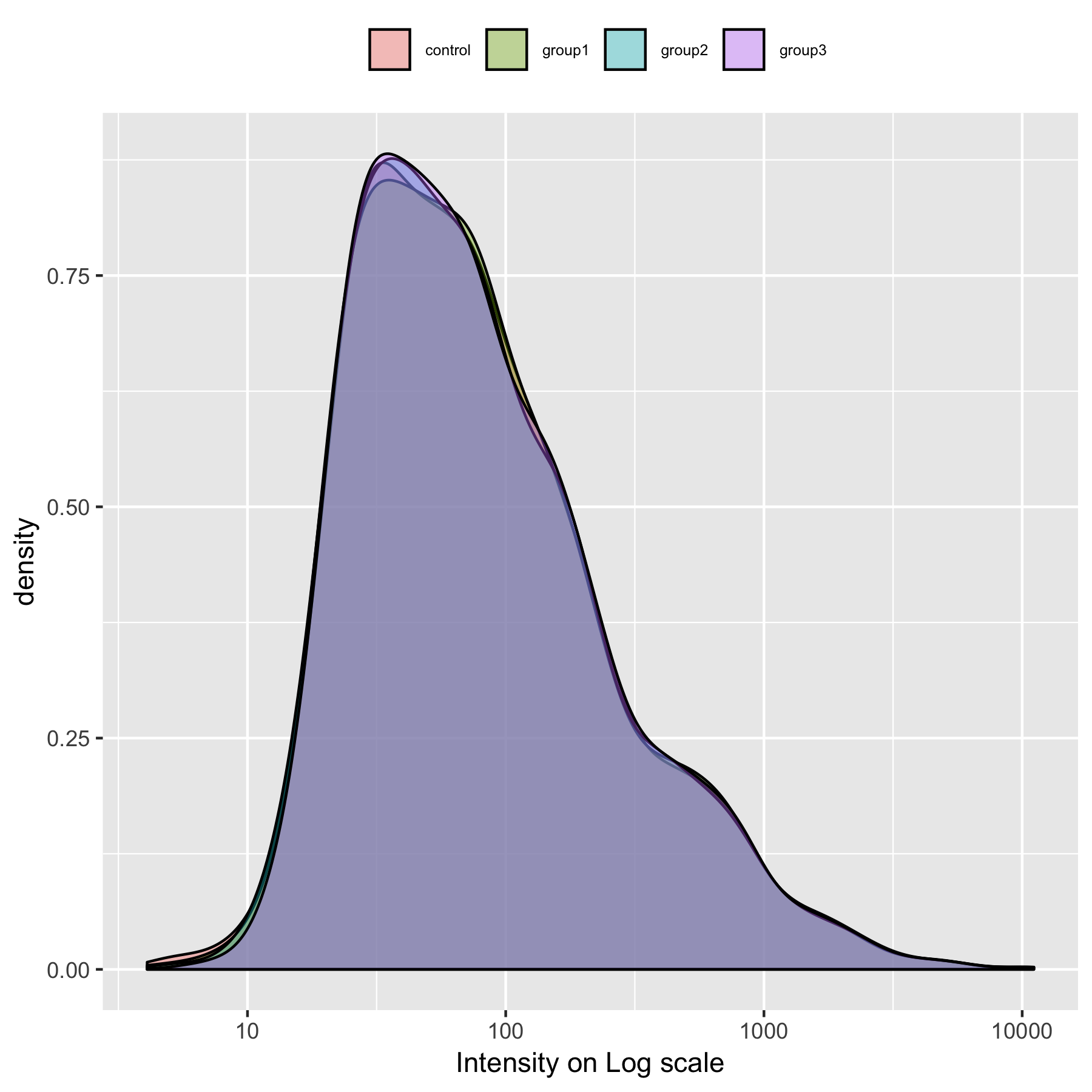

Figure S2. Distribution of peaks’ intensities on Log scale from Dataset 3.

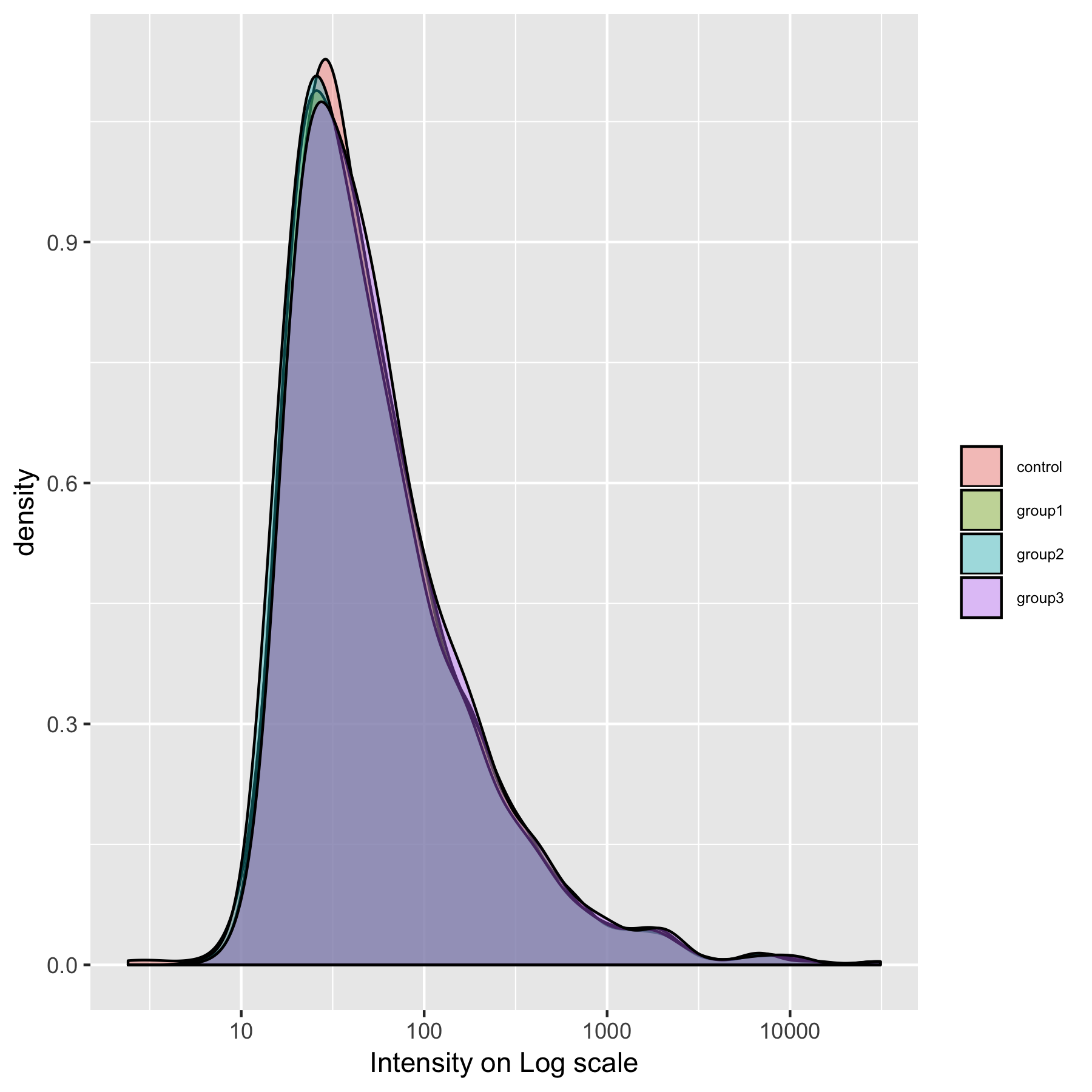

Figure S3. Distribution of peaks’ intensities on Log scale from Dataset 4.

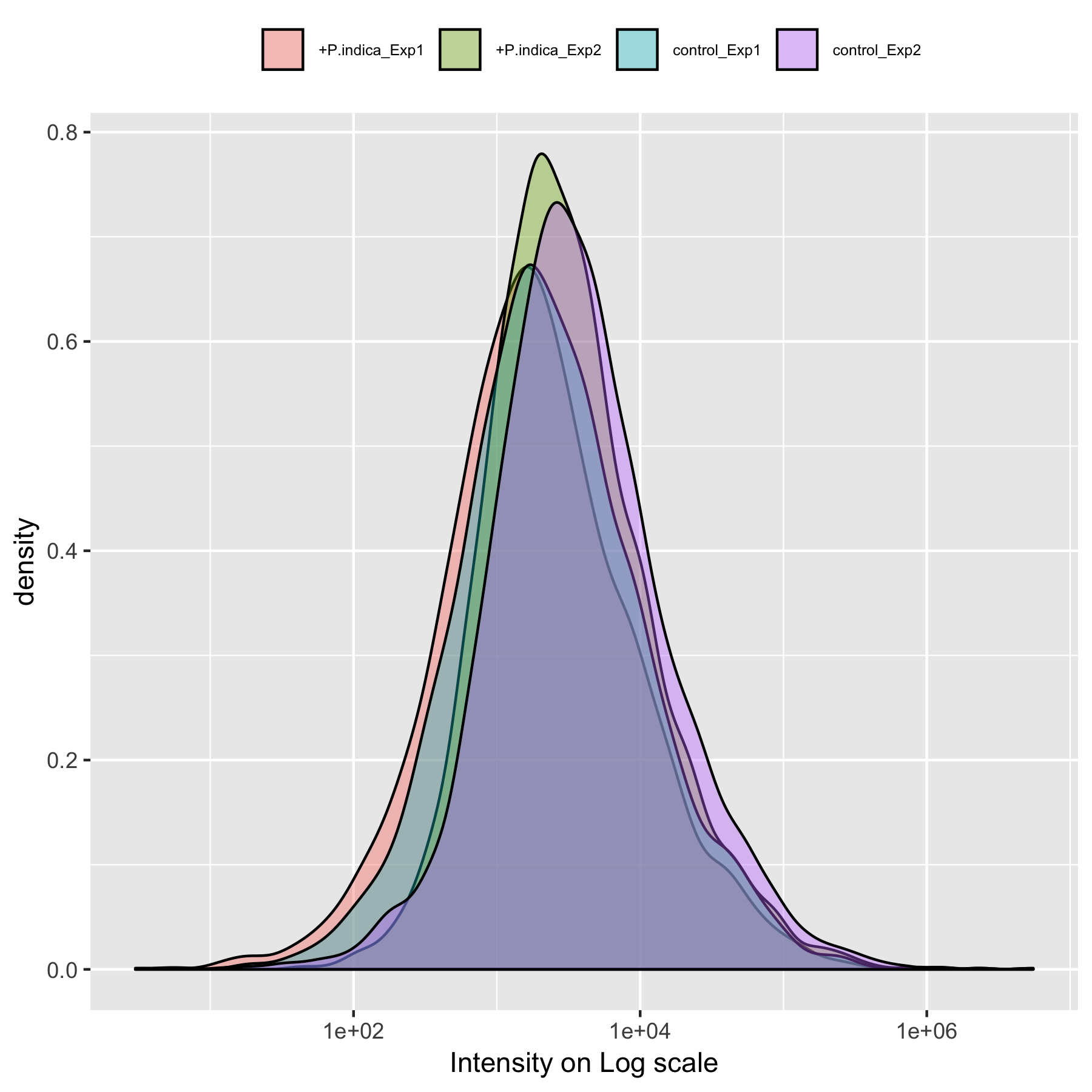

Figure S4. Distribution of peaks’ intensities on Log scale from Dataset 5.

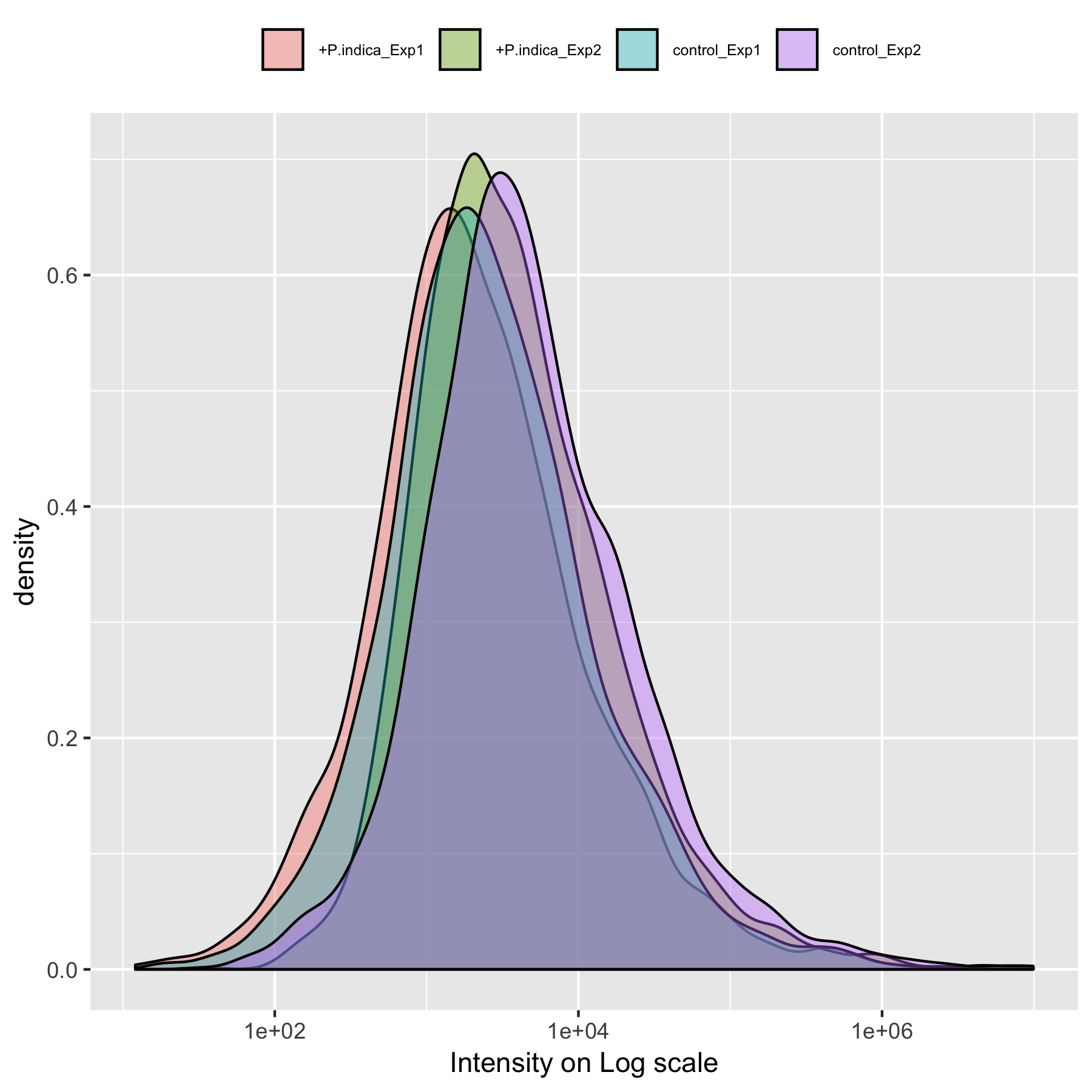

Figure S5. Distribution of peaks’ intensities on Log scale from Dataset 6.

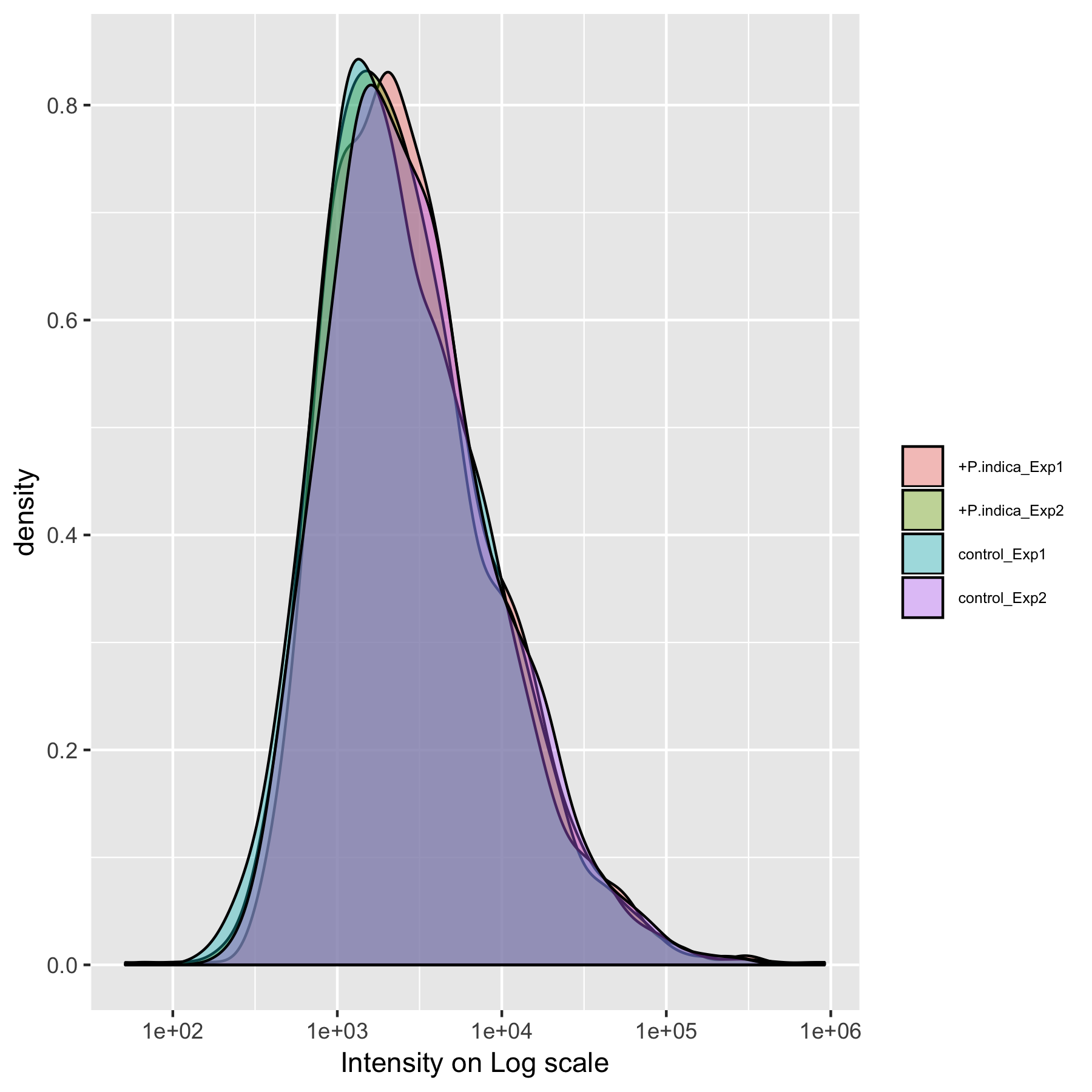

Figure S6. Distribution of peaks’ intensities on Log scale from Dataset 7.

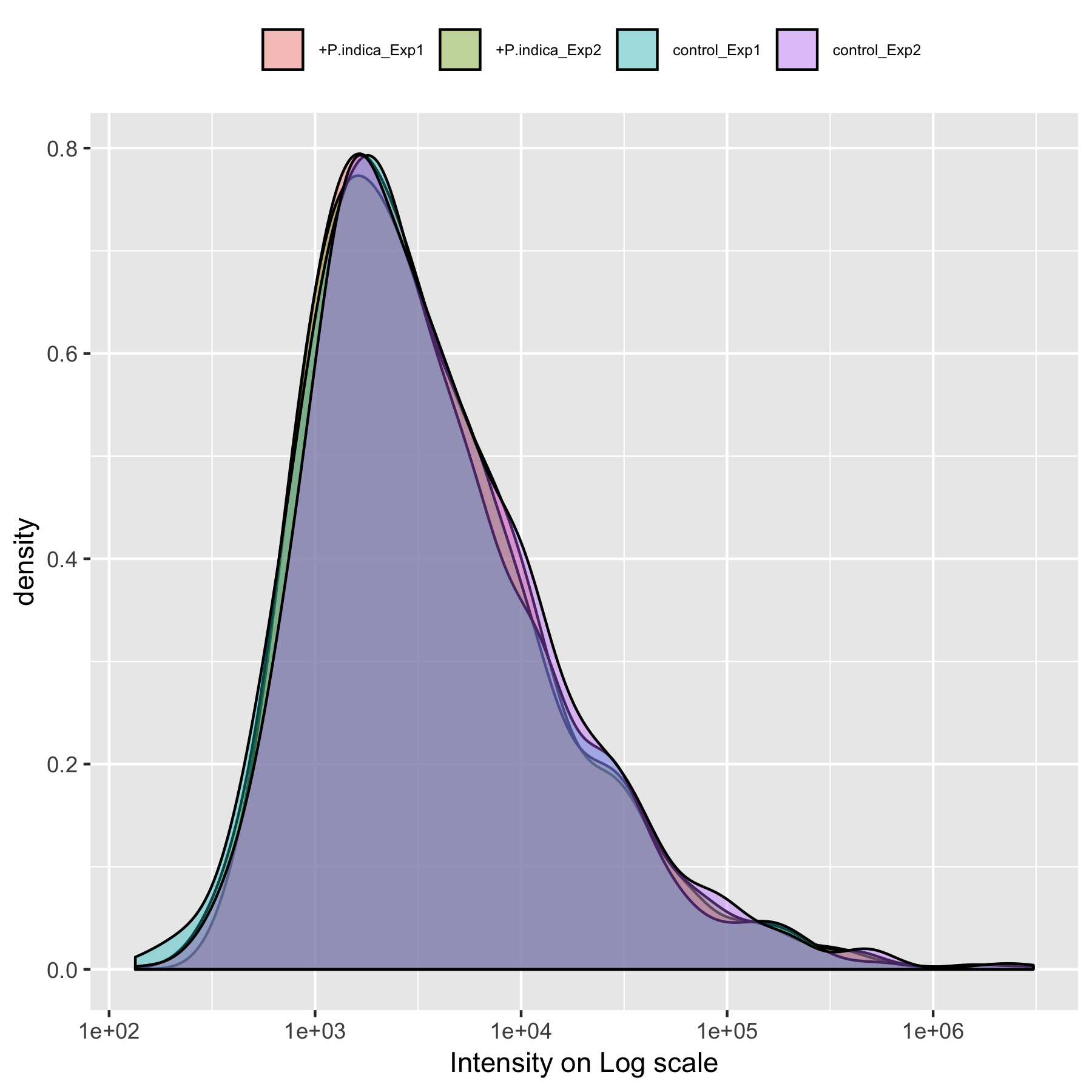

Figure S7. Distribution of peaks’ intensities on Log scale from Dataset 8.

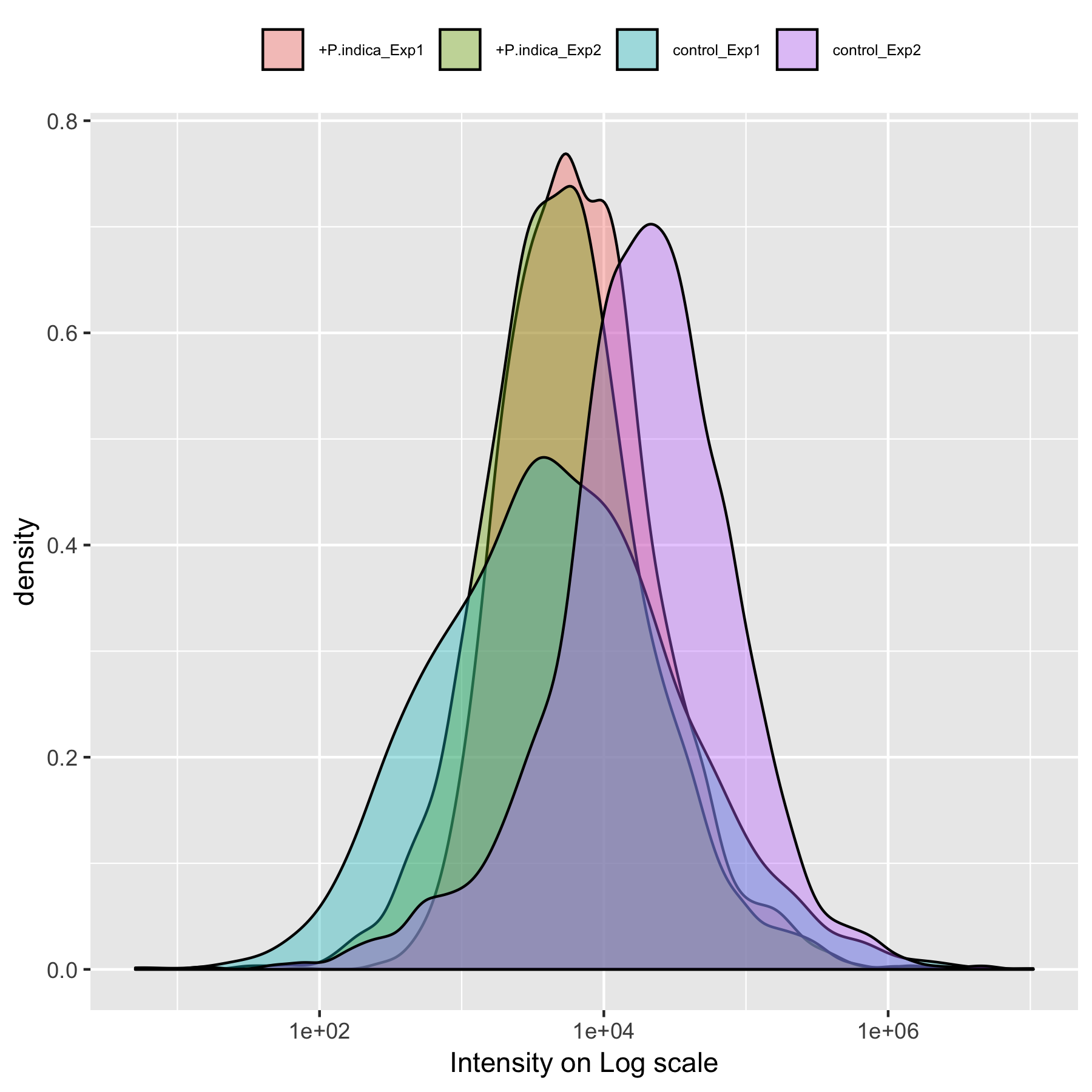

Figure S8. Distribution of peaks’ intensities on Log scale from Dataset 9.

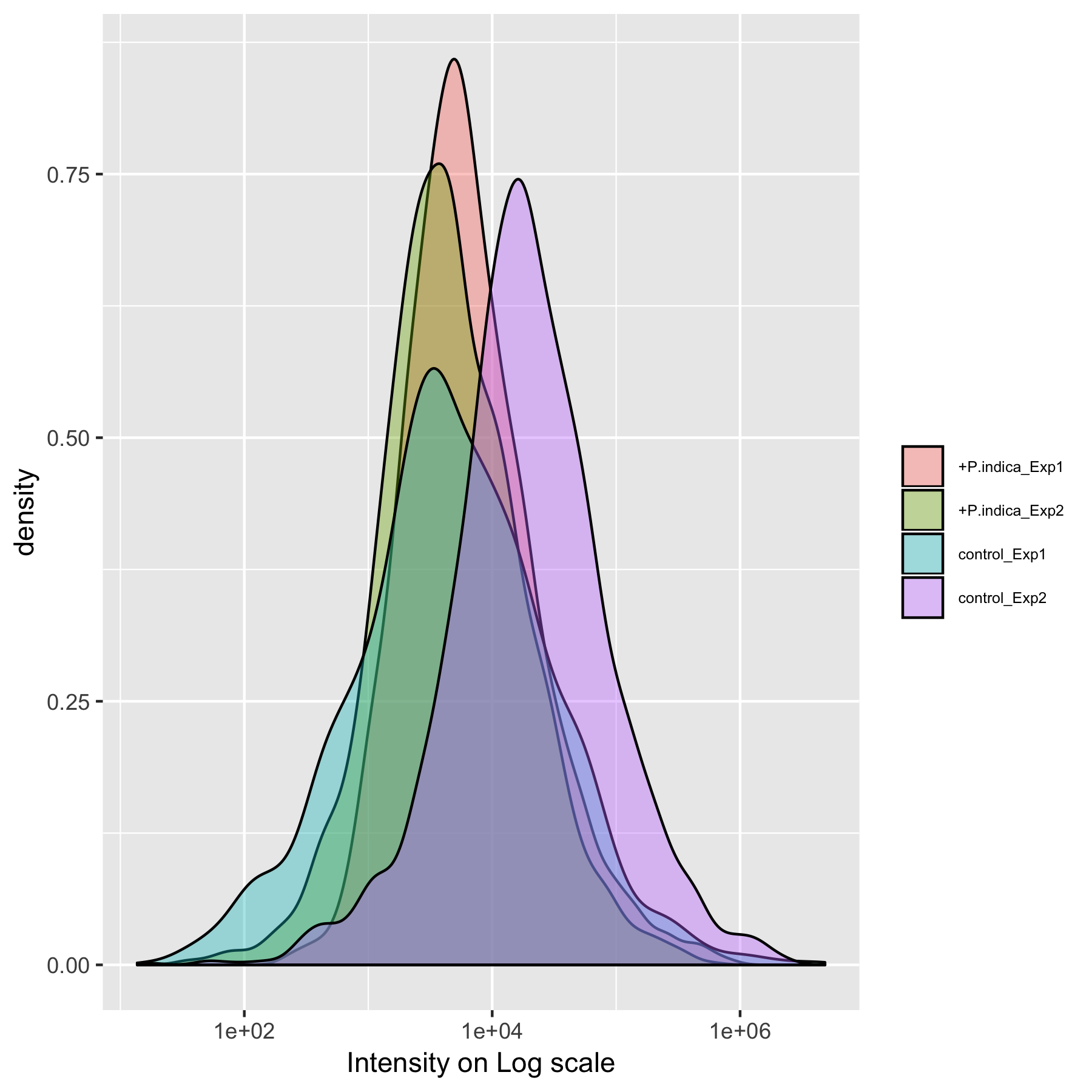

Figure S9. Distribution of peaks’ intensities on Log scale from Dataset 10.

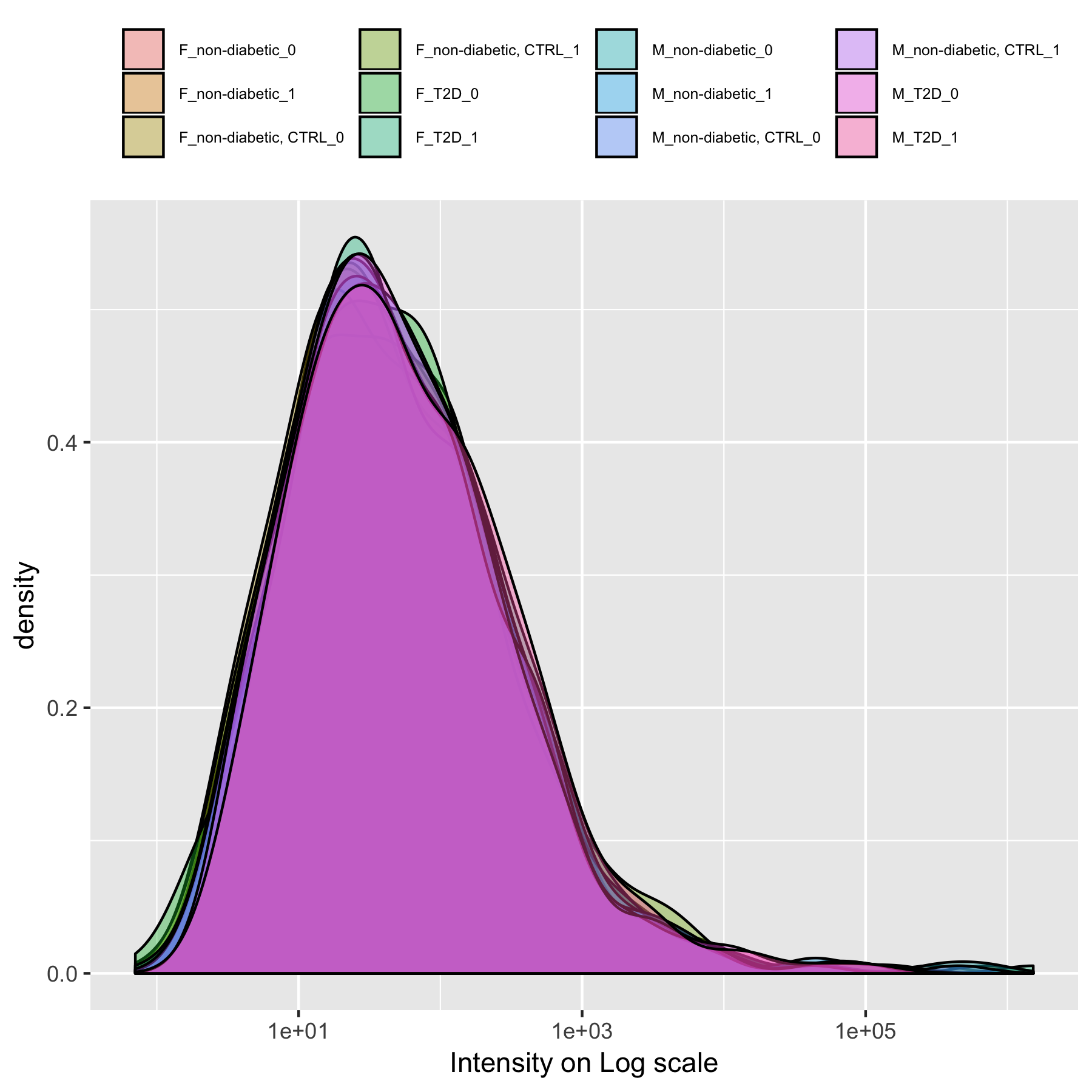

Figure S10. Distribution of peaks’ intensities on Log scale from Dataset 11.

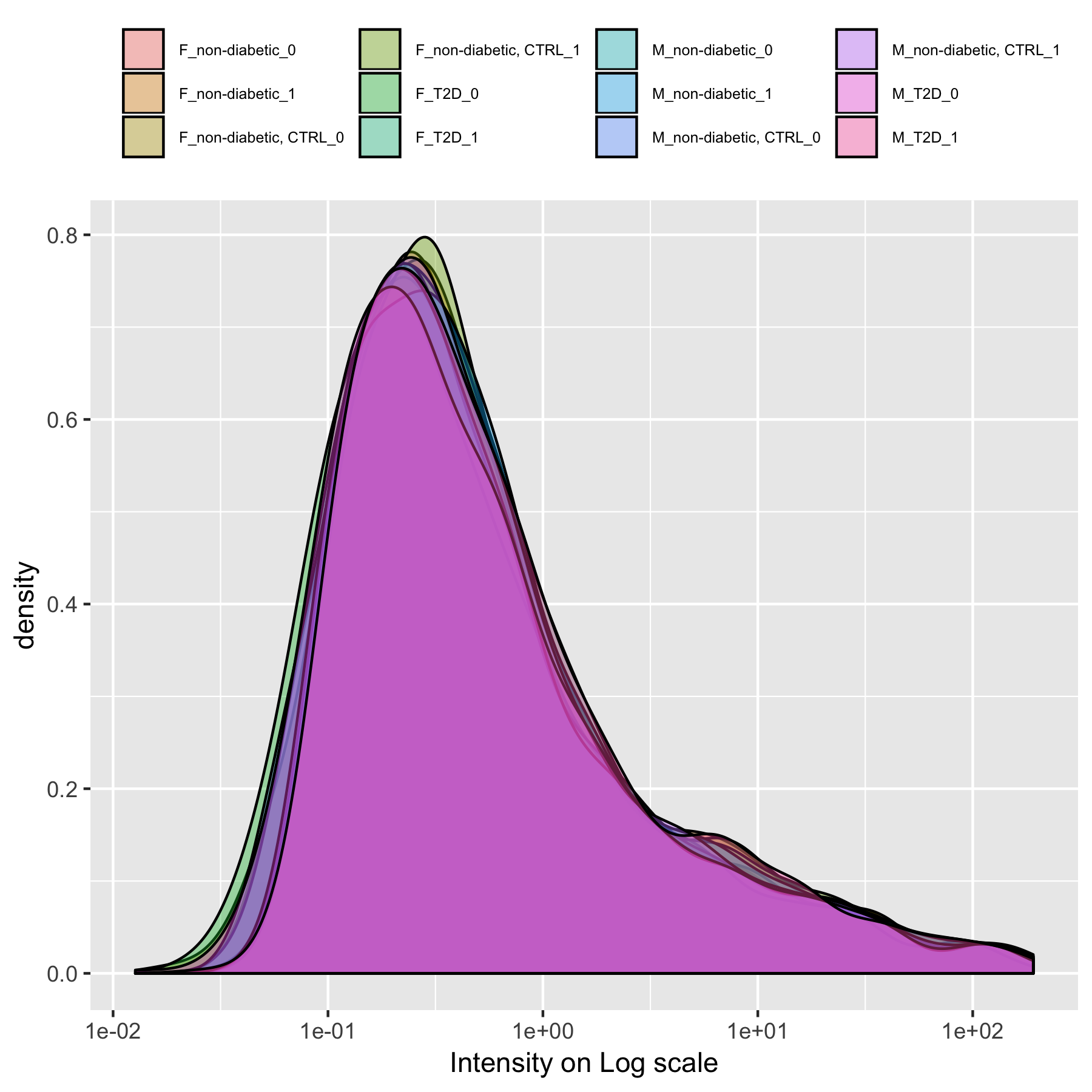

Figure S11. Distribution of peaks’ intensities on Log scale from Dataset 12.

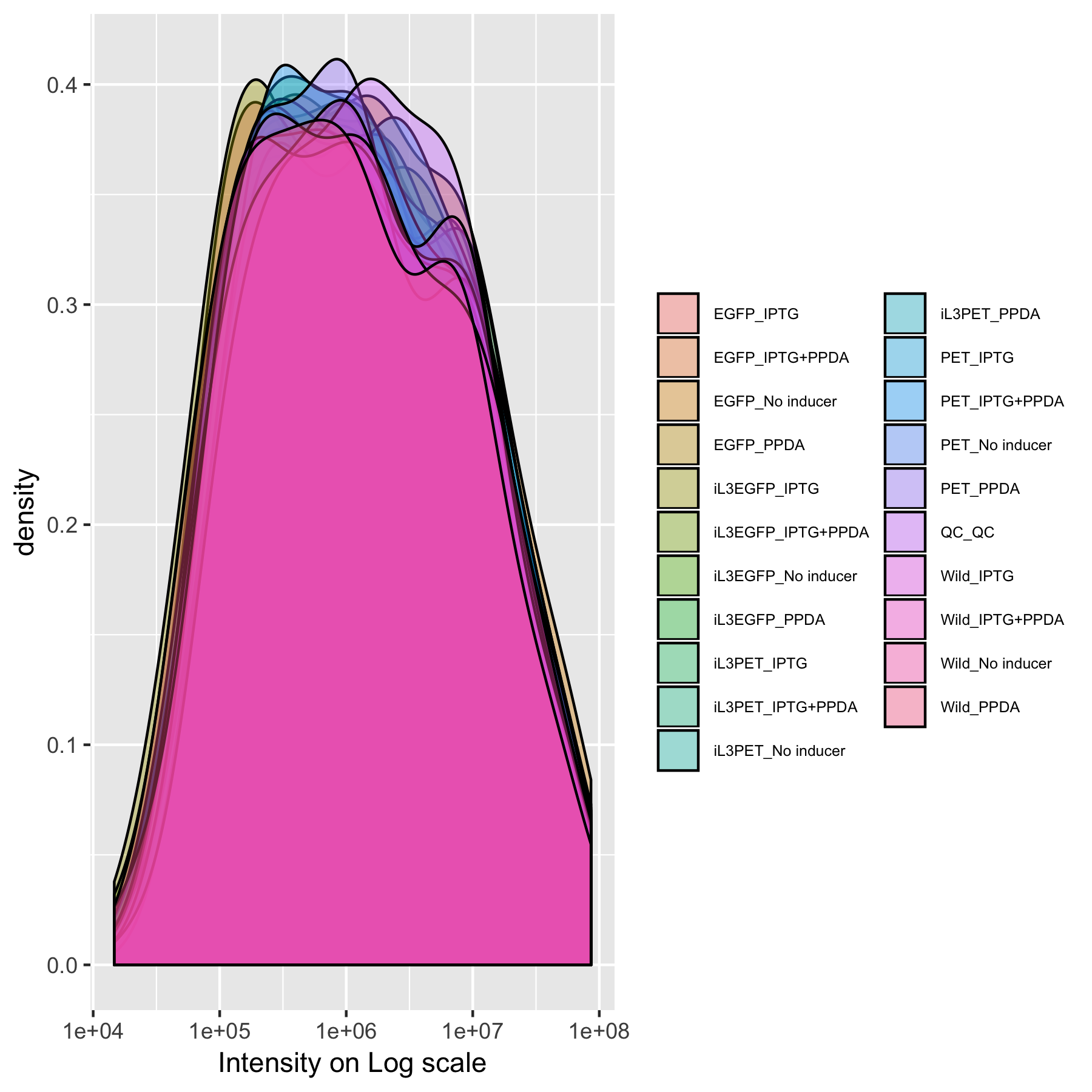

Figure S12. Distribution of peaks’ intensities on Log scale from Dataset 13.

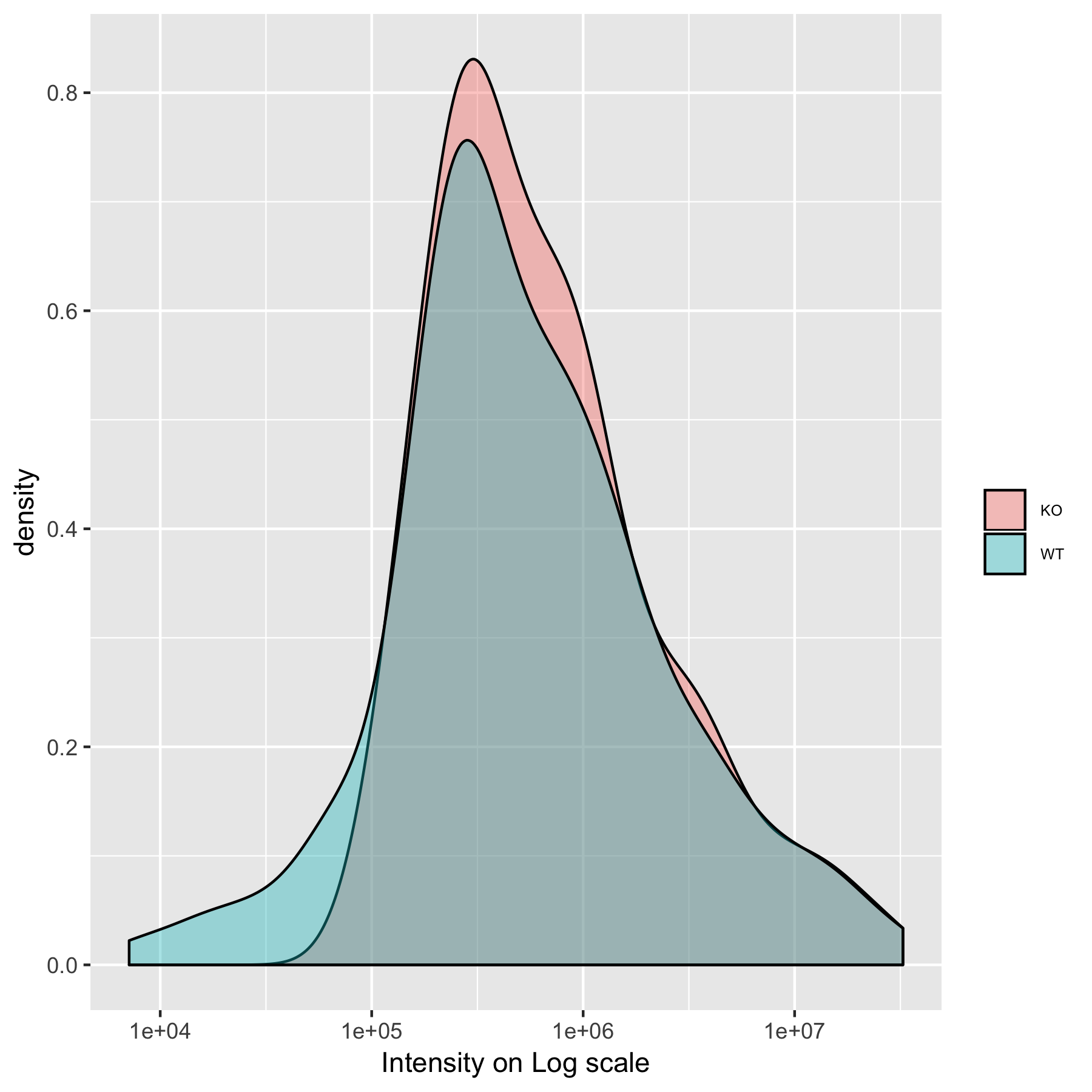

Figure S13. Distribution of peaks’ intensities on Log scale from Dataset 14.

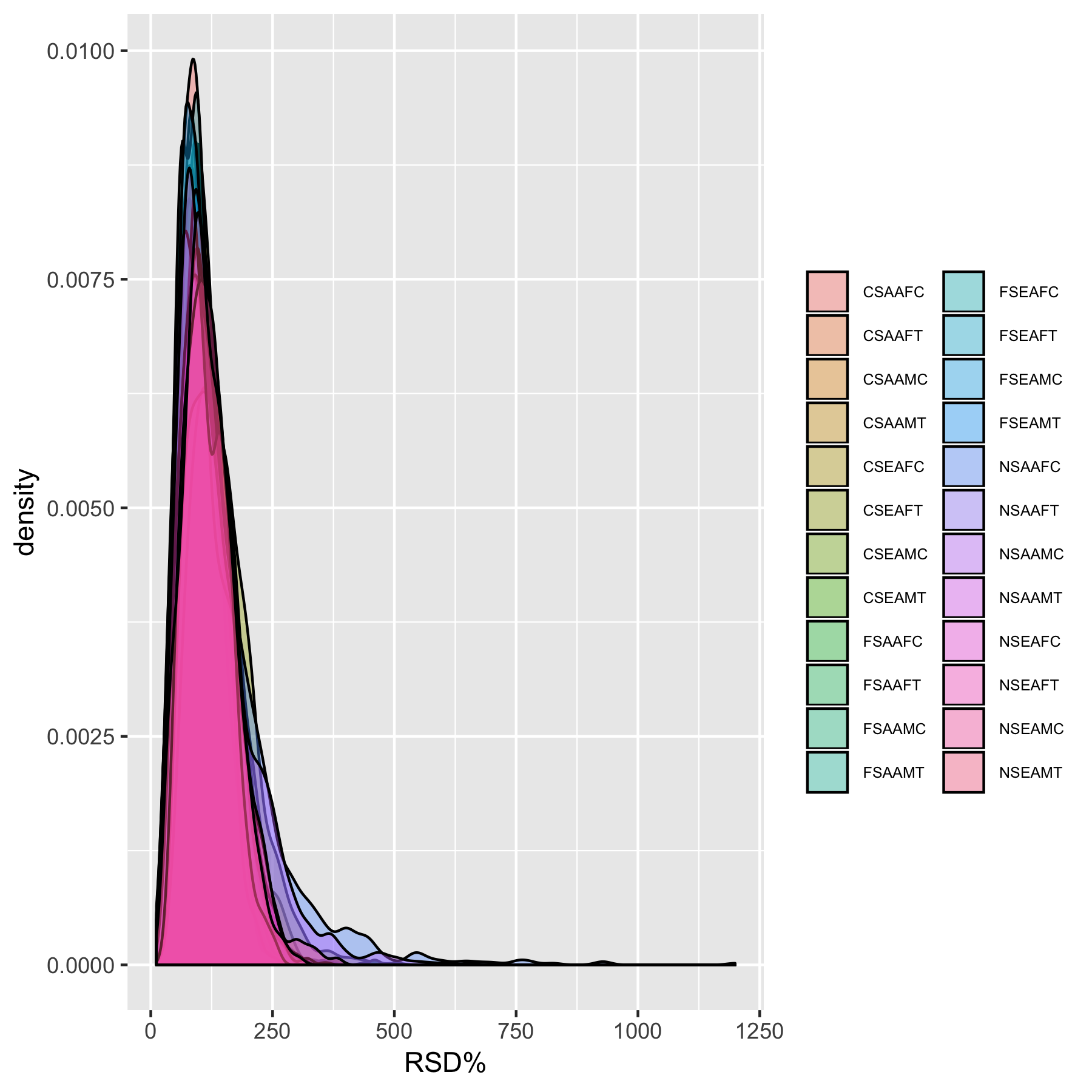

Figure S14. RSD% Distribution of peaks’ intensities on Log scale within groups from Dataset 2.

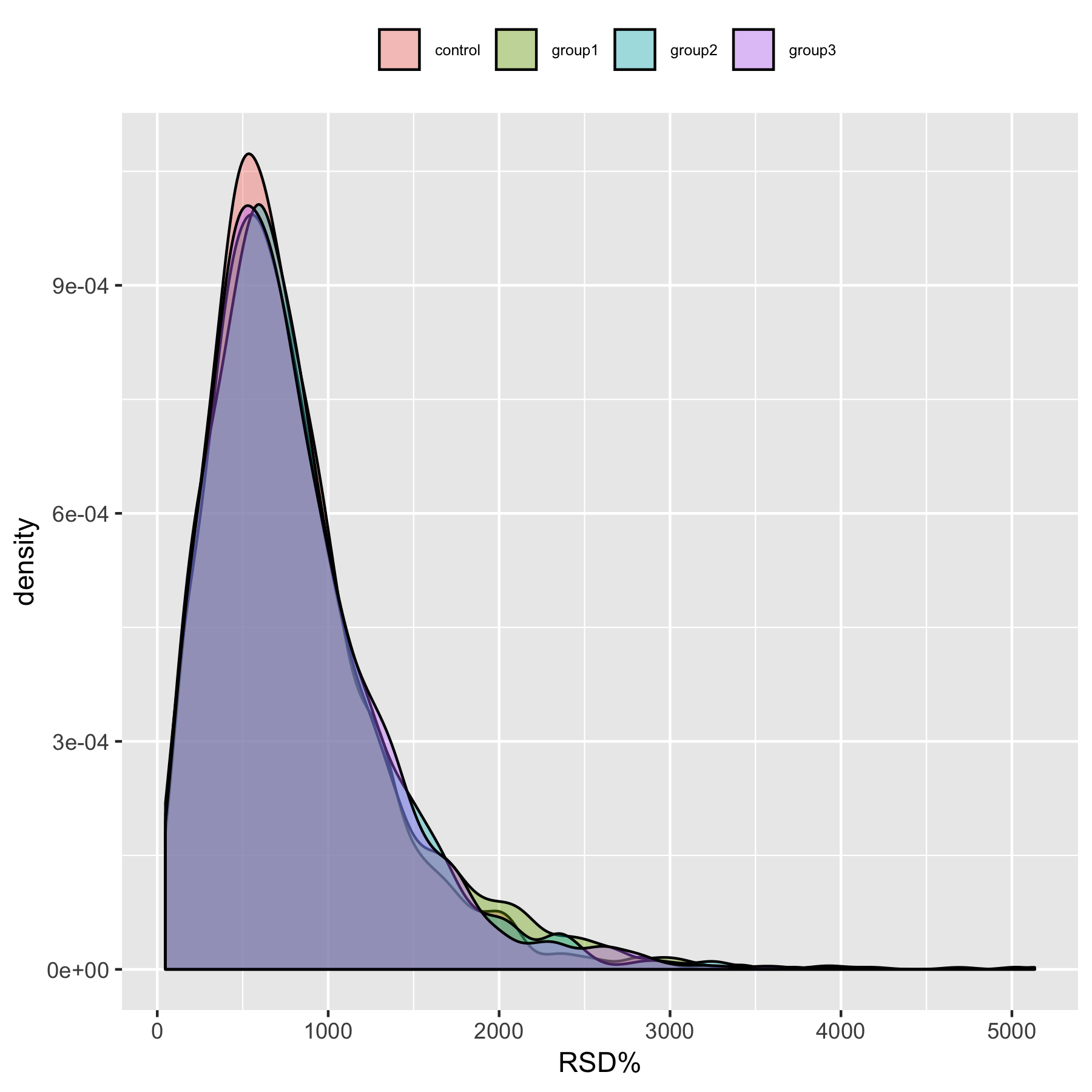

Figure S15. RSD% Distribution of peaks’ intensities on Log scale within groups from Dataset 3.

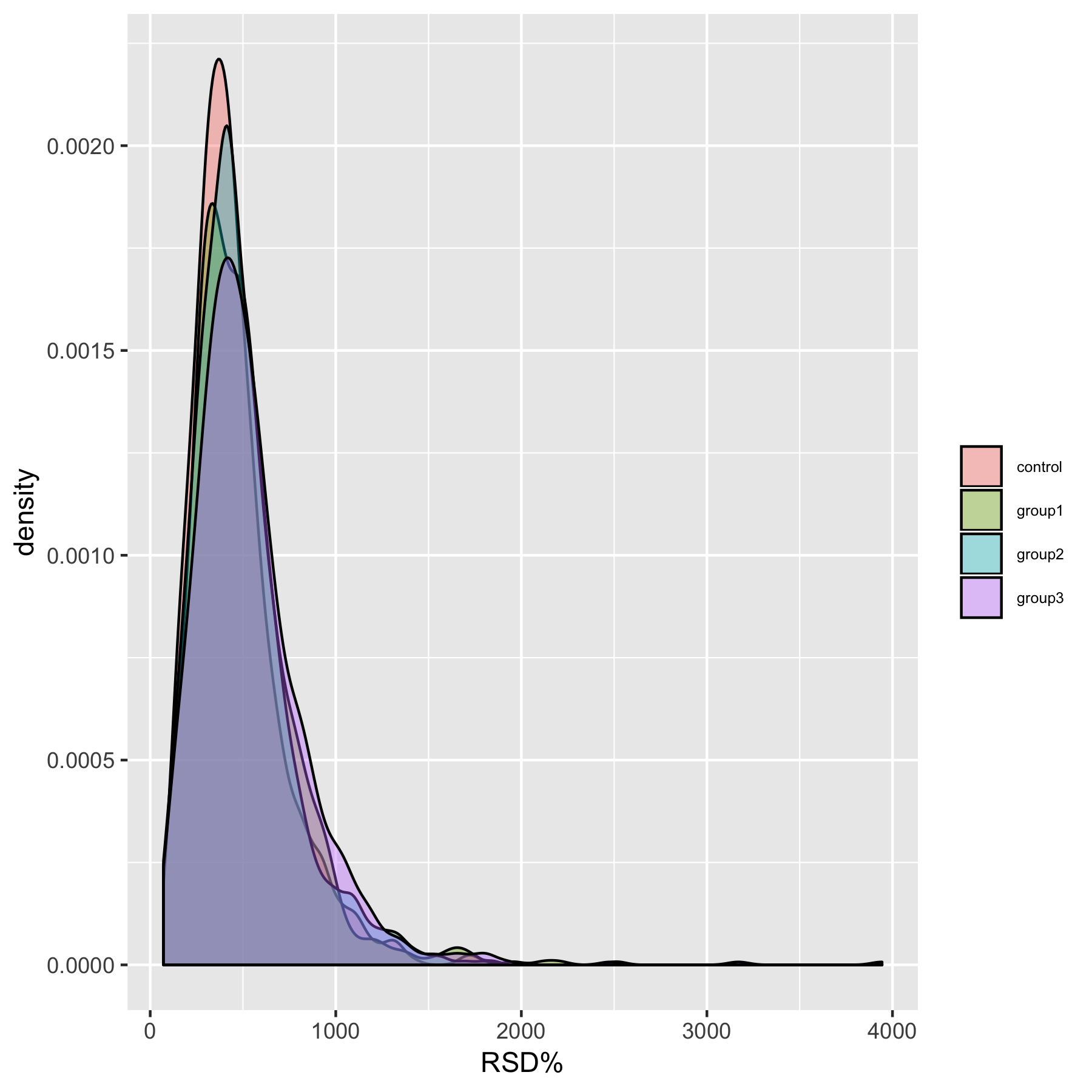

Figure S16. RSD% Distribution of peaks’ intensities on Log scale within groups from Dataset 4.

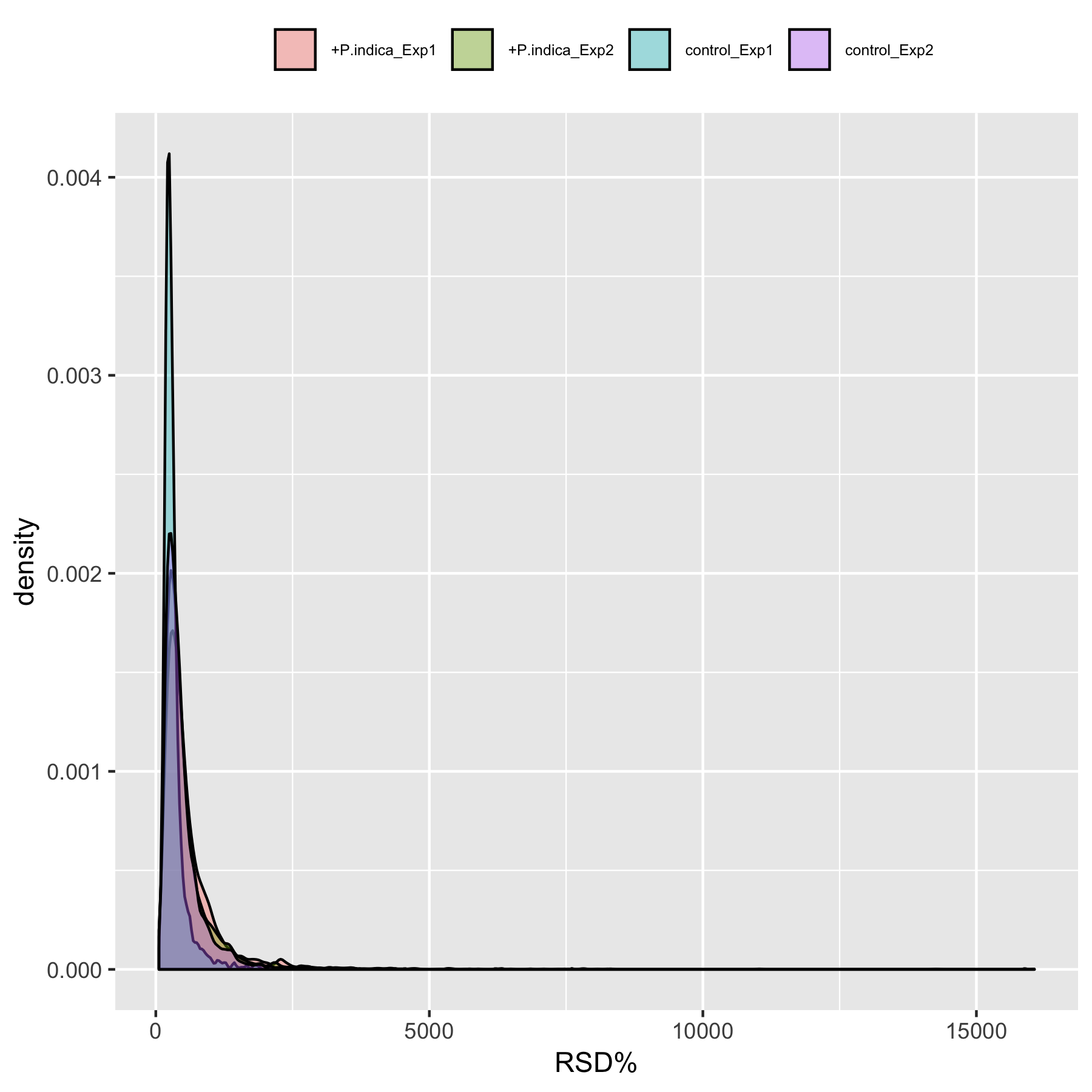

Figure S17. RSD% Distribution of peaks’ intensities on Log scale within groups from Dataset 5.

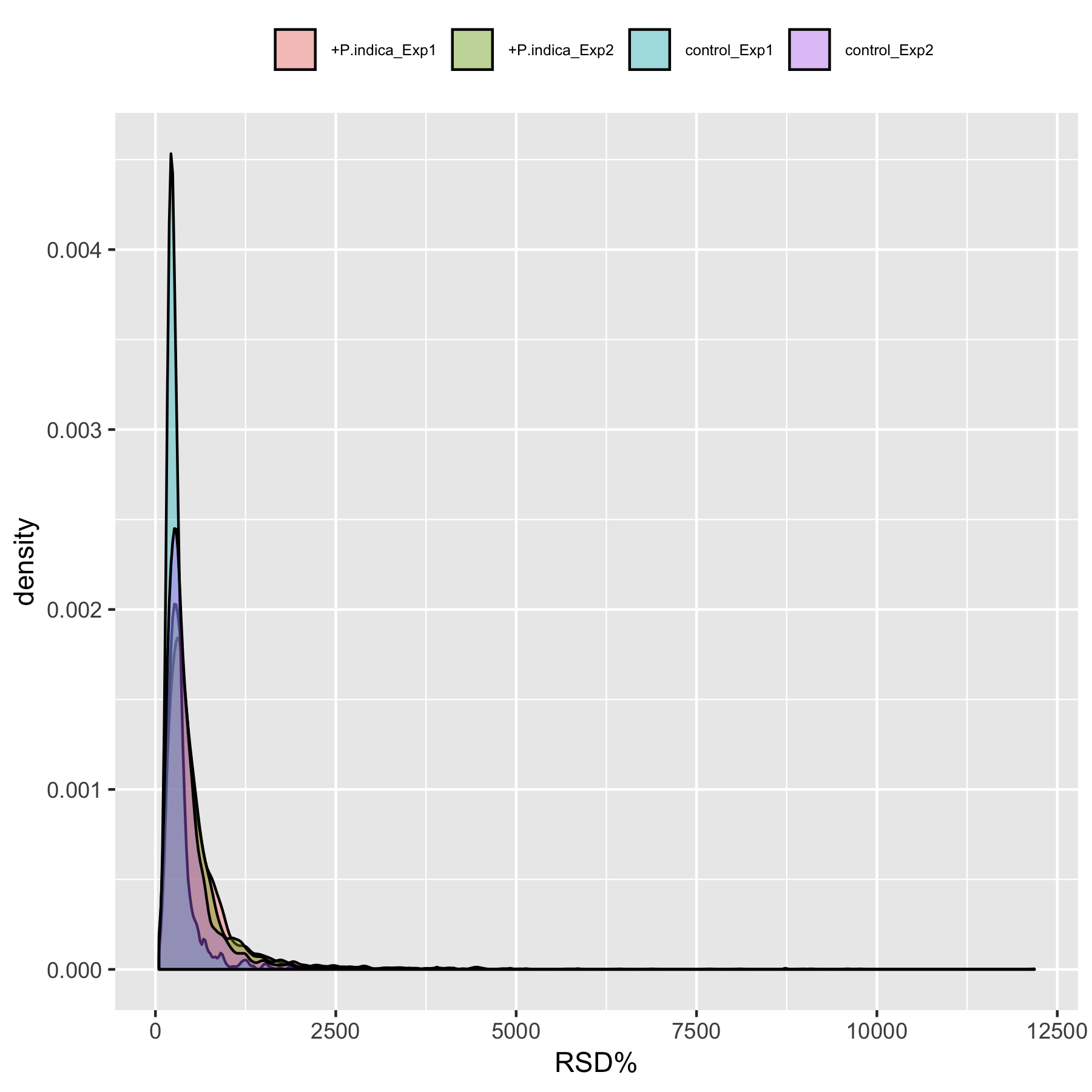

Figure S18. RSD% Distribution of peaks’ intensities on Log scale within groups from Dataset 6.

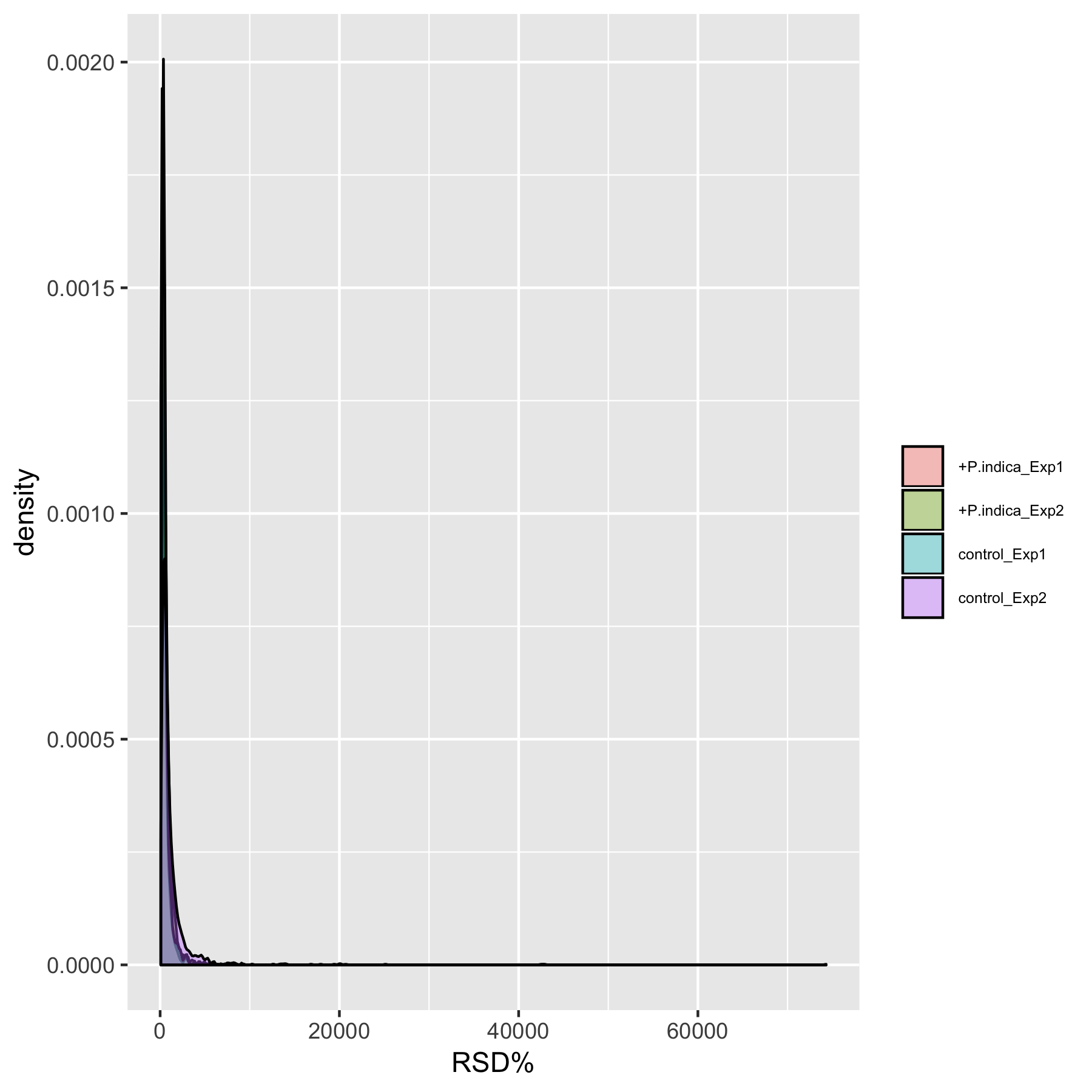

Figure S19. RSD% Distribution of peaks’ intensities on Log scale within groups from Dataset 7.

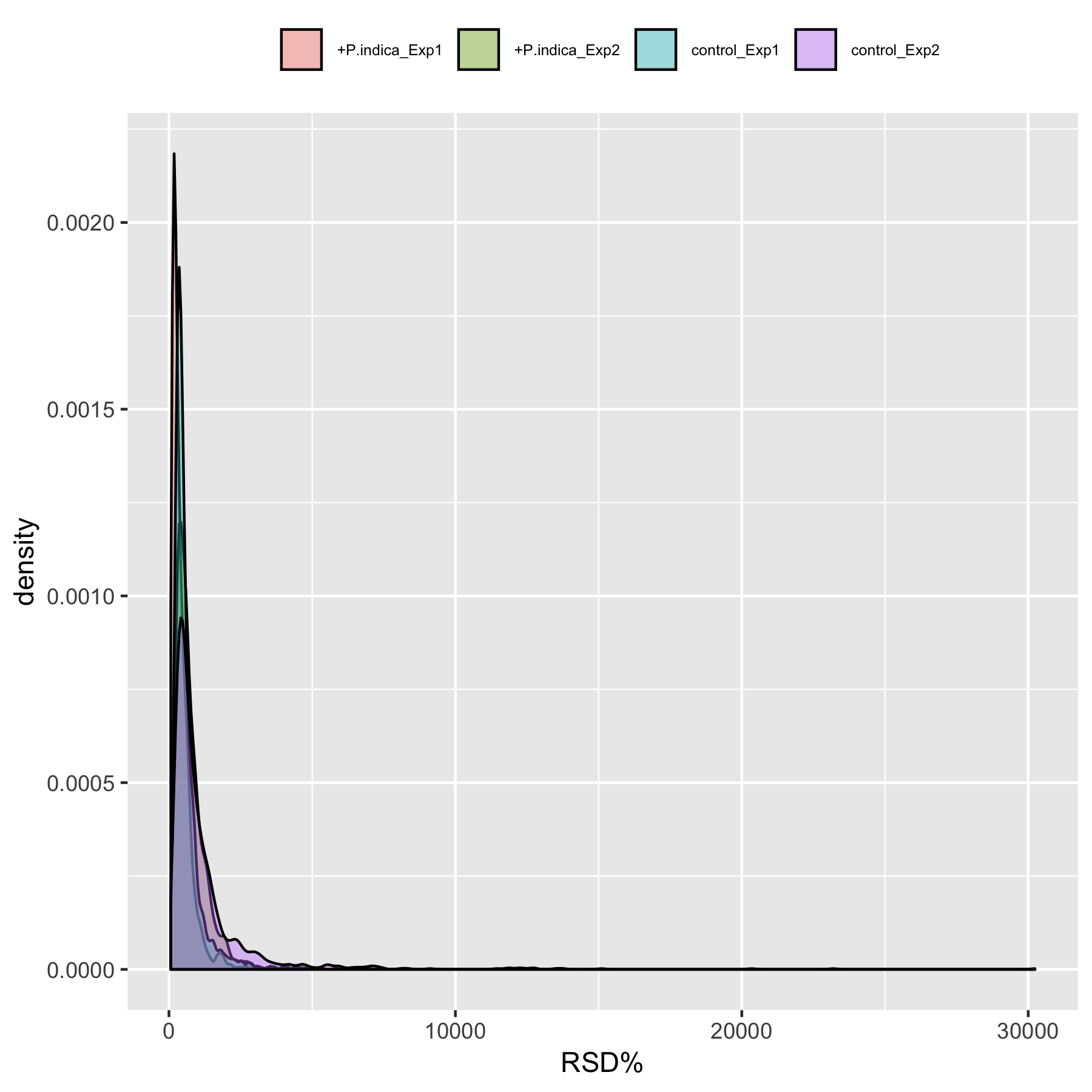

Figure S20. RSD% Distribution of peaks’ intensities on Log scale within groups from Dataset 8.

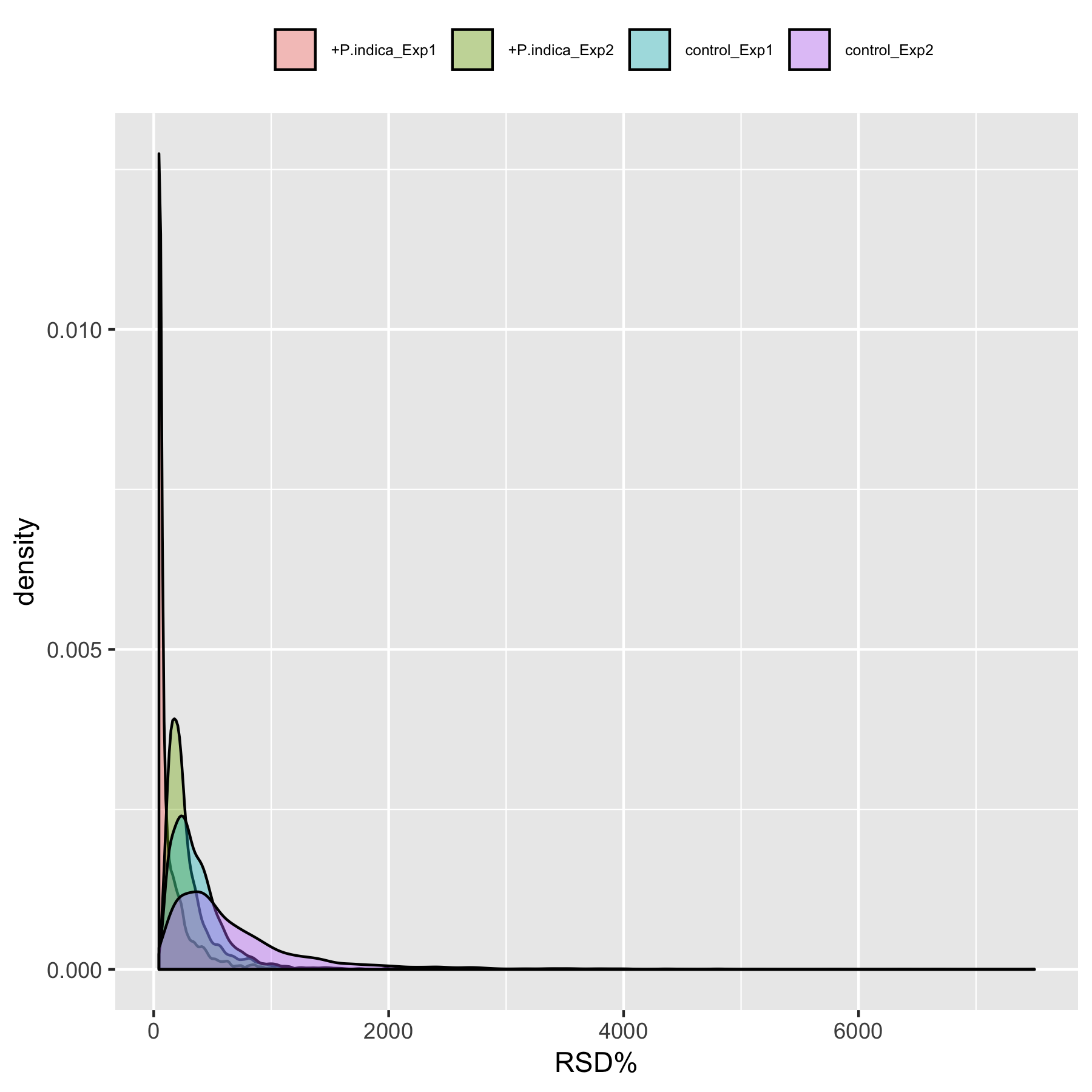

Figure S21. RSD% Distribution of peaks’ intensities on Log scale within groups from Dataset 9.

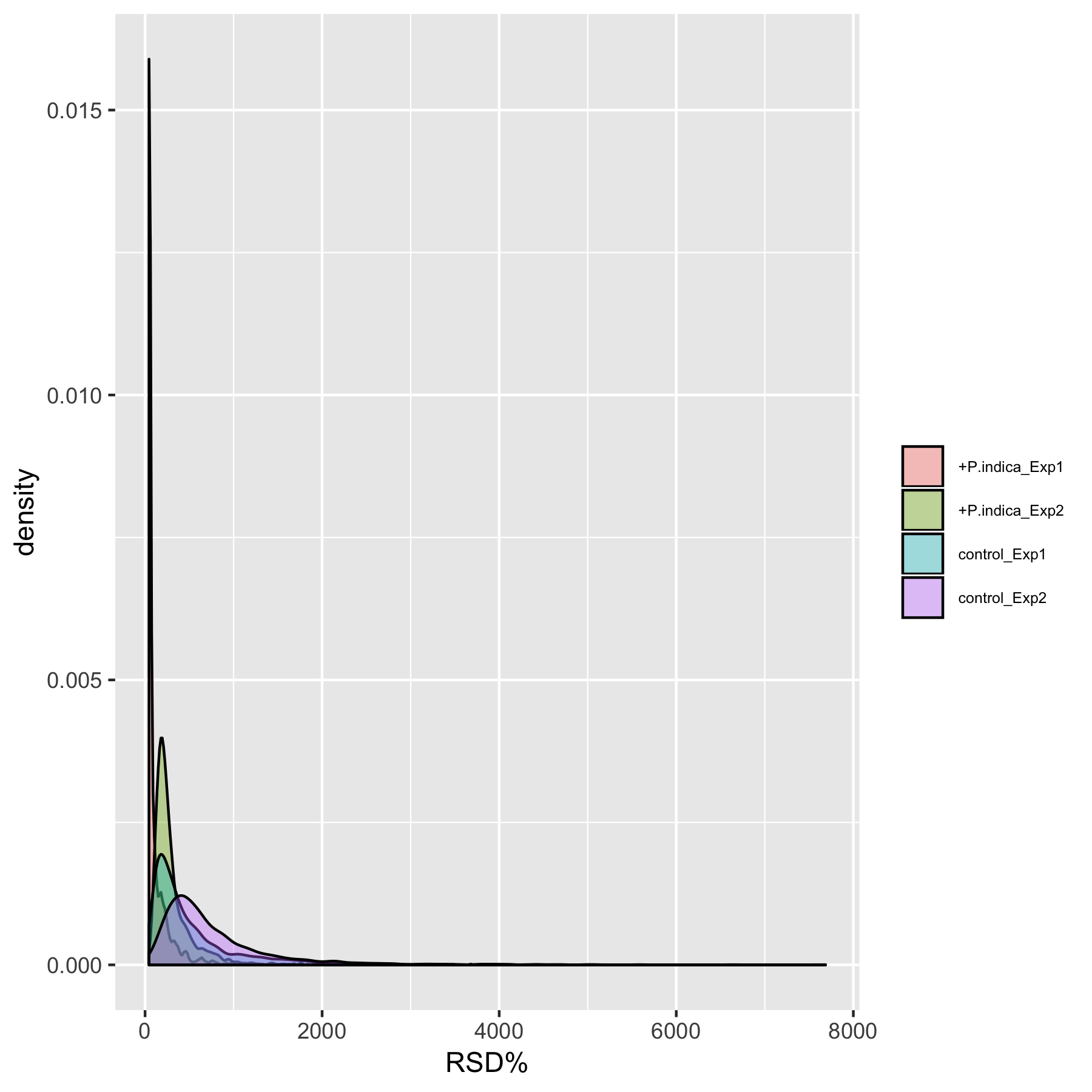

Figure S22. RSD% Distribution of peaks’ intensities on Log scale within groups from Dataset 10.

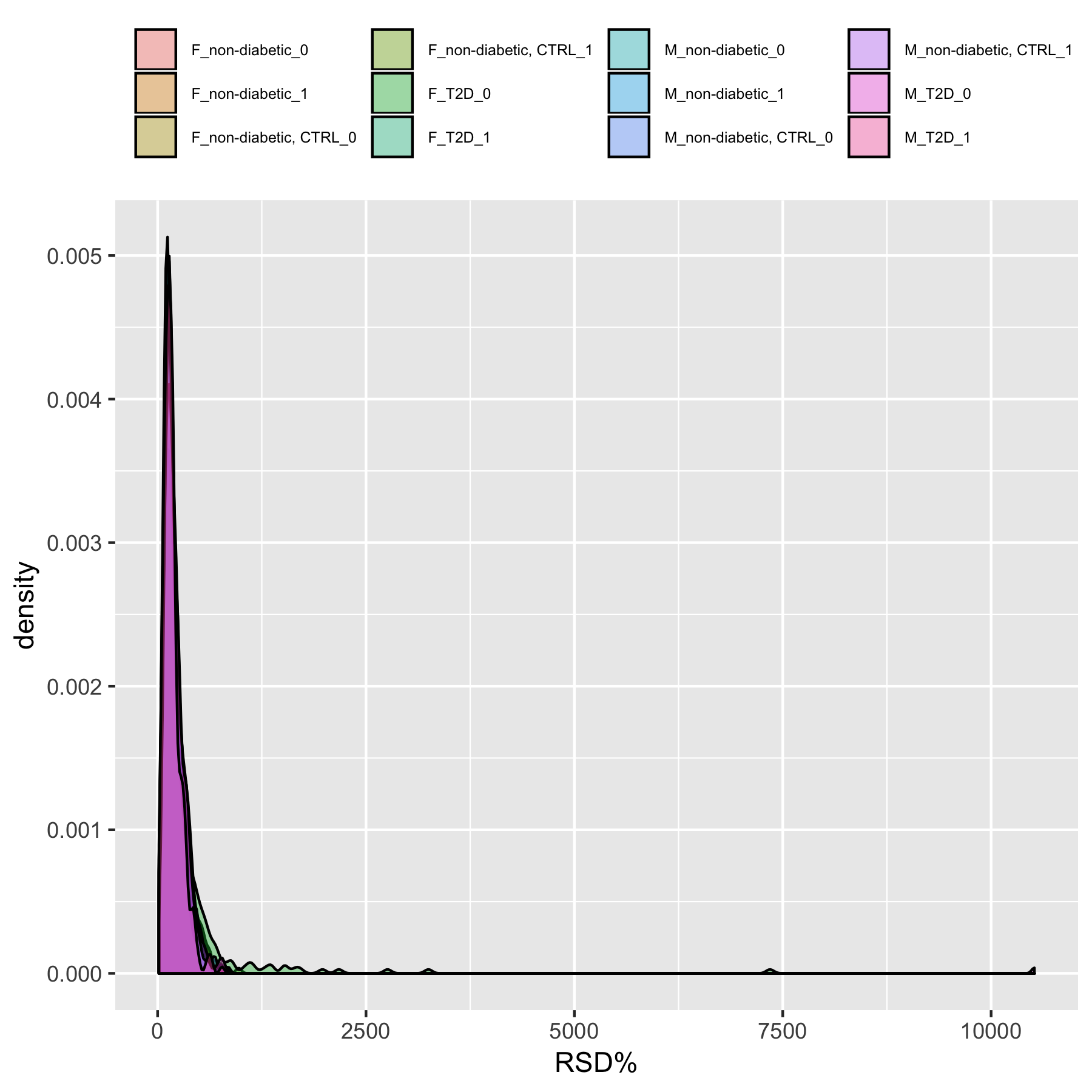

Figure S23. RSD% Distribution of peaks’ intensities on Log scale within groups from Dataset 11.

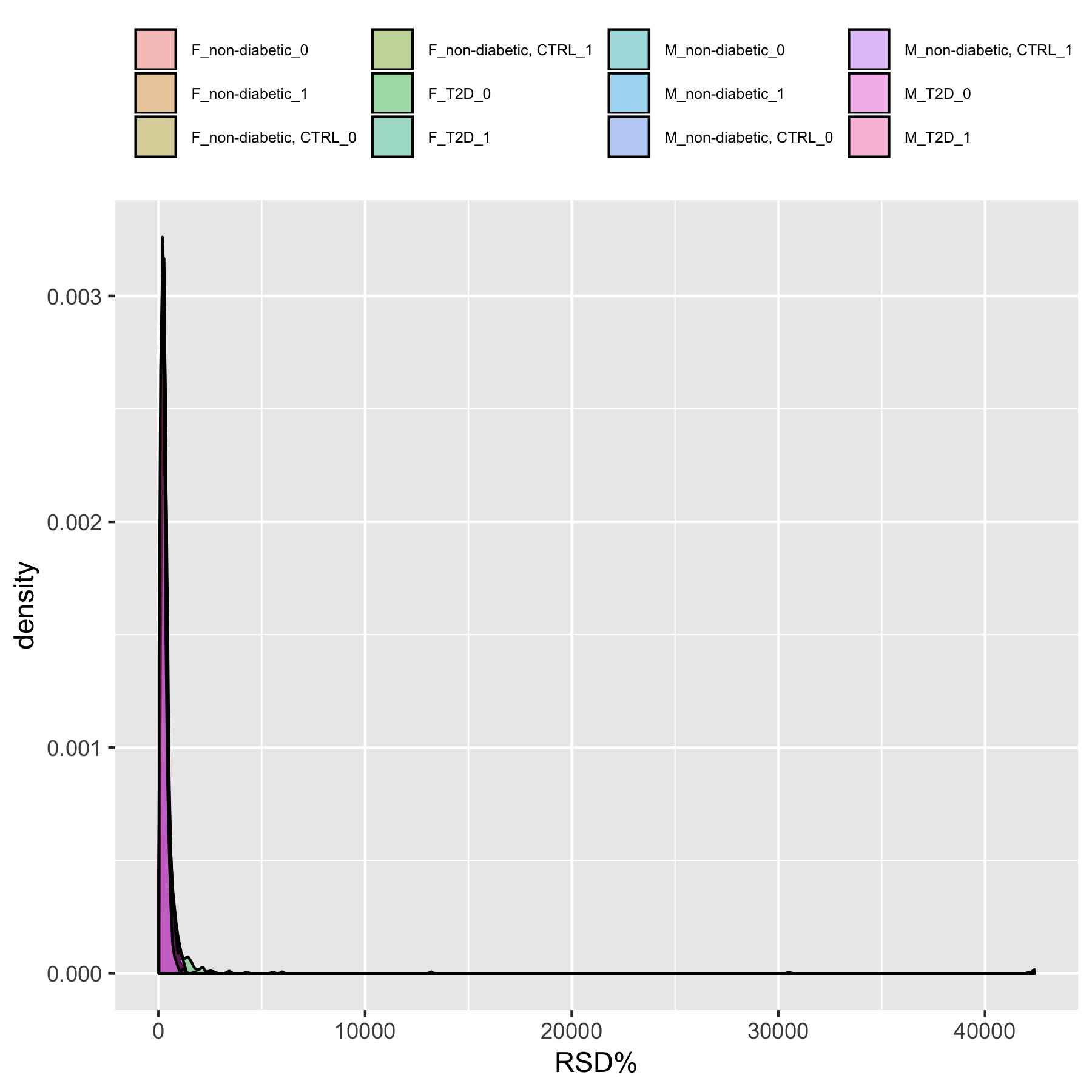

Figure S24. RSD% Distribution of peaks’ intensities on Log scale within groups from Dataset 12.

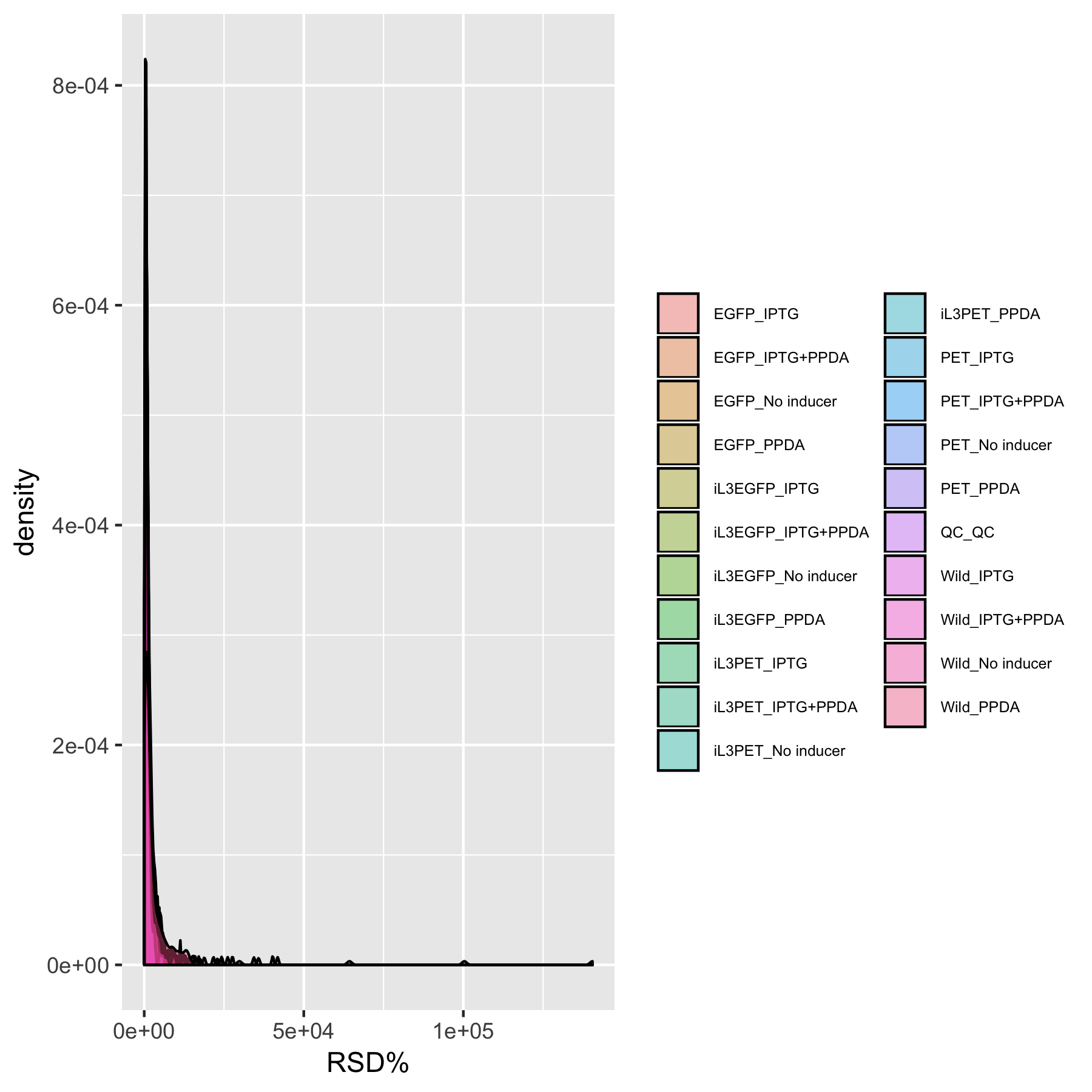

Figure S25. RSD% Distribution of peaks’ intensities on Log scale within groups from Dataset 13.

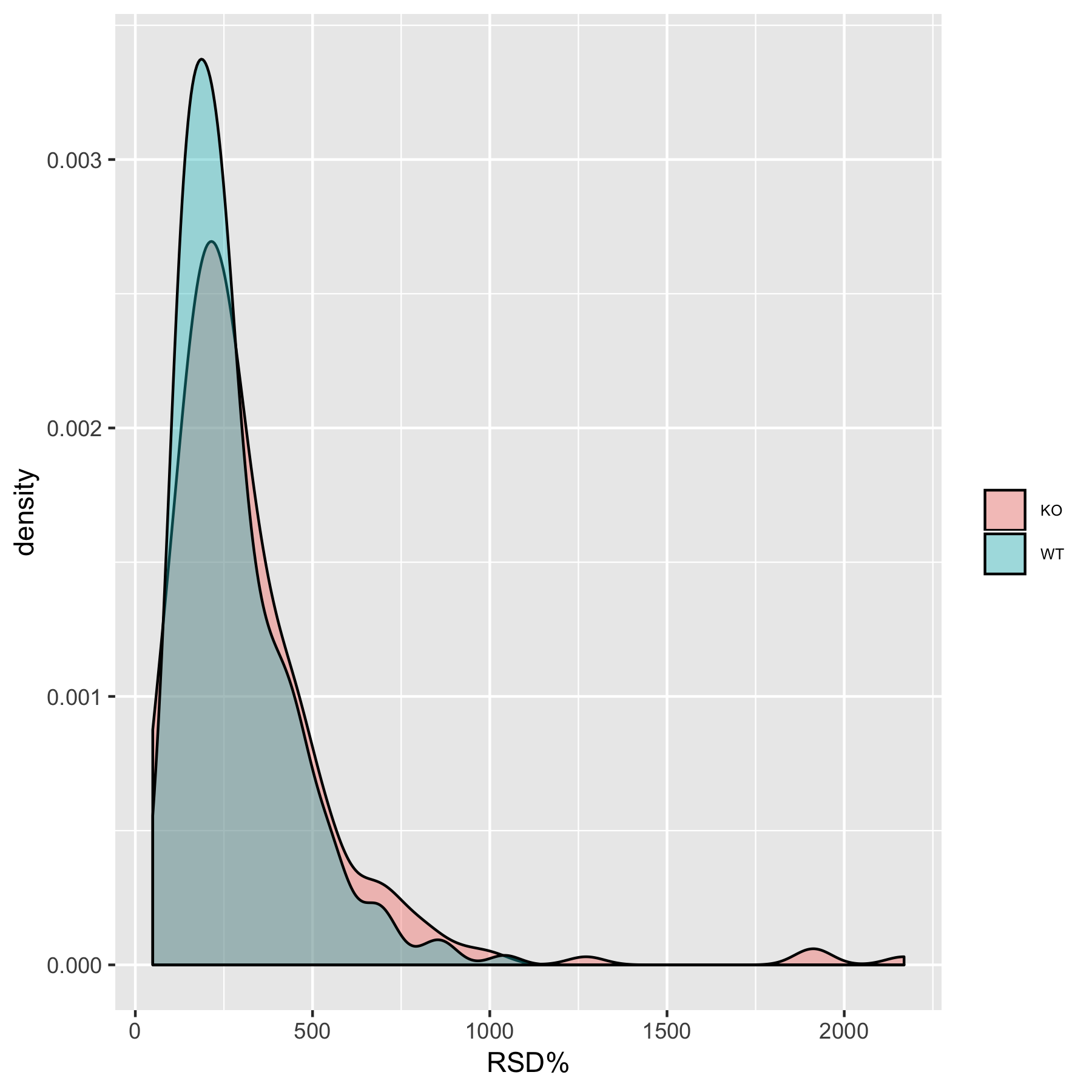

Figure S26. RSD% Distribution of peaks’ intensities on Log scale within groups from Dataset 14.

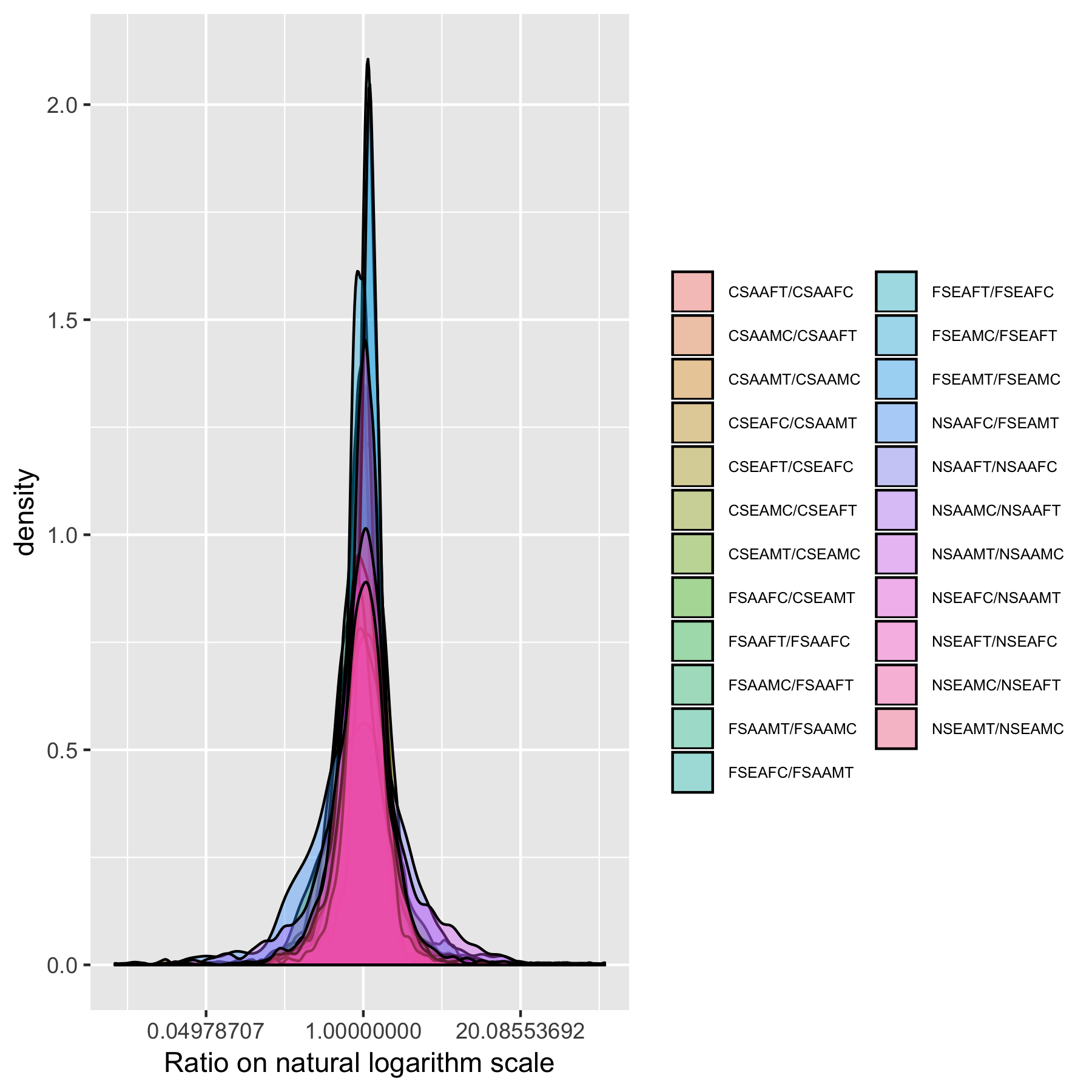

Figure S27. Ratios of group averages distribution from dataset 2.

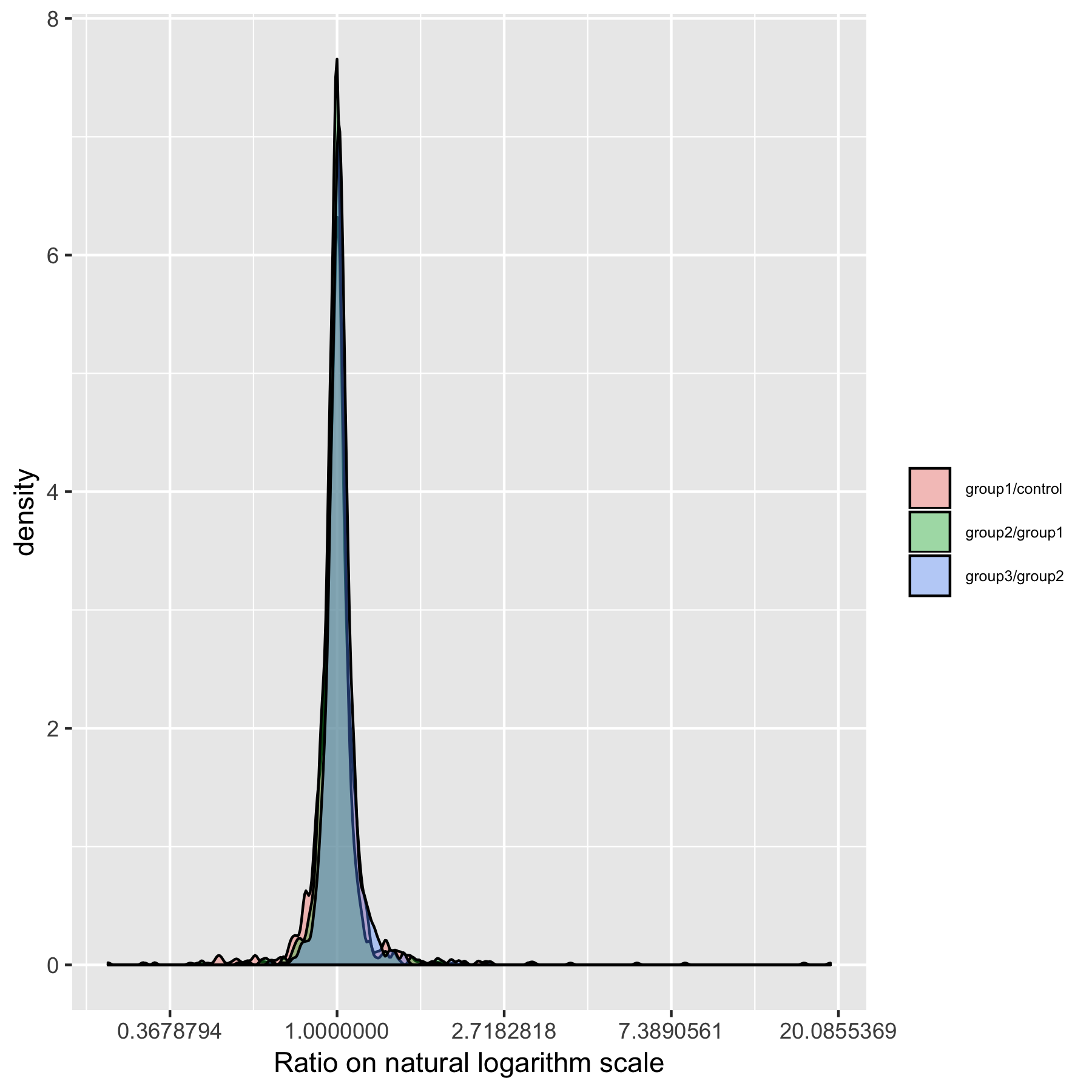

Figure S28. Ratios of group averages distribution from Dataset 3.

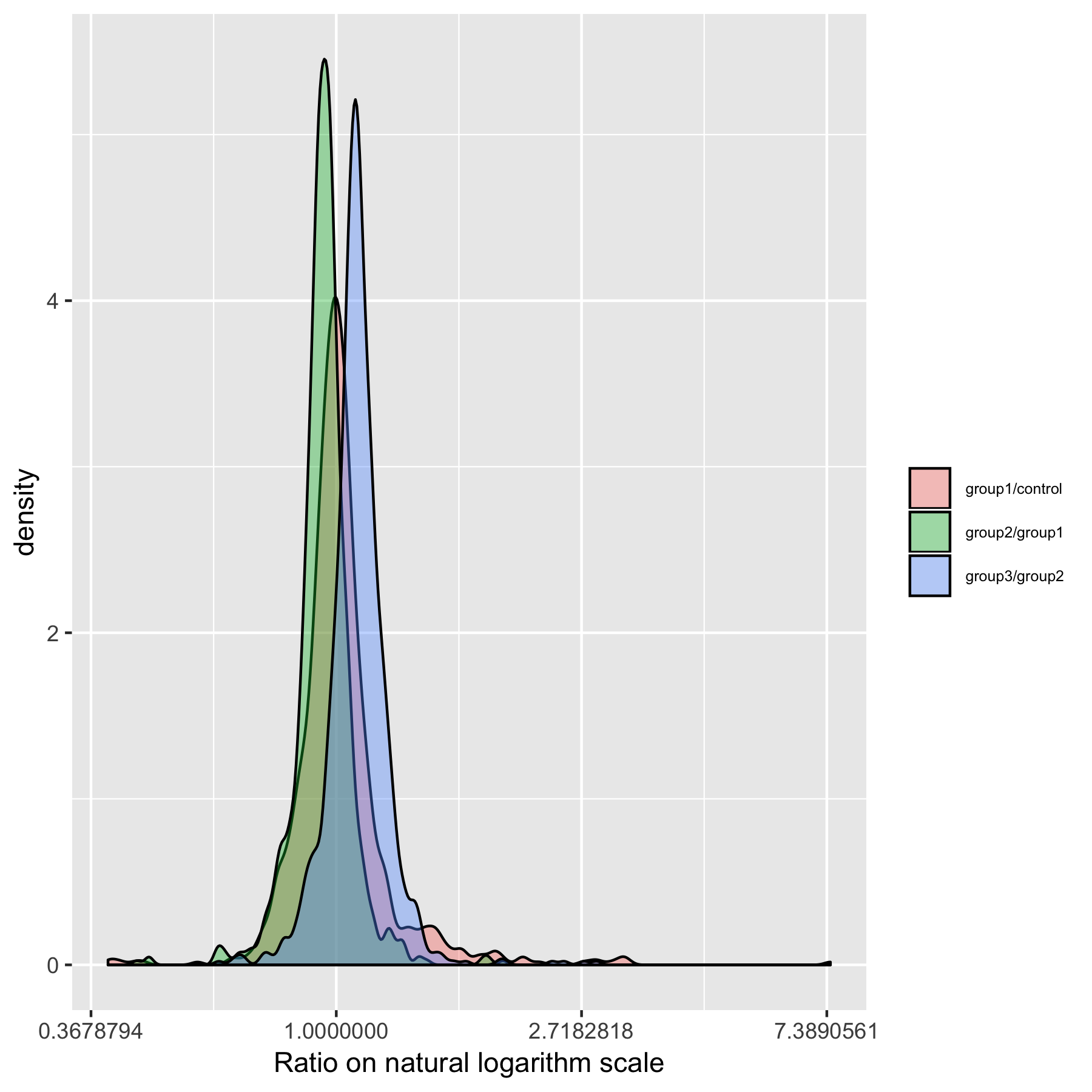

Figure S29. Ratios of group averages distribution from Dataset 4.

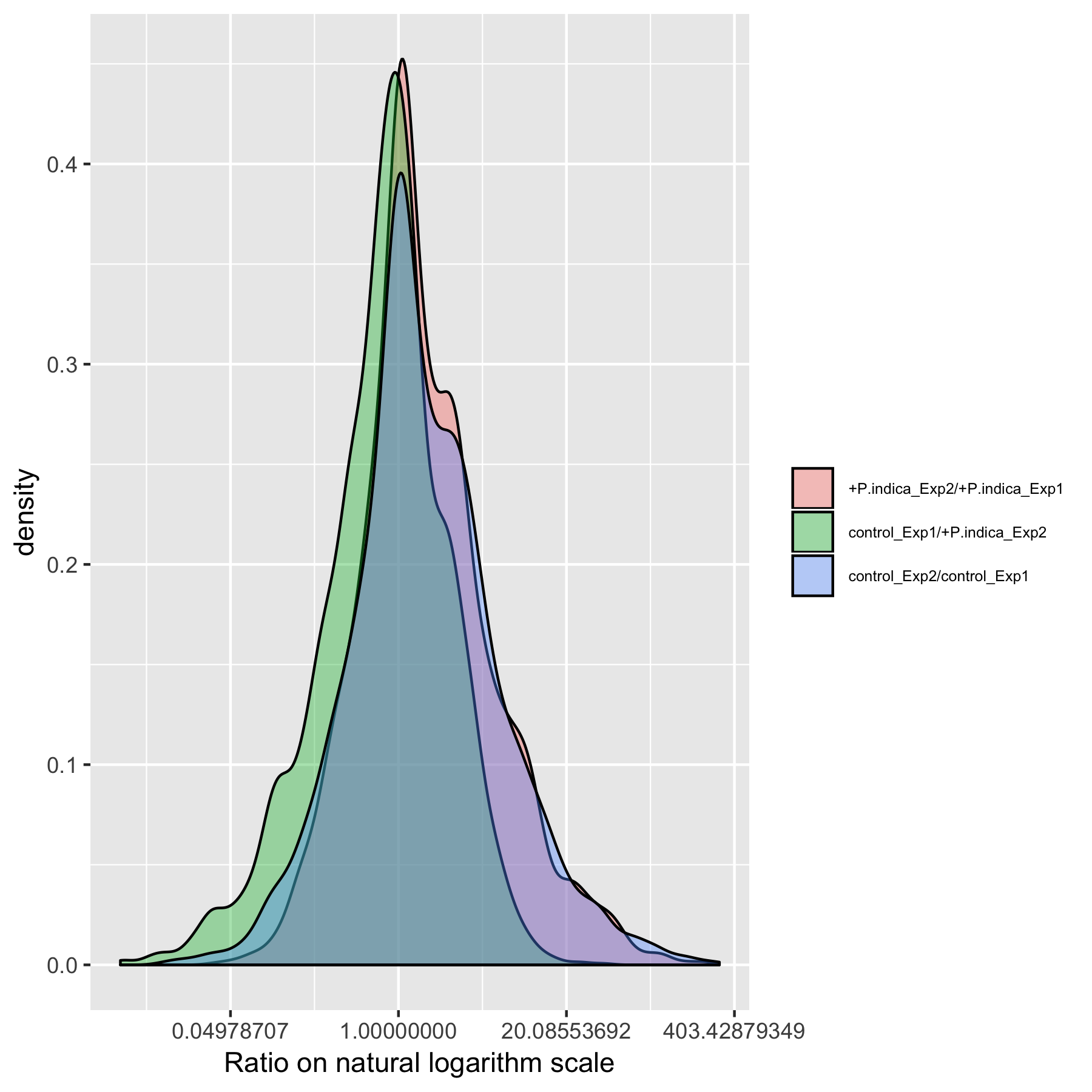

Figure S30. Ratios of group averages distribution from Dataset 5.

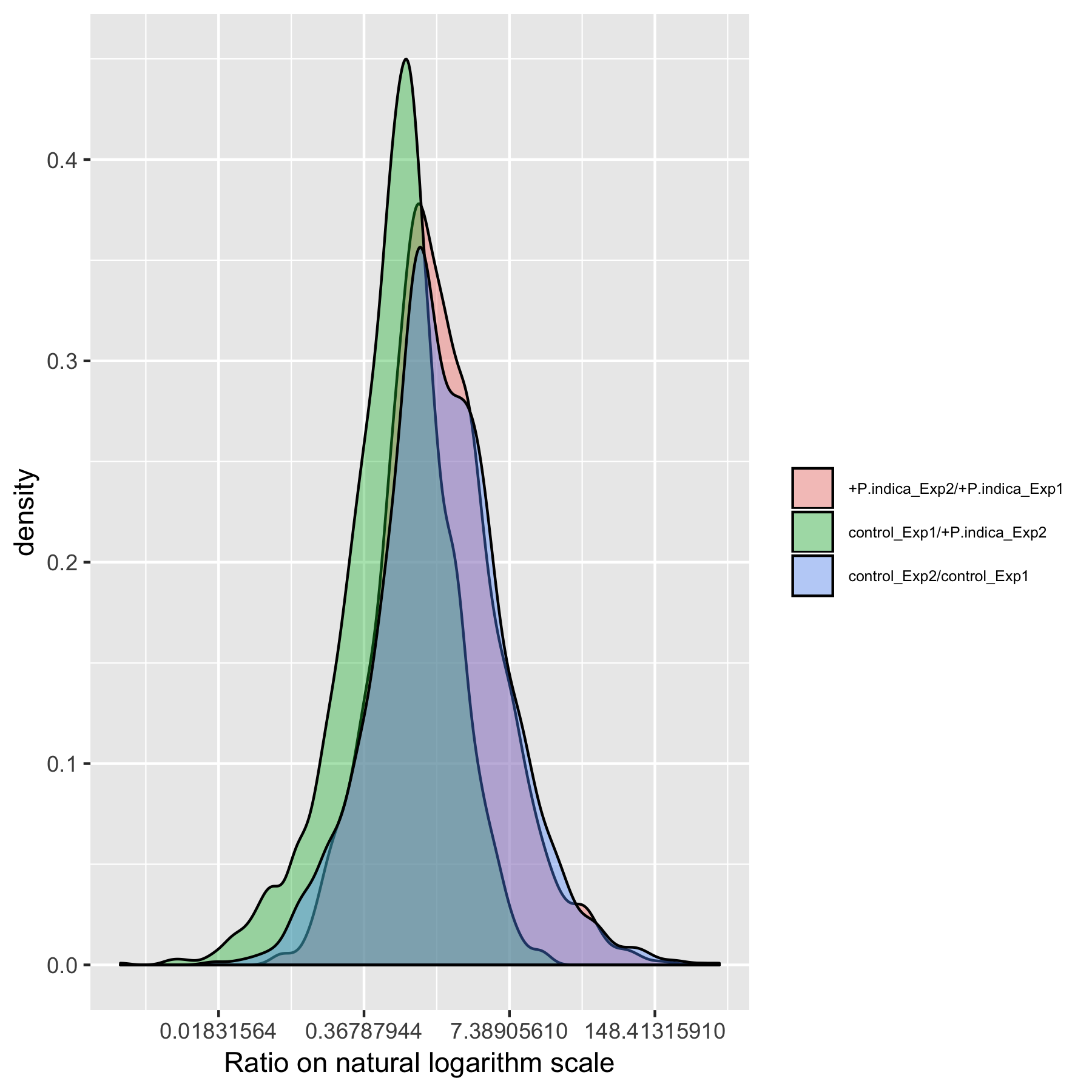

Figure S31. Ratios of group averages distribution from Dataset 6.

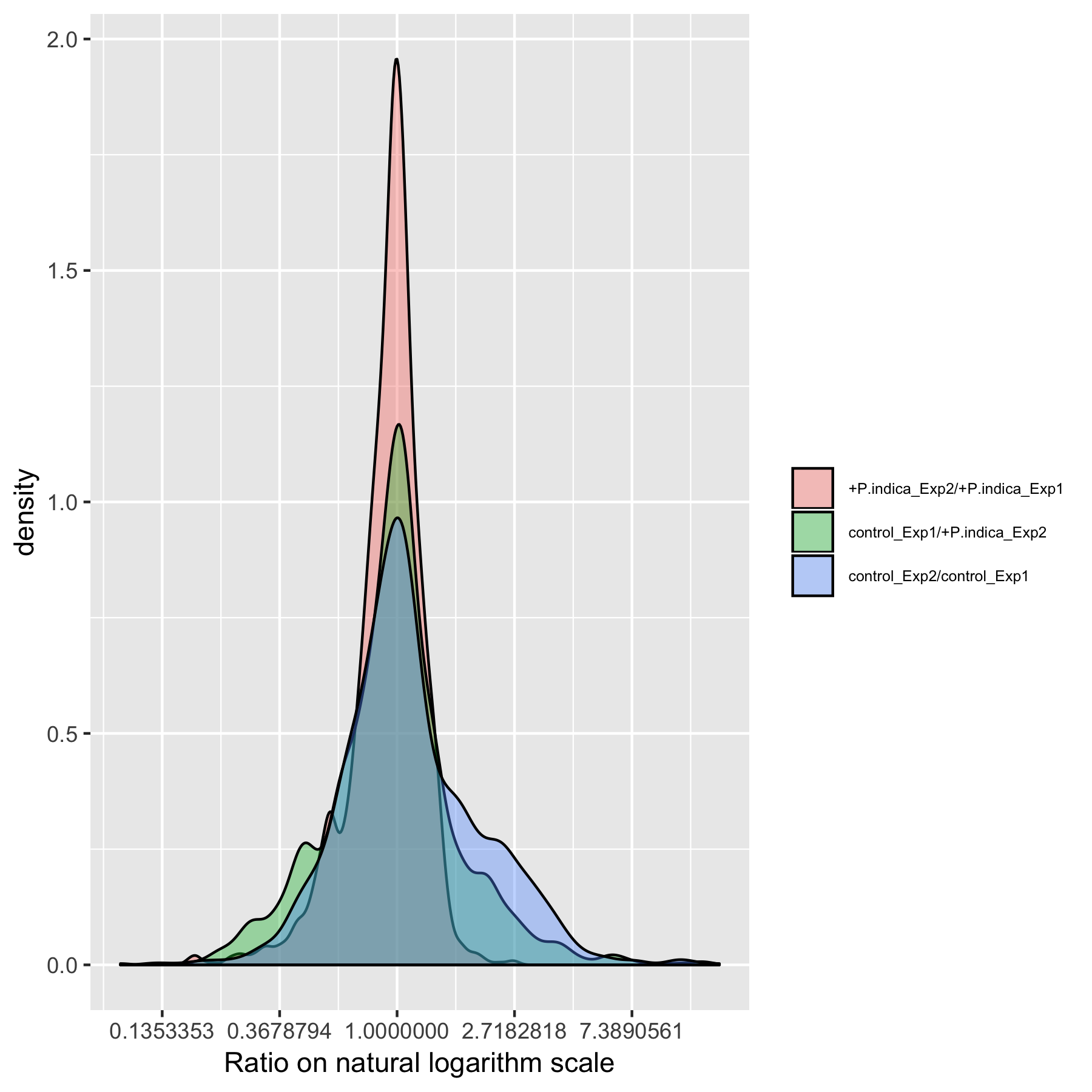

Figure S32. Ratios of group averages distribution from Dataset 7.

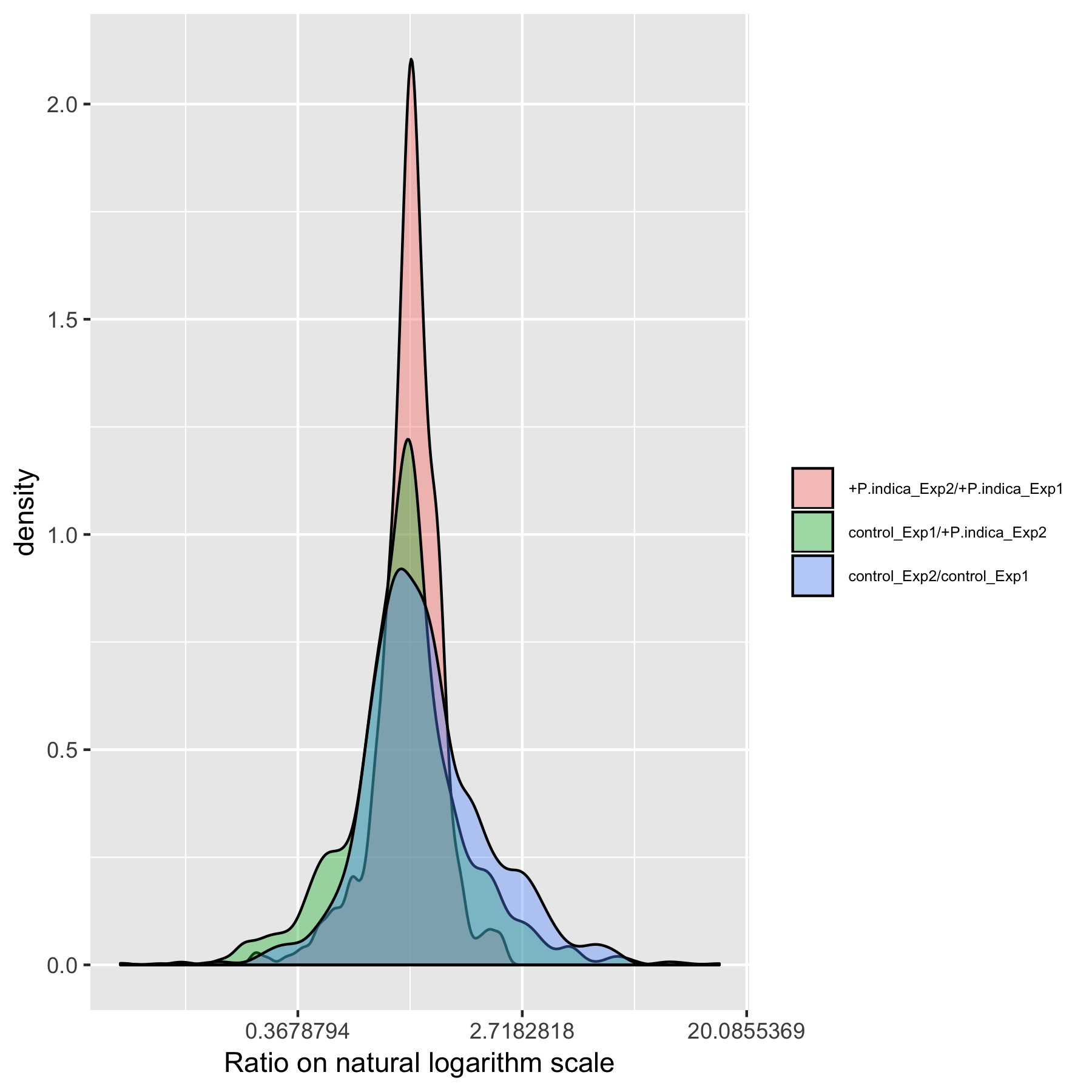

Figure S33. Ratios of group averages distribution from Dataset 8.

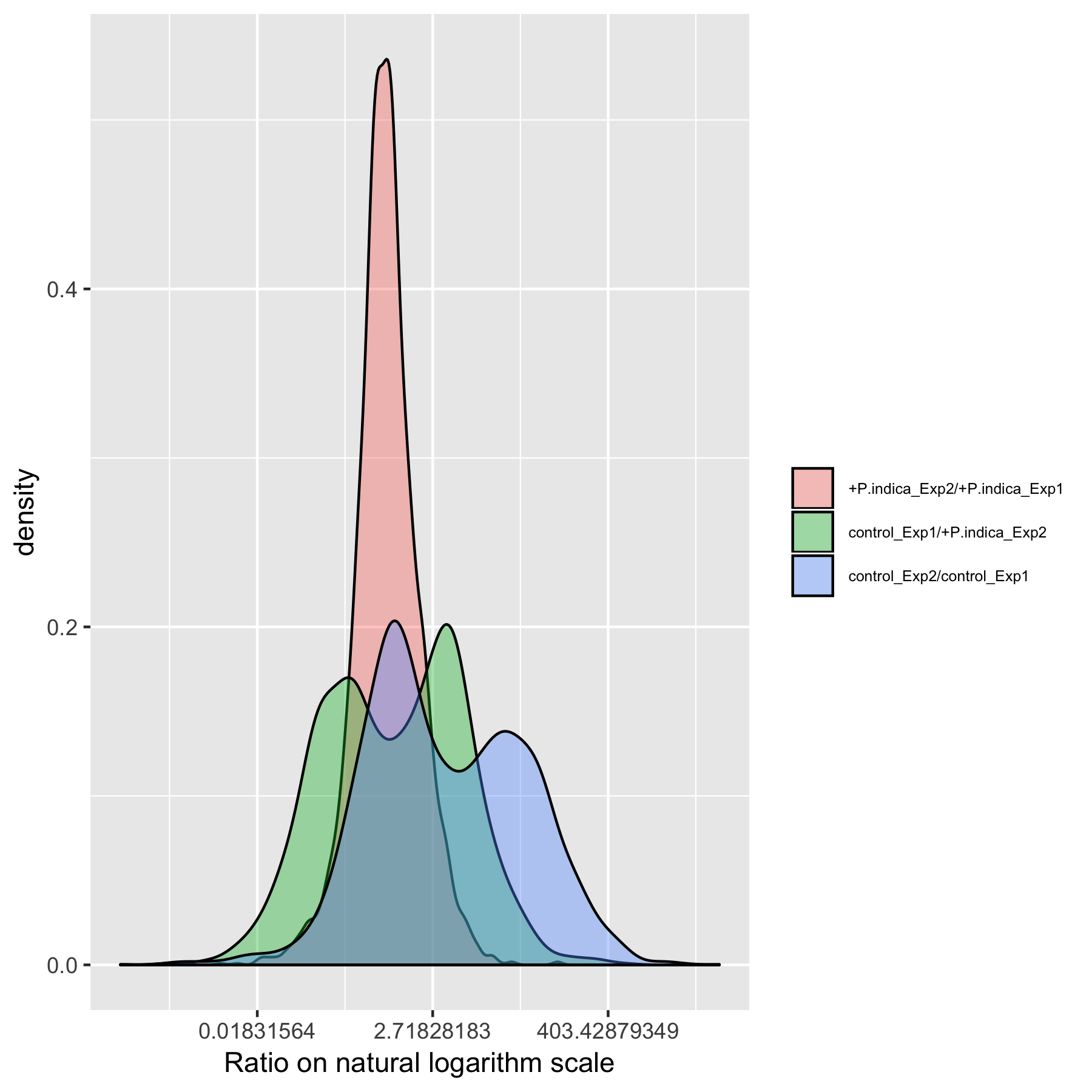

Figure S34. Ratios of group averages distribution from Dataset 9.

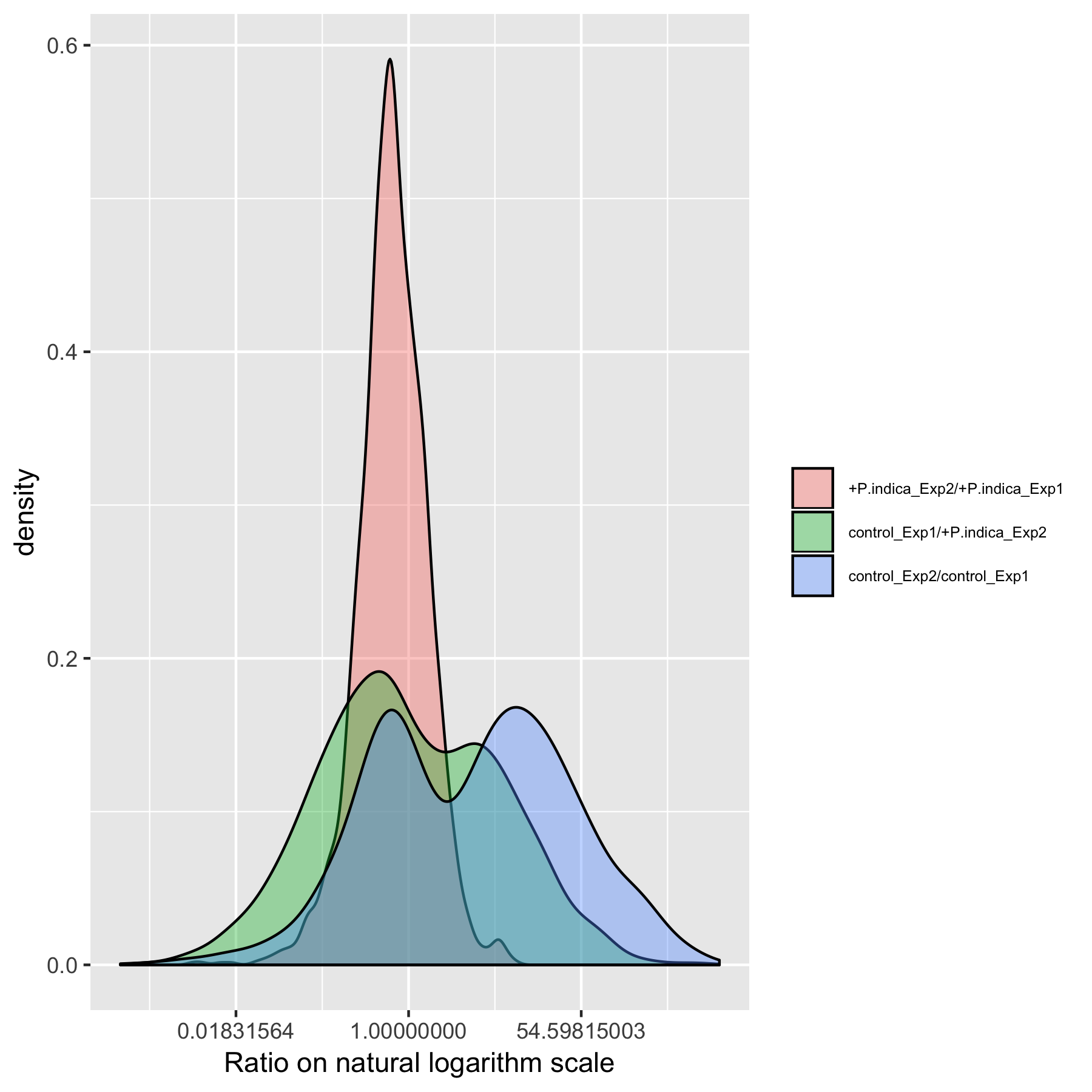

Figure S35. Ratios of group averages distribution from Dataset 10.

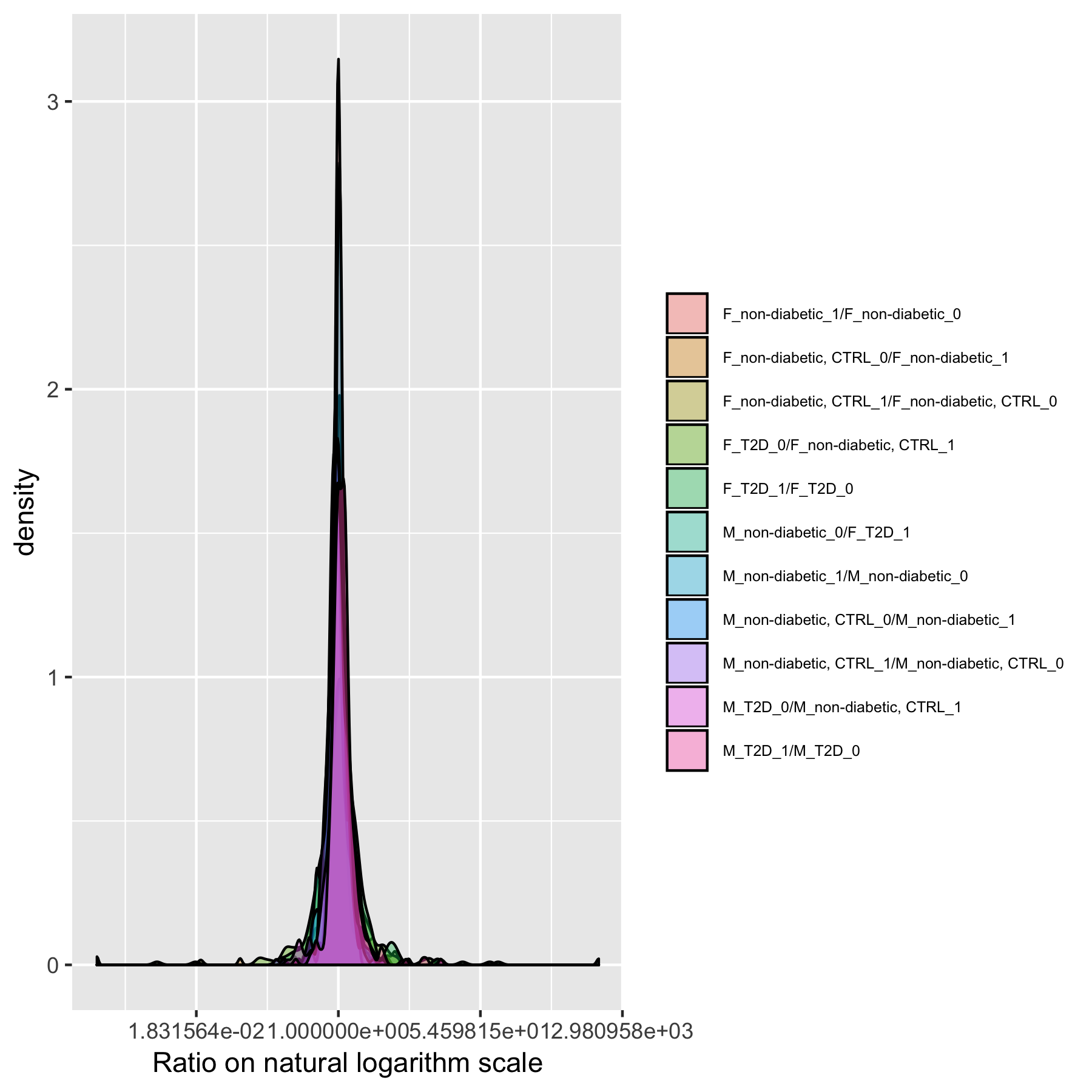

Figure S36. Ratios of group averages distribution from Dataset 11.

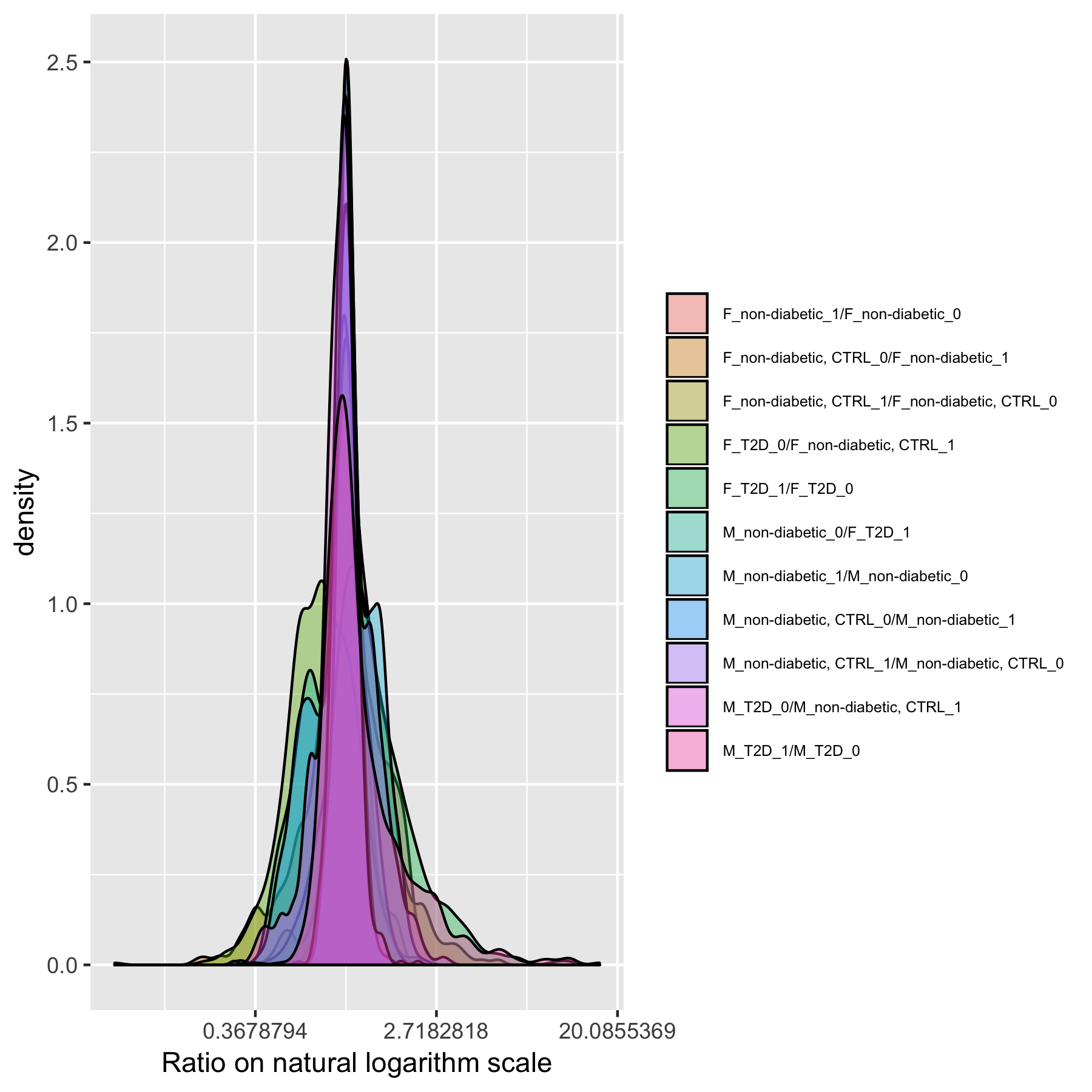

Figure S37. Ratios of group averages distribution from Dataset 12.

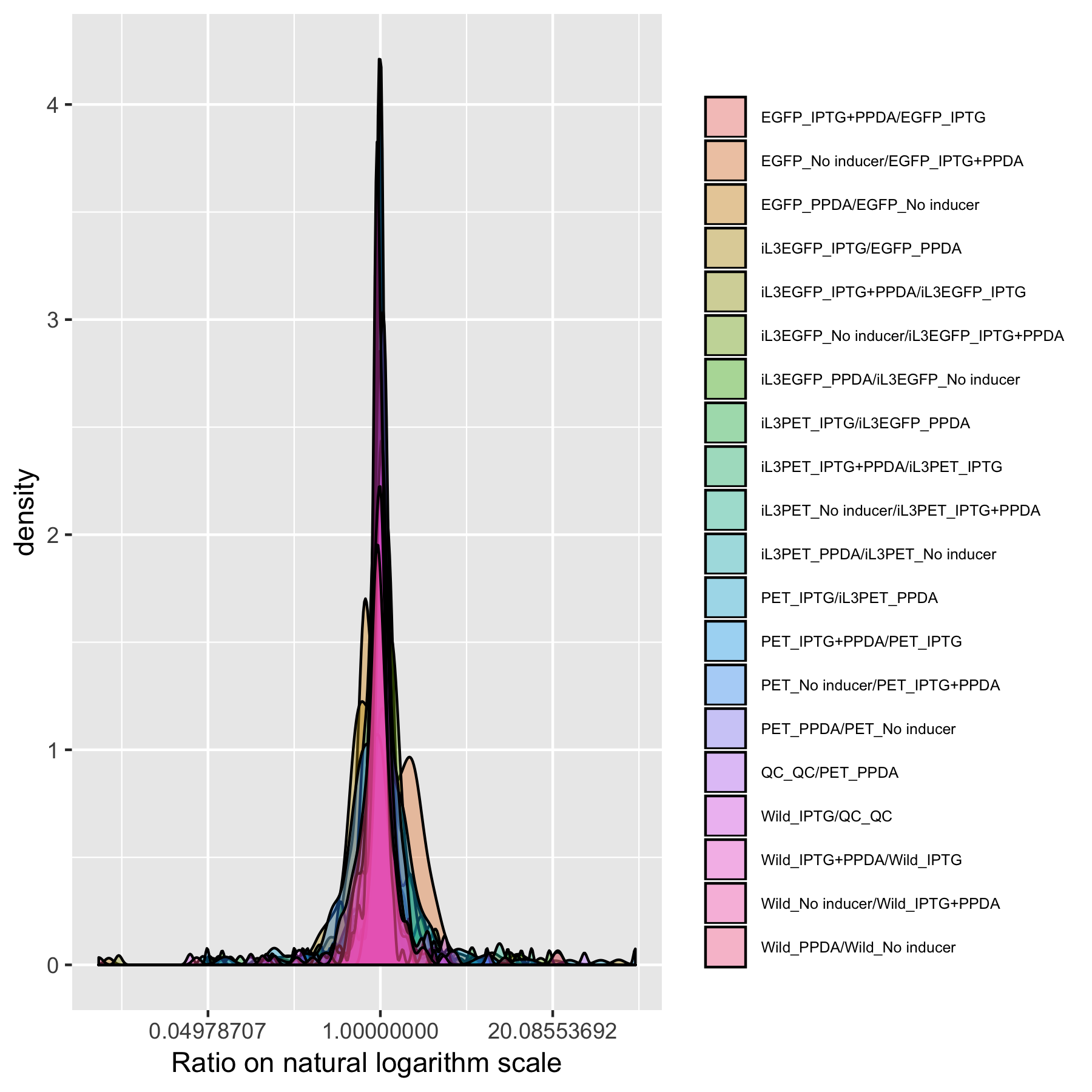

Figure S38. Ratios of group averages distribution from Dataset 13.

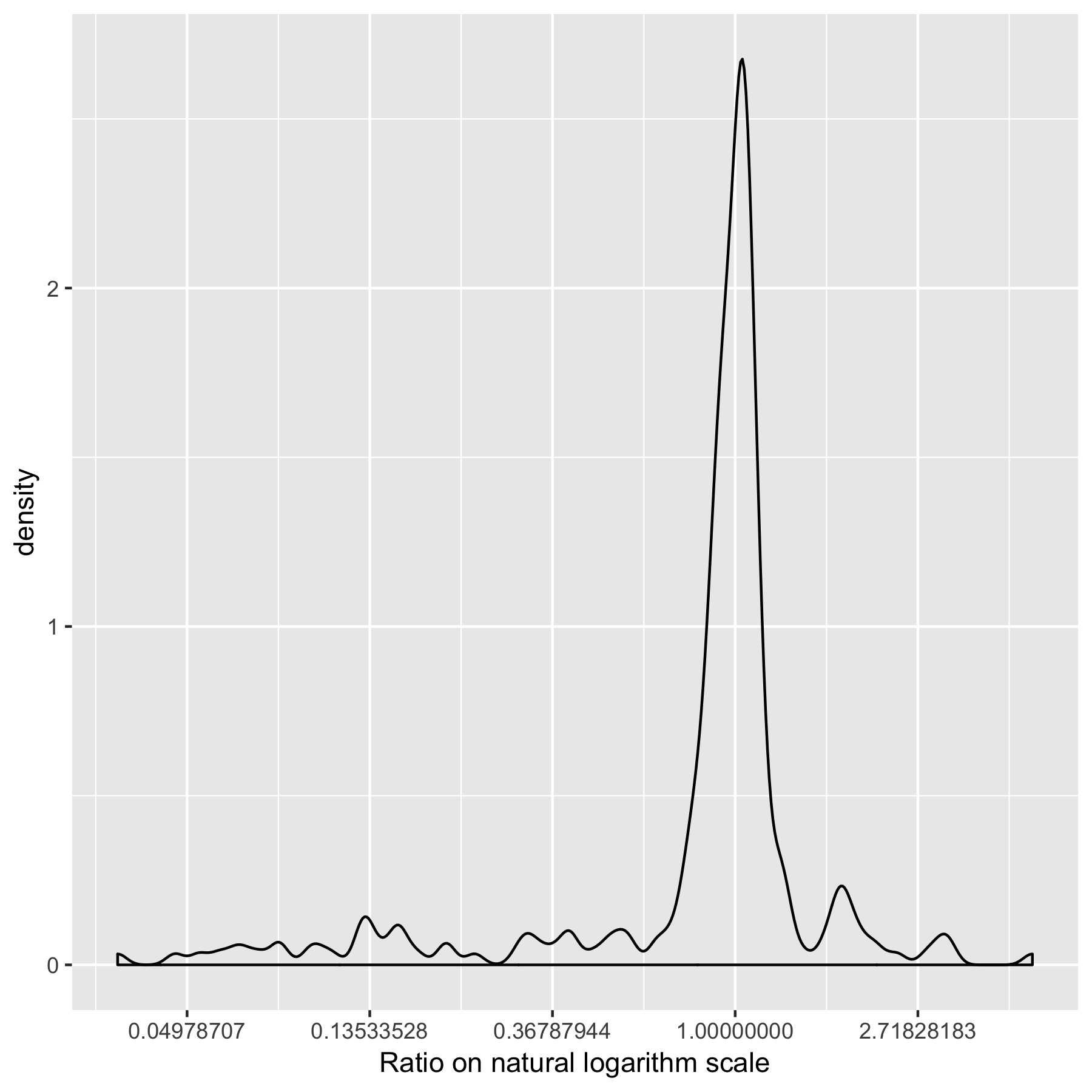

Figure S39. Ratios of group averages distribution from Dataset 14.

R Code for this research

\## ----setup,echo=FALSE,message=FALSE,warning=FALSE------------------------

knitr::opts_chunk$set(echo = F,message=FALSE,warning=FALSE,cache = T)

library(tidyverse)

library(reshape2)

library(scales)

library(fitdistrplus)

library(mzrtsim)

library(readxl)

library(mnormt)

library(readr)

library(reshape2)

\## ------------------------------------------------------------------------

\# import data

pos <- read_tsv('data/MTBLS28/m_mtbls28_POS_v2_maf.tsv')

anno <- read_tsv("data/MTBLS28/s_Diagnostic.txt")

\# find the pos data

posdata <- pos %>%

dplyr::select(contains('POS'))

\# find the mzrt

posmzrt <- pos %>%

dplyr::select(matches("mass|retention"))

\# find the pos sample class

posanno <- anno %>%

filter(grepl("POS",`Sample Name`))

\# check the order

identical(colnames(posdata),posanno$`Sample Name`)

\# get the class

posclass <- posanno %>%

transmute(class = paste(`Factor Value[Smoking]`,`Factor Value[Race]`,`Factor Value[Gender]`,`Factor Value[Sample Type]`,sep = '_')) %>%

as_factor() %>%

c() %>%

unlist()

\# get the data mean by group

data <- posdata %>%

t() %>%

as.tibble() %>%

mutate(class = posclass) %>%

group_by(class) %>%

summarise_all(mean) %>%

ungroup() %>%

t()

\# get the data sd by group

datasd <- posdata %>%

t() %>%

as.tibble() %>%

mutate(class = posclass) %>%

group_by(class) %>%

summarise_all(sd) %>%

ungroup() %>%

t()

colnames(data) <- data[1,]

dataclass <- data[-1,] %>%

apply(2, as.numeric)

colnames(datasd) <- datasd[1,]

dataclasssd <- datasd[-1,] %>%

apply(2, as.numeric)

name <- c('CSAAFC','CSAAFT','CSAAMC','CSAAMT','CSAAFC','CSAAFT','CSEAMC','CSEAMT','FSAAFC','FSAAFT','FSAAMC','FSAAMT','FSEAFC','FSEAFT','FSEAMC','FSEAMT','NSAAFC','NSAAFT','NSAAMC', 'NSAAMT', 'NSEAFC','NSEAFT','NSEAMC','NSEAFT')

colnames(dataclass) <- name

\# get the ratio

ratio <- dataclass[,-1] / dataclass[,-ncol(dataclass)]

colnames(ratio) <- paste(colnames(dataclass)[-1], colnames(dataclass)[-ncol(dataclass)], sep = "/")

dataratio<- melt(ratio)

fig1 <- ggplot(dataratio,aes(x = value, fill = Var2))+geom_density(alpha=0.382) +

xlab('Ratio on natural logarithm scale') +

scale_x_continuous(trans = log_trans()) +

theme(legend.position="right",legend.text=element_text(size=6))+ labs(fill="")

ggsave(fig1,filename = "fig1.png",width = 6,height = 6)

\# get the rsd within each group

library(reshape2)

datamean <- melt(dataclass)

datasd <- melt(dataclasssd)

datamean$rsd <- datamean$value/datasd$value*100

\# plot the intensity

fig2 <- ggplot(datamean,aes(x = value, fill=Var2)) +

geom_density(alpha=0.382) +

xlab('Intensity on Log scale') +

scale_x_log10() +

theme(legend.position="right",legend.text=element_text(size=6))+ labs(fill="")

ggsave(fig2,filename = "fig2.png",width = 6,height = 6)

\# plot the rsd

fig3 <- ggplot(datamean,aes(x = rsd, fill=Var2)) +

geom_density(alpha=0.382) +

xlab('RSD%') +

theme(legend.position="right",legend.text=element_text(size=6))+ labs(fill="")

ggsave(fig3,filename = "fig3.png",width = 6,height = 6)

\# plot the intensity

fig4 <- ggplot(datamean,aes(x = value, fill=Var2)) +

geom_density(alpha=0.382) +

xlab('Intensity') +

theme(legend.position="right",legend.text=element_text(size=6))+labs(fill="")

ggsave(fig4,filename = "fig4.png",width = 6,height = 6)

\# plot the distribution

descdist(c(log(datamean$value+1)),discrete = F)

descdist(c(datamean$rsd),discrete = F)

descdist(c(dataratio$value),discrete = F)

\## ----neg-----------------------------------------------------------------

\# import data

neg <- read_tsv('data/MTBLS28/m_mtbls28_NEG_v2_maf.tsv')

\# find the pos data

negdata <- neg %>%

dplyr::select(contains('NEG'))

\# find the mzrt

negmzrt <- neg %>%

dplyr::select(matches("mass|retention"))

\# find the pos sample class

neganno <- anno %>%

filter(grepl("NEG",`Sample Name`))

\# check the order

identical(colnames(negdata),neganno$`Sample Name`)

\# get the class

negclass <- neganno %>%

transmute(class = paste(`Factor Value[Smoking]`,`Factor Value[Race]`,`Factor Value[Gender]`,`Factor Value[Sample Type]`,sep = '_')) %>%

as_factor() %>%

c() %>%

unlist()

\# get the data mean by group

data <- negdata %>%

t() %>%

as.tibble() %>%

mutate(class = negclass) %>%

group_by(class) %>%

summarise_all(mean) %>%

ungroup() %>%

t()

\# get the data sd by group

datasd <- negdata %>%

t() %>%

as.tibble() %>%

mutate(class = posclass) %>%

group_by(class) %>%

summarise_all(sd) %>%

ungroup() %>%

t()

colnames(data) <- data[1,]

dataclass <- data[-1,] %>%

apply(2, as.numeric)

colnames(datasd) <- datasd[1,]

dataclasssd <- datasd[-1,] %>%

apply(2, as.numeric)

name <- c('CSAAFC','CSAAFT','CSAAMC','CSAAMT','CSAAFC','CSAAFT','CSEAMC','CSEAMT','FSAAFC','FSAAFT','FSAAMC','FSAAMT','FSEAFC','FSEAFT','FSEAMC','FSEAMT','NSAAFC','NSAAFT','NSAAMC', 'NSAAMT', 'NSEAFC','NSEAFT','NSEAMC','NSEAFT')

colnames(dataclass) <- name

\# get the ratio

ratio <- dataclass[,-1] / dataclass[,-ncol(dataclass)]

colnames(ratio) <- paste(colnames(dataclass)[-1], colnames(dataclass)[-ncol(dataclass)], sep = "/")

dataratio<- melt(ratio)

fig5 <- ggplot(dataratio,aes(x = value, fill = Var2))+geom_density(alpha=0.382) +

xlab('Ratio on natural logarithm scale') +

scale_x_continuous(trans = log_trans()) +

theme(legend.position="right",legend.text=element_text(size=6))+ labs(fill="")

ggsave(fig5,filename = "fig5.png",width = 6,height = 6)

\# get the rsd within each group

datamean<- melt(dataclass)

datasd <- melt(dataclasssd)

datamean$rsd <- datamean$value/datasd$value*100

\# plot the intensity

fig6 <- ggplot(datamean,aes(x = value, fill=Var2)) +

geom_density(alpha=0.382) +

xlab('Intensity on Log scale') +

scale_x_log10() +

theme(legend.position="right",legend.text=element_text(size=6))+ labs(fill="")

ggsave(fig6,filename = "fig6.png",width = 6,height = 6)

\# plot the rsd

fig7 <- ggplot(datamean,aes(x = rsd, fill=Var2)) +

geom_density(alpha=0.382) +

xlab('RSD%') +

theme(legend.position="right",legend.text=element_text(size=6))+ labs(fill="")

ggsave(fig7,filename = "fig7.png",width = 6,height = 6)

\# plot the intensity

fig8 <- ggplot(datamean,aes(x = value, fill=Var2)) +

geom_density(alpha=0.382) +

xlab('Intensity') +

theme(legend.position="right",legend.text=element_text(size=6))+labs(fill="")

ggsave(fig8,filename = "fig8.png",width = 6,height = 6)

\# plot the distribution

descdist(c(log(datamean$value+1)),discrete = F)

descdist(c(datamean$rsd),discrete = F)

descdist(c(dataratio$value),discrete = F)

\## ------------------------------------------------------------------------

library(BioMark)

\# import data

data(SpikePos)

data(SpikeNeg)

posdata <- t(SpikePos$data)

negdata <- t(SpikeNeg$data)

\# find the sample class

anno <- SpikePos$classes

posmzrt <- SpikePos$annotation

negmzrt <- SpikeNeg$annotation

list <- NULL

list$mz <- posmzrt$mz

list$rt <- posmzrt$rt

list$data <- posdata

\# get the data mean by group

data <- posdata %>%

t() %>%

as.tibble() %>%

mutate(class = anno) %>%

group_by(class) %>%

summarise_all(mean) %>%

ungroup() %>%

t()

\# get the data sd by group

datasd <- posdata %>%

t() %>%

as.tibble() %>%

mutate(class = anno) %>%

group_by(class) %>%

summarise_all(sd) %>%

ungroup() %>%

t()

colnames(data) <- data[1,]

dataclass <- data[-1,] %>%

apply(2, as.numeric)

colnames(datasd) <- datasd[1,]

dataclasssd <- datasd[-1,] %>%

apply(2, as.numeric)

\# get the ratio

ratio <- dataclass[,-1] / dataclass[,-ncol(dataclass)]

colnames(ratio) <- paste(colnames(dataclass)[-1], colnames(dataclass)[-ncol(dataclass)], sep = "/")

dataratio<- melt(ratio)

fig9 <- ggplot(dataratio,aes(x = value, fill = Var2))+geom_density(alpha=0.382) +

xlab('Ratio on natural logarithm scale') +

scale_x_continuous(trans = log_trans()) +

theme(legend.position="right",legend.text=element_text(size=6))+ labs(fill="")

ggsave(fig9,filename = "fig9.png",width = 6,height = 6)

\# get the rsd within each group

library(reshape2)

datamean<- melt(dataclass)

datasd <- melt(dataclasssd)

datamean$rsd <- datamean$value/datasd$value*100

\# plot the intensity

fig10 <- ggplot(datamean,aes(x = value, fill=Var2)) +

geom_density(alpha=0.382) +

xlab('Intensity on Log scale') +

scale_x_log10() +

theme(legend.position="top",legend.text=element_text(size=6))+ labs(fill="")

ggsave(fig10,filename = "fig10.png",width = 6,height = 6)

\# plot the rsd

fig11 <- ggplot(datamean,aes(x = rsd, fill=Var2)) +

geom_density(alpha=0.382) +

xlab('RSD%') +

theme(legend.position="top",legend.text=element_text(size=6))+ labs(fill="")

ggsave(fig11,filename = "fig11.png",width = 6,height = 6)

\# plot the intensity

fig12 <- ggplot(datamean,aes(x = value, fill=Var2)) +

geom_density(alpha=0.382) +

xlab('Intensity') +

theme(legend.position="top",legend.text=element_text(size=6))+labs(fill="")

ggsave(fig12,filename = "fig12.png",width = 6,height = 6)

\# plot the distribution

descdist(c(log(datamean$value+1)),discrete = F)

descdist(c(datamean$rsd),discrete = F)

descdist(c(dataratio$value),discrete = F)

\# for negative mode

\# get the data mean by group

data <- negdata %>%

t() %>%

as.tibble() %>%

mutate(class = anno) %>%

group_by(class) %>%

summarise_all(mean) %>%

ungroup() %>%

t()

\# get the data sd by group

datasd <- negdata %>%

t() %>%

as.tibble() %>%

mutate(class = anno) %>%

group_by(class) %>%

summarise_all(sd) %>%

ungroup() %>%

t()

colnames(data) <- data[1,]

dataclass <- data[-1,] %>%

apply(2, as.numeric)

\# get the ratio

ratio <- dataclass[,-1] / dataclass[,-ncol(dataclass)]

colnames(ratio) <- paste(colnames(dataclass)[-1], colnames(dataclass)[-ncol(dataclass)], sep = "/")

dataratio<- melt(ratio)

fig13 <- ggplot(dataratio,aes(x = value, fill = Var2))+geom_density(alpha=0.382) +

xlab('Ratio on natural logarithm scale') +

scale_x_continuous(trans = log_trans()) +

theme(legend.position="right",legend.text=element_text(size=6))+ labs(fill="")

ggsave(fig13,filename = "fig13.png",width = 6,height = 6)

colnames(datasd) <- datasd[1,]

dataclasssd <- datasd[-1,] %>%

apply(2, as.numeric)

\# get the rsd within each group

library(reshape2)

datamean<- melt(dataclass)

datasd <- melt(dataclasssd)

datamean$rsd <- datamean$value/datasd$value*100

\# plot the intensity

fig14 <- ggplot(datamean,aes(x = value, fill=Var2)) +

geom_density(alpha=0.382) +

xlab('Intensity on Log scale') +

scale_x_log10() +

theme(legend.position="right",legend.text=element_text(size=6))+ labs(fill="")

ggsave(fig14,filename = "fig14.png",width = 6,height = 6)

\# plot the rsd

fig15 <- ggplot(datamean,aes(x = rsd, fill=Var2)) +

geom_density(alpha=0.382) +

xlab('RSD%') +

theme(legend.position="right",legend.text=element_text(size=6))+ labs(fill="")

ggsave(fig15,filename = "fig15.png",width = 6,height = 6)

\# plot the intensity

fig16 <- ggplot(datamean,aes(x = value, fill=Var2)) +

geom_density(alpha=0.382) +

xlab('Intensity') +

theme(legend.position="right",legend.text=element_text(size=6))+labs(fill="")

ggsave(fig16,filename = "fig16.png",width = 6,height = 6)

\# plot the distribution

descdist(c(log(datamean$value+1)),discrete = F)

descdist(c(datamean$rsd),discrete = F)

descdist(c(dataratio$value),discrete = F)

\## ------------------------------------------------------------------------

\# import data

root <- read_tsv('data/MTBLS341/m_mtbls341_metabolite_profiling_lcms-root_pos_v2_maf.tsv')

rootneg <- read_tsv('data/MTBLS341/m_mtbls341_metabolite_profiling_lcms-root_neg_v2_maf.tsv')

leaf <- read_tsv('data/MTBLS341/m_mtbls341_metabolite_profiling_lcms-leaf_pos_v2_maf.tsv')

leafneg <- read_tsv('data/MTBLS341/m_mtbls341_metabolite_profiling_lcms-leaf_neg_v2_maf.tsv')

exudate <- read_tsv('data/MTBLS341/m_mtbls341_metabolite_profiling_lcms-exudate_pos_v2_maf.tsv')

exudateneg <- read_tsv('data/MTBLS341/m_mtbls341_metabolite_profiling_lcms-exudate_neg_v2_maf.tsv')

anno <- read_tsv("data/MTBLS341/s_MTBLS341.txt")

pos <- root

\# find the pos data

posdata <- pos %>%

dplyr::select(contains('pos'))

\# find the mzrt

posmzrt <- pos %>%

dplyr::select(matches("mass|retention"))

\# find the pos sample class

posanno <- anno %>%

filter(grepl("root",`Characteristics[Organism part]`))

\# find the class

posclass <- posanno %>%

transmute(class = paste(`Factor Value[Treatment]`,`Factor Value[Replicate]`,sep = '_')) %>%

as_factor() %>%

c() %>%

unlist()

\# hack to add

posclass <- c(posclass[1:13],'control_Exp2','control_Exp2',posclass[14:18])

\# get the data mean by group

data <- posdata %>%

t() %>%

as.tibble() %>%

mutate(class = posclass) %>%

group_by(class) %>%

summarise_all(mean) %>%

ungroup() %>%

t()

\# get the data sd by group

datasd <- posdata %>%

t() %>%

as.tibble() %>%

mutate(class = posclass) %>%

group_by(class) %>%

summarise_all(sd) %>%

ungroup() %>%

t()

colnames(data) <- data[1,]

dataclass <- data[-1,] %>%

apply(2, as.numeric)

colnames(datasd) <- datasd[1,]

dataclasssd <- datasd[-1,] %>%

apply(2, as.numeric)

\# get the ratio

ratio <- dataclass[,-1] / dataclass[,-ncol(dataclass)]

colnames(ratio) <- paste(colnames(dataclass)[-1], colnames(dataclass)[-ncol(dataclass)], sep = "/")

dataratio<- melt(ratio)

fig17 <- ggplot(dataratio,aes(x = value, fill = Var2))+geom_density(alpha=0.382) +

xlab('Ratio on natural logarithm scale') +

scale_x_continuous(trans = log_trans()) +

theme(legend.position="right",legend.text=element_text(size=6))+ labs(fill="")

ggsave(fig17,filename = "fig17.png",width = 6,height = 6)

\# get the rsd within each group

datamean<- melt(dataclass)

datasd <- melt(dataclasssd)

datamean$rsd <- datamean$value/datasd$value*100

\# plot the intensity

fig18 <- ggplot(datamean,aes(x = value, fill=Var2)) +

geom_density(alpha=0.382) +

xlab('Intensity on Log scale') +

scale_x_log10() +

theme(legend.position="top",legend.text=element_text(size=6))+ labs(fill="")

ggsave(fig18,filename = "fig18.png",width = 6,height = 6)

\# plot the rsd

fig19 <- ggplot(datamean,aes(x = rsd, fill=Var2)) +

geom_density(alpha=0.382) +

xlab('RSD%') +

theme(legend.position="top",legend.text=element_text(size=6))+ labs(fill="")

ggsave(fig19,filename = "fig19.png",width = 6,height = 6)

\# plot the intensity

fig20 <- ggplot(datamean,aes(x = value, fill=Var2)) +

geom_density(alpha=0.382) +

xlab('Intensity') +

theme(legend.position="top",legend.text=element_text(size=6))+labs(fill="")

ggsave(fig20,filename = "fig20.png",width = 6,height = 6)

\# plot the distribution

descdist(c(log(datamean$value+1)),discrete = F)

descdist(c(na.omit(datamean$rsd)),discrete = F)

descdist(dataratio$value[!is.infinite(dataratio$value)],discrete = F)

pos <- rootneg

\# find the pos data

posdata <- pos %>%

dplyr::select(contains('neg'))

\# find the mzrt

posmzrt <- pos %>%

dplyr::select(matches("mass|retention"))

\# find the pos sample class

posanno <- anno %>%

filter(grepl("root",`Characteristics[Organism part]`))

\# find the class

posclass <- posanno %>%

transmute(class = paste(`Factor Value[Treatment]`,`Factor Value[Replicate]`,sep = '_')) %>%

as_factor() %>%

c() %>%

unlist()

\# hack to add

posclass <- c(posclass[1:13],'control_Exp2','control_Exp2',posclass[14:18])

\# get the data mean by group

data <- posdata %>%

t() %>%

as.tibble() %>%

mutate(class = posclass) %>%

group_by(class) %>%

summarise_all(mean) %>%

ungroup() %>%

t()

\# get the data sd by group

datasd <- posdata %>%

t() %>%

as.tibble() %>%

mutate(class = posclass) %>%

group_by(class) %>%

summarise_all(sd) %>%

ungroup() %>%

t()

colnames(data) <- data[1,]

dataclass <- data[-1,] %>%

apply(2, as.numeric)

colnames(datasd) <- datasd[1,]

dataclasssd <- datasd[-1,] %>%

apply(2, as.numeric)

\# get the ratio

ratio <- dataclass[,-1] / dataclass[,-ncol(dataclass)]

colnames(ratio) <- paste(colnames(dataclass)[-1], colnames(dataclass)[-ncol(dataclass)], sep = "/")

dataratio<- melt(ratio)

fig21 <- ggplot(dataratio,aes(x = value, fill = Var2))+geom_density(alpha=0.382) +

xlab('Ratio on natural logarithm scale') +

scale_x_continuous(trans = log_trans()) +

theme(legend.position="right",legend.text=element_text(size=6))+ labs(fill="")

ggsave(fig21,filename = "fig21.png",width = 6,height = 6)

\# get the rsd within each group

datamean<- melt(dataclass)

datasd <- melt(dataclasssd)

datamean$rsd <- datamean$value/datasd$value*100

\# plot the intensity

fig22 <- ggplot(datamean,aes(x = value, fill=Var2)) +

geom_density(alpha=0.382) +

xlab('Intensity on Log scale') +

scale_x_log10() +

theme(legend.position="top",legend.text=element_text(size=6))+ labs(fill="")

ggsave(fig22,filename = "fig22.png",width = 6,height = 6)

\# plot the rsd

fig23 <- ggplot(datamean,aes(x = rsd, fill=Var2)) +

geom_density(alpha=0.382) +

xlab('RSD%') +

theme(legend.position="top",legend.text=element_text(size=6))+ labs(fill="")

ggsave(fig23,filename = "fig23.png",width = 6,height = 6)

\# plot the intensity

fig24 <- ggplot(datamean,aes(x = value, fill=Var2)) +

geom_density(alpha=0.382) +

xlab('Intensity') +

theme(legend.position="top",legend.text=element_text(size=6))+labs(fill="")

ggsave(fig24,filename = "fig24.png",width = 6,height = 6)

\# plot the distribution

descdist(c(log(datamean$value+1)),discrete = F)

descdist(c(datamean$rsd),discrete = F)

descdist(c(dataratio$value),discrete = F)

pos <- leaf

\# find the pos data

posdata <- pos %>%

dplyr::select(contains('pos'))

\# find the mzrt

posmzrt <- pos %>%

dplyr::select(matches("mass|retention"))

\# find the pos sample class

posanno <- anno %>%

filter(grepl("leaf",`Characteristics[Organism part]`))

\# find the class

posclass <- posanno %>%

transmute(class = paste(`Factor Value[Treatment]`,`Factor Value[Replicate]`,sep = '_')) %>%

as_factor() %>%

c() %>%

unlist()

\# get the data mean by group

data <- posdata %>%

t() %>%

as.tibble() %>%

mutate(class = posclass) %>%

group_by(class) %>%

summarise_all(mean) %>%

ungroup() %>%

t()

\# get the data sd by group

datasd <- posdata %>%

t() %>%

as.tibble() %>%

mutate(class = posclass) %>%

group_by(class) %>%

summarise_all(sd) %>%

ungroup() %>%

t()

colnames(data) <- data[1,]

dataclass <- data[-1,] %>%

apply(2, as.numeric)

colnames(datasd) <- datasd[1,]

dataclasssd <- datasd[-1,] %>%

apply(2, as.numeric)

\# get the ratio

ratio <- dataclass[,-1] / dataclass[,-ncol(dataclass)]

colnames(ratio) <- paste(colnames(dataclass)[-1], colnames(dataclass)[-ncol(dataclass)], sep = "/")

dataratio<- melt(ratio)

fig25 <- ggplot(dataratio,aes(x = value, fill = Var2))+geom_density(alpha=0.382) +

xlab('Ratio on natural logarithm scale') +

scale_x_continuous(trans = log_trans()) +

theme(legend.position="right",legend.text=element_text(size=6))+ labs(fill="")

ggsave(fig25,filename = "fig25.png",width = 6,height = 6)

\# get the rsd within each group

datamean<- melt(dataclass)

datasd <- melt(dataclasssd)

datamean$rsd <- datamean$value/datasd$value*100

\# plot the intensity

fig26 <- ggplot(datamean,aes(x = value, fill=Var2)) +

geom_density(alpha=0.382) +

xlab('Intensity on Log scale') +

scale_x_log10() +

theme(legend.position="right",legend.text=element_text(size=6))+ labs(fill="")

ggsave(fig26,filename = "fig26.png",width = 6,height = 6)

\# plot the rsd

fig27 <- ggplot(datamean,aes(x = rsd, fill=Var2)) +

geom_density(alpha=0.382) +

xlab('RSD%') +

theme(legend.position="right",legend.text=element_text(size=6))+ labs(fill="")

ggsave(fig27,filename = "fig27.png",width = 6,height = 6)

\# plot the intensity

fig28 <- ggplot(datamean,aes(x = value, fill=Var2)) +

geom_density(alpha=0.382) +

xlab('Intensity') +

theme(legend.position="right",legend.text=element_text(size=6))+labs(fill="")

ggsave(fig28,filename = "fig28.png",width = 6,height = 6)

\# plot the distribution

descdist(c(log(datamean$value+1)),discrete = F)

descdist(c(datamean$rsd),discrete = F)

descdist(c(dataratio$value),discrete = F)

pos <- leafneg

\# find the pos data

posdata <- pos %>%

dplyr::select(contains('neg'))

\# find the mzrt

posmzrt <- pos %>%

dplyr::select(matches("mass|retention"))

\# find the pos sample class

posanno <- anno %>%

filter(grepl("leaf",`Characteristics[Organism part]`))

\# find the class

posclass <- posanno %>%

transmute(class = paste(`Factor Value[Treatment]`,`Factor Value[Replicate]`,sep = '_')) %>%

as_factor() %>%

c() %>%

unlist()

\# get the data mean by group

data <- posdata %>%

t() %>%

as.tibble() %>%

mutate(class = posclass) %>%

group_by(class) %>%

summarise_all(mean) %>%

ungroup() %>%

t()

\# get the data sd by group

datasd <- posdata %>%

t() %>%

as.tibble() %>%

mutate(class = posclass) %>%

group_by(class) %>%

summarise_all(sd) %>%

ungroup() %>%

t()

colnames(data) <- data[1,]

dataclass <- data[-1,] %>%

apply(2, as.numeric)

colnames(datasd) <- datasd[1,]

dataclasssd <- datasd[-1,] %>%

apply(2, as.numeric)

\# get the ratio

ratio <- dataclass[,-1] / dataclass[,-ncol(dataclass)]

colnames(ratio) <- paste(colnames(dataclass)[-1], colnames(dataclass)[-ncol(dataclass)], sep = "/")

dataratio<- melt(ratio)

fig29 <- ggplot(dataratio,aes(x = value, fill = Var2))+geom_density(alpha=0.382) +

xlab('Ratio on natural logarithm scale') +

scale_x_continuous(trans = log_trans()) +

theme(legend.position="right",legend.text=element_text(size=6))+ labs(fill="")

ggsave(fig29,filename = "fig29.png",width = 6,height = 6)

\# get the rsd within each group

datamean<- melt(dataclass)

datasd <- melt(dataclasssd)

datamean$rsd <- datamean$value/datasd$value*100

\# plot the intensity

fig30 <- ggplot(datamean,aes(x = value, fill=Var2)) +

geom_density(alpha=0.382) +

xlab('Intensity on Log scale') +

scale_x_log10() +

theme(legend.position="top",legend.text=element_text(size=6))+ labs(fill="")

ggsave(fig30,filename = "fig30.png",width = 6,height = 6)

\# plot the rsd

fig31 <- ggplot(datamean,aes(x = rsd, fill=Var2)) +

geom_density(alpha=0.382) +

xlab('RSD%') +

theme(legend.position="top",legend.text=element_text(size=6))+ labs(fill="")

ggsave(fig31,filename = "fig31.png",width = 6,height = 6)

\# plot the intensity

fig32 <- ggplot(datamean,aes(x = value, fill=Var2)) +

geom_density(alpha=0.382) +

xlab('Intensity') +

theme(legend.position="top",legend.text=element_text(size=6))+labs(fill="")

ggsave(fig32,filename = "fig32.png",width = 6,height = 6)

\# plot the distribution

descdist(c(log(datamean$value+1)),discrete = F)

descdist(c(datamean$rsd),discrete = F)

descdist(c(dataratio$value),discrete = F)

pos <- exudate

\# find the pos data

posdata <- pos %>%

dplyr::select(contains('pos'))

\# find the mzrt

posmzrt <- pos %>%

dplyr::select(matches("mass|retention"))

\# find the pos sample class

posanno <- anno %>%

filter(grepl("Exudate",`Characteristics[Organism part]`))

\# find the class

posclass <- posanno %>%

transmute(class = paste(`Factor Value[Treatment]`,`Factor Value[Replicate]`,sep = '_')) %>%

as_factor() %>%

c() %>%

unlist()

\# get the data mean by group

data <- posdata %>%

t() %>%

as.tibble() %>%

mutate(class = posclass) %>%

group_by(class) %>%

summarise_all(mean) %>%

ungroup() %>%

t()

\# get the data sd by group

datasd <- posdata %>%

t() %>%

as.tibble() %>%

mutate(class = posclass) %>%

group_by(class) %>%

summarise_all(sd) %>%

ungroup() %>%

t()

colnames(data) <- data[1,]

dataclass <- data[-1,] %>%

apply(2, as.numeric)

colnames(datasd) <- datasd[1,]

dataclasssd <- datasd[-1,] %>%

apply(2, as.numeric)

\# get the ratio

ratio <- dataclass[,-1] / dataclass[,-ncol(dataclass)]

colnames(ratio) <- paste(colnames(dataclass)[-1], colnames(dataclass)[-ncol(dataclass)], sep = "/")

dataratio<- melt(ratio)

fig33 <- ggplot(dataratio,aes(x = value, fill = Var2))+geom_density(alpha=0.382) +

xlab('Ratio on natural logarithm scale') +

scale_x_continuous(trans = log_trans()) +

theme(legend.position="right",legend.text=element_text(size=6))+ labs(fill="")

ggsave(fig33,filename = "fig33.png",width = 6,height = 6)

\# get the rsd within each group

datamean<- melt(dataclass)

datasd <- melt(dataclasssd)

datamean$rsd <- datamean$value/datasd$value*100

\# plot the intensity

fig34 <- ggplot(datamean,aes(x = value, fill=Var2)) +

geom_density(alpha=0.382) +

xlab('Intensity on Log scale') +

scale_x_log10() +

theme(legend.position="top",legend.text=element_text(size=6))+ labs(fill="")

ggsave(fig34,filename = "fig34.png",width = 6,height = 6)

\# plot the rsd

fig35 <- ggplot(datamean,aes(x = rsd, fill=Var2)) +

geom_density(alpha=0.382) +

xlab('RSD%') +

theme(legend.position="top",legend.text=element_text(size=6))+ labs(fill="")

ggsave(fig35,filename = "fig35.png",width = 6,height = 6)

\# plot the intensity

fig36 <- ggplot(datamean,aes(x = value, fill=Var2)) +

geom_density(alpha=0.382) +

xlab('Intensity') +

theme(legend.position="top",legend.text=element_text(size=6))+labs(fill="")

ggsave(fig36,filename = "fig36.png",width = 6,height = 6)

\# plot the distribution

descdist(c(log(datamean$value+1)),discrete = F)

descdist(c(datamean$rsd[!is.na(datamean$rsd)]),discrete = F)

descdist(c(dataratio$value[is.finite(dataratio$value)]),discrete = F)

pos <- exudateneg

\# find the pos data

posdata <- pos %>%

dplyr::select(contains('neg'))

\# find the mzrt

posmzrt <- pos %>%

dplyr::select(matches("mass|retention"))

\# find the pos sample class

posanno <- anno %>%

filter(grepl("Exudate",`Characteristics[Organism part]`))

\# find the class

posclass <- posanno %>%

transmute(class = paste(`Factor Value[Treatment]`,`Factor Value[Replicate]`,sep = '_')) %>%

as_factor() %>%

c() %>%

unlist()

\# get the data mean by group

data <- posdata %>%

t() %>%

as.tibble() %>%

mutate(class = posclass) %>%

group_by(class) %>%

summarise_all(mean) %>%

ungroup() %>%

t()

\# get the data sd by group

datasd <- posdata %>%

t() %>%

as.tibble() %>%

mutate(class = posclass) %>%

group_by(class) %>%

summarise_all(sd) %>%

ungroup() %>%

t()

colnames(data) <- data[1,]

dataclass <- data[-1,] %>%

apply(2, as.numeric)

colnames(datasd) <- datasd[1,]

dataclasssd <- datasd[-1,] %>%

apply(2, as.numeric)

\# get the ratio

ratio <- dataclass[,-1] / dataclass[,-ncol(dataclass)]

colnames(ratio) <- paste(colnames(dataclass)[-1], colnames(dataclass)[-ncol(dataclass)], sep = "/")

dataratio<- melt(ratio)

fig37 <- ggplot(dataratio,aes(x = value, fill = Var2))+geom_density(alpha=0.382) +

xlab('Ratio on natural logarithm scale') +

scale_x_continuous(trans = log_trans()) +

theme(legend.position="right",legend.text=element_text(size=6))+ labs(fill="")

ggsave(fig37,filename = "fig37.png",width = 6,height = 6)

\# get the rsd within each group

datamean<- melt(dataclass)

datasd <- melt(dataclasssd)

datamean$rsd <- datamean$value/datasd$value*100

\# plot the intensity

fig38 <- ggplot(datamean,aes(x = value, fill=Var2)) +

geom_density(alpha=0.382) +

xlab('Intensity on Log scale') +

scale_x_log10() +

theme(legend.position="right",legend.text=element_text(size=6))+ labs(fill="")

ggsave(fig38,filename = "fig38.png",width = 6,height = 6)

\# plot the rsd

fig39 <- ggplot(datamean,aes(x = rsd, fill=Var2)) +

geom_density(alpha=0.382) +

xlab('RSD%') +

theme(legend.position="right",legend.text=element_text(size=6))+ labs(fill="")

ggsave(fig39,filename = "fig39.png",width = 6,height = 6)

\# plot the intensity

fig40 <- ggplot(datamean,aes(x = value, fill=Var2)) +

geom_density(alpha=0.382) +

xlab('Intensity') +

theme(legend.position="right",legend.text=element_text(size=6))+labs(fill="")

ggsave(fig40,filename = "fig40.png",width = 6,height = 6)

\# plot the distribution

descdist(c(log(datamean$value+1)),discrete = F)

descdist(c(datamean$rsd[!is.na(datamean$rsd)]),discrete = F)

descdist(c(dataratio$value[is.finite(dataratio$value)]),discrete = F)

\## ------------------------------------------------------------------------

\# import data

pos <- read_tsv('data/MTBLS351/m_GCxGC_mass_spectrometry_v2_maf.tsv')

pos2 <- read_tsv("data/MTBLS351/m_UPLC_mass_spectrometry_v2_maf.tsv")

anno <- read_tsv("data/MTBLS351/s_MTBLS351.txt")

\# find the pos data

posdata <- pos %>%

dplyr::select(contains('MH'))

\# find the pos sample class

class <- anno %>%

transmute(class = paste(`Factor Value[Gender]`,`Factor Value[T2D status]`,`Factor Value[Metabolic Syndrome]`,sep = '_')) %>%

as_factor() %>%

c() %>%

unlist()

\# check the order

identical(colnames(posdata),anno$`Sample Name`)

\# get the data mean by group

data <- posdata %>%

t() %>%

as.tibble() %>%

mutate(class = class) %>%

group_by(class) %>%

summarise_all(mean) %>%

ungroup() %>%

t()

\# get the data sd by group

datasd <- posdata %>%

t() %>%

as.tibble() %>%

mutate(class = class) %>%

group_by(class) %>%

summarise_all(sd) %>%

ungroup() %>%

t()

colnames(data) <- data[1,]

dataclass <- data[-1,] %>%

apply(2, as.numeric)

colnames(datasd) <- datasd[1,]

dataclasssd <- datasd[-1,] %>%

apply(2, as.numeric)

\# get the ratio

ratio <- dataclass[,-1] / dataclass[,-ncol(dataclass)]

colnames(ratio) <- paste(colnames(dataclass)[-1], colnames(dataclass)[-ncol(dataclass)], sep = "/")

dataratio<- melt(ratio)

fig41 <- ggplot(dataratio,aes(x = value, fill = Var2))+geom_density(alpha=0.382) +

xlab('Ratio on natural logarithm scale') +

scale_x_continuous(trans = log_trans()) +

theme(legend.position="right",legend.text=element_text(size=6))+ labs(fill="")

ggsave(fig41,filename = "fig41.png",width = 6,height = 6)

\# get the rsd within each group

datamean<- melt(dataclass)

datasd <- melt(dataclasssd)

datamean$rsd <- datamean$value/datasd$value*100

\# plot the intensity

fig42 <- ggplot(datamean,aes(x = value, fill=Var2)) +

geom_density(alpha=0.382) +

xlab('Intensity on Log scale') +

scale_x_log10() +

theme(legend.position="top",legend.text=element_text(size=6))+ labs(fill="")

ggsave(fig42,filename = "fig42.png",width = 6,height = 6)

\# plot the rsd

fig43 <- ggplot(datamean,aes(x = rsd, fill=Var2)) +

geom_density(alpha=0.382) +

xlab('RSD%') +

theme(legend.position="top",legend.text=element_text(size=6))+ labs(fill="")

ggsave(fig43,filename = "fig43.png",width = 6,height = 6)

\# plot the intensity

fig44 <- ggplot(datamean,aes(x = value, fill=Var2)) +

geom_density(alpha=0.382) +

xlab('Intensity') +

theme(legend.position="top",legend.text=element_text(size=6))+labs(fill="")

ggsave(fig44,filename = "fig44.png",width = 6,height = 6)

\# plot the distribution

descdist(c(log(datamean$value+1)),discrete = F)

descdist(c(datamean$rsd),discrete = F)

descdist(c(dataratio$value),discrete = F)

\# find the pos data

posdata <- pos2 %>%

dplyr::select(contains('MH'))

\# check the order

identical(colnames(posdata),anno$`Sample Name`)

\# get the data mean by group

data <- posdata %>%

t() %>%

as.tibble() %>%

mutate(class = class) %>%

group_by(class) %>%

summarise_all(mean) %>%

ungroup() %>%

t()

\# get the data sd by group

datasd <- posdata %>%

t() %>%

as.tibble() %>%

mutate(class = class) %>%

group_by(class) %>%

summarise_all(sd) %>%

ungroup() %>%

t()

colnames(data) <- data[1,]

dataclass <- data[-1,] %>%

apply(2, as.numeric)

colnames(datasd) <- datasd[1,]

dataclasssd <- datasd[-1,] %>%

apply(2, as.numeric)

\# get the ratio

ratio <- dataclass[,-1] / dataclass[,-ncol(dataclass)]

colnames(ratio) <- paste(colnames(dataclass)[-1], colnames(dataclass)[-ncol(dataclass)], sep = "/")

dataratio<- melt(ratio)

fig45 <- ggplot(dataratio,aes(x = value, fill = Var2))+geom_density(alpha=0.382) +

xlab('Ratio on natural logarithm scale') +

scale_x_continuous(trans = log_trans()) +

theme(legend.position="right",legend.text=element_text(size=6))+ labs(fill="")

ggsave(fig45,filename = "fig45.png",width = 6,height = 6)

\# get the rsd within each group

datamean<- melt(dataclass)

datasd <- melt(dataclasssd)

datamean$rsd <- datamean$value/datasd$value*100

\# plot the intensity

fig46 <- ggplot(datamean,aes(x = value, fill=Var2)) +

geom_density(alpha=0.382) +

xlab('Intensity on Log scale') +

scale_x_log10() +

theme(legend.position="top",legend.text=element_text(size=6))+ labs(fill="")

ggsave(fig46,filename = "fig46.png",width = 6,height = 6)

\# plot the rsd

fig47 <- ggplot(datamean,aes(x = rsd, fill=Var2)) +

geom_density(alpha=0.382) +

xlab('RSD%') +

theme(legend.position="top",legend.text=element_text(size=6))+ labs(fill="")

ggsave(fig47,filename = "fig47.png",width = 6,height = 6)

\# plot the intensity

fig48 <- ggplot(datamean,aes(x = value, fill=Var2)) +

geom_density(alpha=0.382) +

xlab('Intensity') +

theme(legend.position="top",legend.text=element_text(size=6))+labs(fill="")

ggsave(fig48,filename = "fig48.png",width = 6,height = 6)

\# plot the distribution

descdist(c(log(datamean$value+1)),discrete = F)

descdist(c(datamean$rsd),discrete = F)

descdist(c(dataratio$value),discrete = F)

\## ------------------------------------------------------------------------

\# import data

pos <- read_tsv('data/MTBLS393/m_mtbls393_metabolite_profiling_mass_spectrometry_v2_maf.tsv')

anno <- read_tsv("data/MTBLS393/s_mtbls393.txt")

\# find the pos data

posdata <- pos[,22:103]

\# find the pos sample class

class <- anno %>%

transmute(class = paste(`Factor Value[Genotype]`,`Factor Value[Inducer conditions]`,sep = '_')) %>%

as_factor() %>%

c() %>%

unlist()

\# check the order

identical(colnames(posdata),anno$`Sample Name`)

\# change the order

posdata <- posdata[,match(anno$`Sample Name`,colnames(posdata))]

identical(colnames(posdata),anno$`Sample Name`)

\# get the data mean by group

data <- posdata %>%

t() %>%

as.tibble() %>%

mutate(class = class) %>%

group_by(class) %>%

summarise_all(mean) %>%

ungroup() %>%

t()

\# get the data sd by group

datasd <- posdata %>%

t() %>%

as.tibble() %>%

mutate(class = class) %>%

group_by(class) %>%

summarise_all(sd) %>%

ungroup() %>%

t()

colnames(data) <- data[1,]

dataclass <- data[-1,] %>%

apply(2, as.numeric)

colnames(datasd) <- datasd[1,]

dataclasssd <- datasd[-1,] %>%

apply(2, as.numeric)

\# get the ratio

ratio <- dataclass[,-1] / dataclass[,-ncol(dataclass)]

colnames(ratio) <- paste(colnames(dataclass)[-1], colnames(dataclass)[-ncol(dataclass)], sep = "/")

dataratio<- melt(ratio)

fig49 <- ggplot(dataratio,aes(x = value, fill = Var2))+geom_density(alpha=0.382) +

xlab('Ratio on natural logarithm scale') +

scale_x_continuous(trans = log_trans()) +

theme(legend.position="right",legend.text=element_text(size=6))+ labs(fill="")

ggsave(fig49,filename = "fig49.png",width = 6,height = 6)

\# get the rsd within each group

datamean<- melt(dataclass)

datasd <- melt(dataclasssd)

datamean$rsd <- datamean$value/datasd$value*100

\# plot the intensity

fig50 <- ggplot(datamean,aes(x = value, fill=Var2)) +

geom_density(alpha=0.382) +

xlab('Intensity on Log scale') +

scale_x_log10() +

theme(legend.position="right",legend.text=element_text(size=6))+ labs(fill="")

ggsave(fig50,filename = "fig50.png",width = 6,height = 6)

\# plot the rsd

fig51 <- ggplot(datamean,aes(x = rsd, fill=Var2)) +

geom_density(alpha=0.382) +

xlab('RSD%') +

theme(legend.position="right",legend.text=element_text(size=6))+ labs(fill="")

ggsave(fig51,filename = "fig51.png",width = 6,height = 6)

\# plot the intensity

fig52 <- ggplot(datamean,aes(x = value, fill=Var2)) +

geom_density(alpha=0.382) +

xlab('Intensity') +

theme(legend.position="right",legend.text=element_text(size=6))+labs(fill="")

ggsave(fig52,filename = "fig52.png",width = 6,height = 6)

\# plot the distribution

descdist(c(log(datamean$value[!is.na(datamean$value)]+1)),discrete = F)

descdist(c(datamean$rsd[!is.na(datamean$rsd)]),discrete = F)

descdist(c(dataratio$value[is.finite(dataratio$value)]),discrete = F)

\## ------------------------------------------------------------------------

library(enviGCMS)

\# dataset 1

library(faahKO)

data("faahko")

\# get the demo data in faahKO packages

cdfpath <- system.file("cdf",package = "faahKO")

list <- getmr(cdfpath, pmethod = ' ')

\# get the data

\# import data

pos <- list$data

anno <- list$group

\# get the data mean by group

data <- pos %>%

t() %>%

as.tibble() %>%

mutate(class = anno) %>%

group_by(class) %>%

summarise_all(mean) %>%

ungroup() %>%

t()

\# get the data sd by group

datasd <- pos %>%

t() %>%

as.tibble() %>%

mutate(class = anno) %>%

group_by(class) %>%

summarise_all(sd) %>%

ungroup() %>%

t()

colnames(data) <- data[1,]

dataclass <- data[-1,] %>%

apply(2, as.numeric)

colnames(datasd) <- datasd[1,]

dataclasssd <- datasd[-1,] %>%

apply(2, as.numeric)

\# get the ratio

ratio <- dataclass[,-1] / dataclass[,-ncol(dataclass)]

names(ratio) <- rep(paste(colnames(dataclass)[-1], colnames(dataclass)[-ncol(dataclass)], sep = "/"),length(ratio))

dataratio<- melt(ratio)

fig53 <- ggplot(dataratio,aes(x = value))+geom_density(alpha=0.382) +

xlab('Ratio on natural logarithm scale') +

scale_x_continuous(trans = log_trans()) +

theme(legend.position="right",legend.text=element_text(size=6))+ labs(fill="")

ggsave(fig53,filename = "fig53.png",width = 6,height = 6)

\# get the rsd within each group

datamean<- melt(dataclass)

datasd <- melt(dataclasssd)

datamean$rsd <- datamean$value/datasd$value*100

\# plot the intensity

fig54 <- ggplot(datamean,aes(x = value, fill=Var2)) +

geom_density(alpha=0.382) +

xlab('Intensity on Log scale') +

scale_x_log10() +

theme(legend.position="right",legend.text=element_text(size=6))+ labs(fill="")

ggsave(fig54,filename = "fig54.png",width = 6,height = 6)

\# plot the rsd

fig55 <- ggplot(datamean,aes(x = rsd, fill=Var2)) +

geom_density(alpha=0.382) +

xlab('RSD%') +

theme(legend.position="right",legend.text=element_text(size=6))+ labs(fill="")

ggsave(fig55,filename = "fig55.png",width = 6,height = 6)

\# plot the intensity

fig56 <- ggplot(datamean,aes(x = value, fill=Var2)) +

geom_density(alpha=0.382) +

xlab('Intensity') +

theme(legend.position="right",legend.text=element_text(size=6))+labs(fill="")

ggsave(fig56,filename = "fig56.png",width = 6,height = 6)

\# plot the distribution

descdist(c(log(datamean$value[!is.na(datamean$value)]+1)),discrete = F)

descdist(c(datamean$rsd[!is.na(datamean$rsd)]),discrete = F)

descdist(c(dataratio$value[is.finite(dataratio$value)]),discrete = F)

\## ----sim-----------------------------------------------------------------

\# PMID: 16762068

\# None

limmafit <- function(data, lv, batch = NULL,log=F){

if(log) data <- log(data+1)

mod <- stats::model.matrix(~lv)

mod0 <- as.matrix(c(rep(1, ncol(data))))

datacor <- signal <- error <- pValues <- qValues <- NULL

if(is.null(batch)){

batch <- NULL

\# limma fit

lmfit <- limma::lmFit(data, mod)

signal <- lmfit$coef[, 1:nlevels(lv)] %*% t(mod[, 1:nlevels(lv)])

error <- data - signal

rownames(signal) <- rownames(error) <- rownames(data)

colnames(signal) <- colnames(error) <- colnames(data)

\# find the peaks with significant differences by F test

\# with BH correction for fdr control without correction

pValues = sva::f.pvalue(data, mod, mod0)

qValues = stats::p.adjust(pValues, method = "BH")

}else{

modcor <- cbind(mod,batch)

modcor0 <- cbind(mod0,batch)

lmfit <- limma::lmFit(data, modcor)

\# data decomposition with batch

batch <- lmfit$coef[, (nlevels(lv) + 1):(nlevels(lv) + NCOL(batch))] %*% t(modcor[, (nlevels(lv) + 1):(nlevels(lv) + NCOL(batch))])

signal <- lmfit$coef[, 1:nlevels(lv)] %*% t(modcor[, 1:nlevels(lv)])

error <- data - signal - batch

datacor <- signal + error

rownames(datacor) <- rownames(batch) <- rownames(signal) <- rownames(error) <- rownames(data)

colnames(datacor) <- colnames(batch) <- colnames(signal) <- colnames(error) <- colnames(data)

\# find the peaks with significant differences by F test

\# with BH correction for fdr control

pValues = sva::f.pvalue(data, modcor, modcor0)

qValues = stats::p.adjust(pValues, method = "BH")

}

\# get the results as list

li <- list(data, datacor, signal, batch, error, pValues, qValues)

names(li) <- c("data","dataCorrected","signal","batch", "error", "p-values", "q-values")

return(li)

}

\# normalize to zero mean and unit variance

AutoScaling <- function(data,lv,log=T){

if(log) data <- log(data+1)

r <- rownames(data)

c <- colnames(data)

data2 <- t(apply(data, 1, function(x) (x - mean(x))/sd(x, na.rm=T)))

rownames(data2) <- r

colnames(data2) <- c

li <- limmafit(data2,lv)

return(li)

}

\# normalize to zero mean and squared root variance

ParetoScaling <- function(data,lv,log=T){

if(log) data <- log(data+1)

r <- rownames(data)

c <- colnames(data)

data2 <- t(apply(data, 1, function(x) (x - mean(x))/sqrt(sd(x, na.rm=T))))

rownames(data2) <- r

colnames(data2) <- c

li <- limmafit(data2,lv)

return(li)

}

\# normalize to zero mean but variance/SE

RangeScaling <- function(data,lv,log=T){

if(log) data <- log(data+1)

r <- rownames(data)

c <- colnames(data)

data2 <- t(apply(data, 1, function(x) (x - mean(x))/(max(x)-min(x))))

rownames(data2) <- r

colnames(data2) <- c

li <- limmafit(data2,lv)

return(li)

}

\# vast scaling

VastScaling <- function(data,lv,log=T){

if(log) data <- log(data+1)

r <- rownames(data)

c <- colnames(data)

data2 <- t(apply(data, 1, function(x) (x - mean(x))/sd(x, na.rm=T) * mean(x)/sd(x, na.rm=T)))

rownames(data2) <- r

colnames(data2) <- c

li <- limmafit(data2,lv)

return(li)

}

\# level scaling

LevelScaling <- function(data,lv,log=T){

if(log) data <- log(data+1)

r <- rownames(data)

c <- colnames(data)

data2 <- t(apply(data, 1, function(x) (x - mean(x))/mean(x)))

rownames(data2) <- r

colnames(data2) <- c

li <- limmafit(data2,lv)

return(li)

}

\# total sum row

TotalSum <- function(data,lv,log=T){

if(log) data <- log(data+1)

r <- rownames(data)

c <- colnames(data)

data2 <- apply(data, 2, function(x) x/sum(x, na.rm=T))

rownames(data2) <- r

colnames(data2) <- c

li <- limmafit(data2,lv)

return(li)

}

\# Median row

MedianNorm <- function(data,lv,log=T){

if(log) data <- log(data+1)

r <- rownames(data)

c <- colnames(data)

data2 <- apply(data, 2, function(x) x/median(x, na.rm=T))

rownames(data2) <- r

colnames(data2) <- c

li <- limmafit(data2,lv)

return(li)

}

\# Mean row

MeanNorm <- function(data,lv,log=T){

if(log) data <- log(data+1)

r <- rownames(data)

c <- colnames(data)

data2 <- apply(data, 2, function(x) x/mean(x, na.rm=T))

rownames(data2) <- r

colnames(data2) <- c

li <- limmafit(data2,lv)

return(li)

}

\# PQN

PQNorm <- function(data,lv,log=T){

if(log) data <- log(data+1)

r <- rownames(data)

c <- colnames(data)

ref <- apply(data[,lv == lv[1]],1,mean)

data2 <- apply(data, 2, function(x) x/median(as.numeric(x/ref), na.rm=T))

rownames(data2) <- r

colnames(data2) <- c

li <- limmafit(data2,lv)

return(li)

}

\# VSN

VSNNorm <- function(data,lv,log=T){

if(log) data <- log(data+1)

fit <- vsn::vsnMatrix(data)

data2 <- fit@hx

li <- limmafit(data2,lv)

return(li)

}

\# Quantile

QuanNorm <- function(data,lv,log=T){

if(log) data <- log(data+1)

data2 <- preprocessCore::normalize.quantiles(data, copy=FALSE)

li <- limmafit(data2,lv)

return(li)

}

\# lumi rsn

LumiRobustSpline <- function(data,lv,log=T){

if(log) data <- log(data+1)

data2 <- lumi::rsn(data)

li <- limmafit(data2,lv)

return(li)

}

\# Limma CyclicLoess

LimmaCyclicLoess <- function(data,lv,log=T){

if(log) data <- log(data+1)

data2 <- limma::normalizeCyclicLoess(data)

li <- limmafit(data2,lv)

return(li)

}

\# AFFA CUBICSpline

LimmaCubicSpline <- function(data,lv,log=T){

if(log) data <- log(data+1)

data2 <- affy::normalize.qspline(data)

li <- limmafit(data2,lv)

return(li)

}

\# SVA

svacor <- function(data,lv,log=T) {

if(log) data <- log(data+1)

mod <- stats::model.matrix(~lv)

svafit <- sva::sva(data, mod)

if (svafit$n.sv == 0) {

message("No surrogate variable found")

li <- limmafit(data,lv)

} else {

message("Data is correcting ...")

batch <- svafit$sv

li <- limmafit(data,lv,batch)

message("Done!")

}

return(li)

}

\# iSVA

isvacor <- function(data, lv,log=T) {

if(log) data <- log(data+1)

isvafit <- try(isva::DoISVA(data, lv, factor.log = T),T)

if(class(isvafit) == 'try-error') {

li <- limmafit(data,lv)

}else{

if (isvafit$nsv == 0) {

message("No surrogate variable found or error for computation.")

li <- limmafit(data,lv)

} else {

message("Data is correcting ...")

batch <- isvafit$isv

li <- limmafit(data,lv,batch)

message("Done!")

}

}

return(li)

}

\# PCR

pcacor <- function(data, lv,log=T) {

if(log) data <- log(data+1)

batch <- svd(data - rowMeans(data))$v[,1]

message("Data is correcting ...")

li <- limmafit(data,lv,batch)

message("Done!")

return(li)

}

\# simulation and compare function

simScenario <- function(name,...){

anstpr <- matrix(nrow = 1000,ncol = 18)

ansfpr <- matrix(nrow = 1000,ncol = 18)

for(i in 1:1000){

sim <- mzrtsim(seed = i,...)

re0 <- limmafit(sim$data,sim$con)

re1 <- AutoScaling(sim$data,sim$con)

re2 <- ParetoScaling(sim$data,sim$con)

re3 <- RangeScaling(sim$data,sim$con)

re4 <- VastScaling(sim$data,sim$con)

re5 <- LevelScaling(sim$data,sim$con)

re6 <- TotalSum(sim$data,sim$con)

re7 <- MedianNorm(sim$data,sim$con)

re8 <- MeanNorm(sim$data,sim$con)

re9 <- PQNorm(sim$data,sim$con)

re10 <- VSNNorm(sim$data,sim$con)

re11 <- QuanNorm(sim$data,sim$con)

re12 <- LumiRobustSpline(sim$data,sim$con)

re13 <- LimmaCyclicLoess(sim$data,sim$con)

re14 <- LimmaCubicSpline(sim$data,sim$con)

re15 <- svacor(sim$data,sim$con)

re16 <- isvacor(sim$data,sim$con)

re17 <- pcacor(sim$data,sim$con)

indexi <- which(re0$`q-values` < 0.05, arr.ind = T)

indexc <- sim$conp

TPR <- FPR <- NULL

TPR[1] <-

length(intersect(indexi, indexc)) / length(indexc)

FPR[1] <-

(length(indexi) - length(intersect(indexi, indexc))) / (nrow(sim$data) - length(indexc))

indexi <- which(re1$`q-values` < 0.05, arr.ind = T)

TPR[2] <-

length(intersect(indexi, indexc)) / length(indexc)

FPR[2] <-

(length(indexi) - length(intersect(indexi, indexc))) / (nrow(sim$data) - length(indexc))

indexi <- which(re2$`q-values` < 0.05, arr.ind = T)

TPR[3] <-

length(intersect(indexi, indexc)) / length(indexc)

FPR[3] <-

(length(indexi) - length(intersect(indexi, indexc))) / (nrow(sim$data) - length(indexc))

indexi <- which(re3$`q-values` < 0.05, arr.ind = T)

TPR[4] <-

length(intersect(indexi, indexc)) / length(indexc)

FPR[4] <-

(length(indexi) - length(intersect(indexi, indexc))) / (nrow(sim$data) - length(indexc))

indexi <- which(re4$`q-values` < 0.05, arr.ind = T)

TPR[5] <-

length(intersect(indexi, indexc)) / length(indexc)

FPR[5] <-

(length(indexi) - length(intersect(indexi, indexc))) / (nrow(sim$data) - length(indexc))

indexi <- which(re5$`q-values` < 0.05, arr.ind = T)

TPR[6] <-

length(intersect(indexi, indexc)) / length(indexc)

FPR[6] <-

(length(indexi) - length(intersect(indexi, indexc))) / (nrow(sim$data) - length(indexc))

indexi <- which(re6$`q-values` < 0.05, arr.ind = T)

TPR[7] <-

length(intersect(indexi, indexc)) / length(indexc)

FPR[7] <-

(length(indexi) - length(intersect(indexi, indexc))) / (nrow(sim$data) - length(indexc))

indexi <- which(re7$`q-values` < 0.05, arr.ind = T)

TPR[8] <-

length(intersect(indexi, indexc)) / length(indexc)

FPR[8] <-

(length(indexi) - length(intersect(indexi, indexc))) / (nrow(sim$data) - length(indexc))

indexi <- which(re8$`q-values` < 0.05, arr.ind = T)

TPR[9] <-

length(intersect(indexi, indexc)) / length(indexc)

FPR[9] <-

(length(indexi) - length(intersect(indexi, indexc))) / (nrow(sim$data) - length(indexc))

indexi <- which(re9$`q-values` < 0.05, arr.ind = T)

TPR[10] <-

length(intersect(indexi, indexc)) / length(indexc)

FPR[10] <-

(length(indexi) - length(intersect(indexi, indexc))) / (nrow(sim$data) - length(indexc))

indexi <- which(re10$`q-values` < 0.05, arr.ind = T)

TPR[11] <-

length(intersect(indexi, indexc)) / length(indexc)

FPR[11] <-

(length(indexi) - length(intersect(indexi, indexc))) / (nrow(sim$data) - length(indexc))

indexi <- which(re11$`q-values` < 0.05, arr.ind = T)

TPR[12] <-

length(intersect(indexi, indexc)) / length(indexc)

FPR[12] <-

(length(indexi) - length(intersect(indexi, indexc))) / (nrow(sim$data) - length(indexc))

indexi <- which(re12$`q-values` < 0.05, arr.ind = T)

TPR[13] <-

length(intersect(indexi, indexc)) / length(indexc)

FPR[13] <-

(length(indexi) - length(intersect(indexi, indexc))) / (nrow(sim$data) - length(indexc))

indexi <- which(re13$`q-values` < 0.05, arr.ind = T)

TPR[14] <-

length(intersect(indexi, indexc)) / length(indexc)

FPR[14] <-

(length(indexi) - length(intersect(indexi, indexc))) / (nrow(sim$data) - length(indexc))

indexi <- which(re14$`q-values` < 0.05, arr.ind = T)

TPR[15] <-

length(intersect(indexi, indexc)) / length(indexc)

FPR[15] <-

(length(indexi) - length(intersect(indexi, indexc))) / (nrow(sim$data) - length(indexc))

indexi <- which(re15$`q-values` < 0.05, arr.ind = T)

TPR[16] <-

length(intersect(indexi, indexc)) / length(indexc)

FPR[16] <-

(length(indexi) - length(intersect(indexi, indexc))) / (nrow(sim$data) - length(indexc))

indexi <- which(re16$`q-values` < 0.05, arr.ind = T)

TPR[17] <-

length(intersect(indexi, indexc)) / length(indexc)

FPR[17] <-

(length(indexi) - length(intersect(indexi, indexc))) / (nrow(sim$data) - length(indexc))

indexi <- which(re17$`q-values` < 0.05, arr.ind = T)

TPR[18] <-

length(intersect(indexi, indexc)) / length(indexc)

FPR[18] <-

(length(indexi) - length(intersect(indexi, indexc))) / (nrow(sim$data) - length(indexc))

anstpr[i,] <- TPR

ansfpr[i,] <- FPR

}

name1 <- paste0('tpr',name,'.csv')

name2 <- paste0('fpr',name,'.csv')

write.csv(anstpr, name1)

write.csv(ansfpr, name1)

}

simScenarioraw <- function(name,...){

anstpr <- matrix(nrow = 1000,ncol = 18)

ansfpr <- matrix(nrow = 1000,ncol = 18)

for(i in 1:1000){

sim <- mzrtsim(seed = i,...)

re0 <- limmafit(sim$data,sim$con,log = F)

re1 <- AutoScaling(sim$data,sim$con,log = F)

re2 <- ParetoScaling(sim$data,sim$con,log = F)

re3 <- RangeScaling(sim$data,sim$con,log = F)

re4 <- VastScaling(sim$data,sim$con,log = F)

re5 <- LevelScaling(sim$data,sim$con,log = F)

re6 <- TotalSum(sim$data,sim$con,log = F)

re7 <- MedianNorm(sim$data,sim$con,log = F)

re8 <- MeanNorm(sim$data,sim$con,log = F)

re9 <- PQNorm(sim$data,sim$con,log = F)

re10 <- VSNNorm(sim$data,sim$con,log = F)

re11 <- QuanNorm(sim$data,sim$con,log = F)

re12 <- LumiRobustSpline(sim$data,sim$con,log = F)

re13 <- LimmaCyclicLoess(sim$data,sim$con,log = F)

re14 <- LimmaCubicSpline(sim$data,sim$con,log = F)

re15 <- svacor(sim$data,sim$con,log = F)

re16 <- isvacor(sim$data,sim$con,log = F)

re17 <- pcacor(sim$data,sim$con,log = F)

indexi <- which(re0$`q-values` < 0.05, arr.ind = T)

indexc <- sim$conp

TPR <- FPR <- NULL

TPR[1] <-

length(intersect(indexi, indexc)) / length(indexc)

FPR[1] <-

(length(indexi) - length(intersect(indexi, indexc))) / (nrow(sim$data) - length(indexc))

indexi <- which(re1$`q-values` < 0.05, arr.ind = T)

TPR[2] <-

length(intersect(indexi, indexc)) / length(indexc)

FPR[2] <-

(length(indexi) - length(intersect(indexi, indexc))) / (nrow(sim$data) - length(indexc))

indexi <- which(re2$`q-values` < 0.05, arr.ind = T)

TPR[3] <-

length(intersect(indexi, indexc)) / length(indexc)

FPR[3] <-

(length(indexi) - length(intersect(indexi, indexc))) / (nrow(sim$data) - length(indexc))

indexi <- which(re3$`q-values` < 0.05, arr.ind = T)

TPR[4] <-

length(intersect(indexi, indexc)) / length(indexc)

FPR[4] <-

(length(indexi) - length(intersect(indexi, indexc))) / (nrow(sim$data) - length(indexc))

indexi <- which(re4$`q-values` < 0.05, arr.ind = T)

TPR[5] <-

length(intersect(indexi, indexc)) / length(indexc)

FPR[5] <-

(length(indexi) - length(intersect(indexi, indexc))) / (nrow(sim$data) - length(indexc))

indexi <- which(re5$`q-values` < 0.05, arr.ind = T)

TPR[6] <-

length(intersect(indexi, indexc)) / length(indexc)

FPR[6] <-

(length(indexi) - length(intersect(indexi, indexc))) / (nrow(sim$data) - length(indexc))

indexi <- which(re6$`q-values` < 0.05, arr.ind = T)

TPR[7] <-

length(intersect(indexi, indexc)) / length(indexc)

FPR[7] <-

(length(indexi) - length(intersect(indexi, indexc))) / (nrow(sim$data) - length(indexc))

indexi <- which(re7$`q-values` < 0.05, arr.ind = T)

TPR[8] <-

length(intersect(indexi, indexc)) / length(indexc)

FPR[8] <-

(length(indexi) - length(intersect(indexi, indexc))) / (nrow(sim$data) - length(indexc))

indexi <- which(re8$`q-values` < 0.05, arr.ind = T)

TPR[9] <-

length(intersect(indexi, indexc)) / length(indexc)

FPR[9] <-

(length(indexi) - length(intersect(indexi, indexc))) / (nrow(sim$data) - length(indexc))

indexi <- which(re9$`q-values` < 0.05, arr.ind = T)

TPR[10] <-

length(intersect(indexi, indexc)) / length(indexc)

FPR[10] <-

(length(indexi) - length(intersect(indexi, indexc))) / (nrow(sim$data) - length(indexc))

indexi <- which(re10$`q-values` < 0.05, arr.ind = T)

TPR[11] <-

length(intersect(indexi, indexc)) / length(indexc)

FPR[11] <-

(length(indexi) - length(intersect(indexi, indexc))) / (nrow(sim$data) - length(indexc))

indexi <- which(re11$`q-values` < 0.05, arr.ind = T)

TPR[12] <-

length(intersect(indexi, indexc)) / length(indexc)

FPR[12] <-

(length(indexi) - length(intersect(indexi, indexc))) / (nrow(sim$data) - length(indexc))

indexi <- which(re12$`q-values` < 0.05, arr.ind = T)

TPR[13] <-

length(intersect(indexi, indexc)) / length(indexc)

FPR[13] <-

(length(indexi) - length(intersect(indexi, indexc))) / (nrow(sim$data) - length(indexc))

indexi <- which(re13$`q-values` < 0.05, arr.ind = T)

TPR[14] <-

length(intersect(indexi, indexc)) / length(indexc)

FPR[14] <-

(length(indexi) - length(intersect(indexi, indexc))) / (nrow(sim$data) - length(indexc))

indexi <- which(re14$`q-values` < 0.05, arr.ind = T)

TPR[15] <-

length(intersect(indexi, indexc)) / length(indexc)

FPR[15] <-

(length(indexi) - length(intersect(indexi, indexc))) / (nrow(sim$data) - length(indexc))

indexi <- which(re15$`q-values` < 0.05, arr.ind = T)

TPR[16] <-

length(intersect(indexi, indexc)) / length(indexc)

FPR[16] <-

(length(indexi) - length(intersect(indexi, indexc))) / (nrow(sim$data) - length(indexc))

indexi <- which(re16$`q-values` < 0.05, arr.ind = T)

TPR[17] <-

length(intersect(indexi, indexc)) / length(indexc)

FPR[17] <-

(length(indexi) - length(intersect(indexi, indexc))) / (nrow(sim$data) - length(indexc))

indexi <- which(re17$`q-values` < 0.05, arr.ind = T)

TPR[18] <-

length(intersect(indexi, indexc)) / length(indexc)

FPR[18] <-

(length(indexi) - length(intersect(indexi, indexc))) / (nrow(sim$data) - length(indexc))

anstpr[i,] <- TPR

ansfpr[i,] <- FPR

}

name1 <- paste0('tprraw',name,'.csv')

name2 <- paste0('fprraw',name,'.csv')

write.csv(anstpr,name)

write.csv(ansfpr,name)

}

\## ----compare-------------------------------------------------------------

\# scenario 1 log

simScenario(name = '1')

\# scenario 2 log

simScenario(name = '2', ncomp = 0.9)

\# scenario 3 log

simScenario(name = '3', ncpeaks = 0.5)

\# scenario 4 log

simScenario(name = '4', nbpeaks = 0.5)

\# scenario 5 log

simScenario(name = '5', ncpeaks = 0.5, nbpeaks = 0.5)

\# scenario 6 log

simScenario(name = '6',batchtype = 'm')

\# scenario 7 log

simScenario(name = '7',batchtype = 'b')

\# scenario 1

simScenarioraw(name = '1')

\# scenario 2

simScenarioraw(name = '2', ncomp = 0.9)

\# scenario 3

simScenarioraw(name = '3', ncpeaks = 0.5)

\# scenario 4

simScenarioraw(name = '4', nbpeaks = 0.5)

\# scenario 5

simScenarioraw(name = '5', ncpeaks = 0.5, nbpeaks = 0.5)

\# scenario 6

simScenarioraw(name = '6',batchtype = 'm')

\# scenario 7

simScenarioraw(name = '7',batchtype = 'b')

\## ------------------------------------------------------------------------

plotbcm <- function(path1,path2,path3,path4, name){

tpr1 <- read_csv(path1, col_types = cols(X1 = col_skip()))

fpr1 <- read_csv(path2, col_types = cols(X1 = col_skip()))

tpr2 <- read_csv(path3, col_types = cols(X1 = col_skip()))

fpr2 <- read_csv(path4, col_types = cols(X1 = col_skip()))

lv <- colnames(fpr2) <- colnames(tpr2) <- colnames(fpr1) <- colnames(tpr1) <- c('None','Auto Scaling','Pareto Scaling','Range Scaling','Vast Scaling','Level Scaling','Total Sum','Median Normalization','Mean Normalization', 'PQN', 'VSN', 'Quantile', 'Robust Spline', 'Cyclic Loess', 'Cubic Spline', 'SVA', 'iSVA', 'PCR')

tpr <- melt(tpr1)

fpr <- melt(fpr1)

tpr2 <- melt(tpr2)

fpr2 <- melt(fpr2)

tpr <- cbind.data.frame(tpr,m='True Positive Rate(log transformed)')

fpr <- cbind.data.frame(fpr,m='False Positive Rate(log transformed)')

tpr2 <- cbind.data.frame(tpr2,m='True Positive Rate(Raw)')

fpr2 <- cbind.data.frame(fpr2,m='False Positive Rate(Raw)')

all <- rbind.data.frame(tpr,fpr,tpr2,fpr2)

\# plot result

ggplot2::ggplot(all,

ggplot2::aes(

x = value,

y = variable,

fill = m

)) + ggridges::geom_density_ridges() + ggplot2::xlim(0, 1) + ggplot2::scale_fill_discrete(name = "Group") +

ggplot2::labs(x = "True Positive Rate / False Positive Rate", y = "Batch correction methods")+ggplot2::ggtitle(name) + ggplot2::theme(legend.position='right',legend.title = ggplot2::element_blank())+

ggplot2::scale_fill_manual(values = c("#D55E0050", "#ff000050","#0072B250","#8080ff50")) +

ggplot2::ggsave(filename = paste0(name,'.png'),width = 8,height = 8)

}

for(i in 1:7){

path1 <- paste0('sim/tpr',i,'.csv')

path2 <- paste0('sim/fpr',i,'.csv')

path3 <- paste0('sim/tprraw',i,'.csv')

path4 <- paste0('sim/fprraw',i,'.csv')

name <- paste('scenario',i)

plotbcm(path1,path2,path3,path4,name = name)

}
